# Disease severity in coinfected hosts: the importance of infection order

**DOI:** 10.1101/2025.03.13.642968

**Authors:** Aloïs Dusuel, Luc Bourbon, Emma Groetz, Nicolas Pernet, Mickaël Rialland, Benjamin Roche, Bruno Faivre, Gabriele Sorci

## Abstract

When hosts are simultaneously infected by different pathogens, the severity of the disease might be altered compared to hosts harboring single infections. The reasons underlying these changes in parasite virulence are manifold. Here, we investigated the importance of order and timing of infection. We used a model of rodent coinfection between two parasites that do not compete for common resources, an intestinal nematode (*Heligmosomoides polygyrus,* Hp) and an apicomplexan protozoan (*Plasmodium yoelii,* Py). During single infections, Hp induced only mild disease symptoms. *Plasmodium* produced a substantial reduction in the number of red blood cells but all mice recovered from the infection. A different picture emerged in coinfected hosts. Hp maintained a profile of mostly asymptomatic infection when infecting hosts that had been previously infected with Py. On the contrary, Py incurred substantially higher costs in hosts that had been previously infected with Hp. We then investigated the possible reasons underlying the increase of Py virulence in hosts that had been previously infected with Hp. We found that coinfected hosts were less able to control Py multiplication and to recover from infection-induced anemia. Coinfected hosts had similar levels of erythropoietin and similar renewal of lost red blood cells compared to single Py infected hosts, resulting in decreased tolerance to Py infection. Experimental administration of erythropoietin in coinfected (Hp infecting first) hosts, partially decreased the severity of disease symptoms and improved tolerance. The detoxification of free heme released during the lysis of red blood cells, and the expression of Th1 and anti-inflammatory cytokine genes were also similar between coinfected and single infected hosts. However, coinfected mice had higher proportions of regulatory T cells expressing the CTLA-4 immune checkpoint, suggesting an enhanced immunosuppressive activity of Tregs. Py infection also induced the exhaustion of CD8^+^ T cells, as coinfected mice had higher proportions of both PD-1^+^ and LAG-3^+^ CD8^+^ T cells, and an increase in the CD4^+^/CD8^+^ ratio. Overall, these results stress the importance of the order of infection as a major determinant of malaria severity in hosts harboring a gastrointestinal nematode infection. We discuss the possible epidemiological and evolutionary consequences of these results.

## Introduction

Coinfection is a widespread phenomenon in plants, wildlife, livestock and humans (Viney and Graham 2013), and describes the occurrence of different parasites/pathogens (hereafter we will use the term parasite as a general term) at the same time in the same individual host. Coinfection has the potential to change the two principal drivers of the infection dynamics, the within host multiplication and the between host transmission (Wait et al. 2021; Imrie et al. 2023), and therefore parasite fitness (Clay and Rudolf 2019) and virulence (i.e., infection-induced damage) (Alizon et al. 2013). During single infections, parasites might have a better access to host resources, while during coinfection, access to resources might be limited if competing parasites share the same infection site or consume the same host resources (Graham 2008; Griffiths et al. 2015). This might affect the within host multiplication rate (for parasites that do multiply within the host) and the transmission rate if coinfection reduces the number of propagules produced per unit of time. This bottom-up regulation of coinfecting parasites might also change the cost paid by the host (e.g., Szabo et al. 2024) if, for instance, the overall host resource consumption exceeds the consumption during single infections. In this case, parasite fitness can also be jeopardized if coinfection compromises the survival of the host. The regulation of the coinfection dynamics might also involve top-down mechanisms, where parasites strive to exploit the host in the face of the immune defenses deployed by the host (Graham 2008). Under this scenario, depending on the nature of the coinfecting parasites, coinfection can have protective [when, for instance, coinfecting parasites elicit similar immune effectors (Filbey et al. 2019)] or antagonistic effects on the host ability to clear the infection (Shen et al. 2019). Specifically, top-down regulation of coinfecting parasites might be compromised when different immune effectors are involved in the process of parasite clearance. Broadly speaking, this is usually the case when hosts are simultaneously infected with microparasites that elicit a Th1 immune response (e.g., viruses and protozoa) and macroparasites that induce a Th2 response (e.g., helminths) (Supali et al. 2010; Ezenwa and Jolles 2011; Ma et al. 2020).

Coinfection between micro and macroparasites is a public health issue in several countries of the intertropical areas, given the high prevalence of helminthiases in humans and the overlap with the endemicity of major infectious diseases, such as malaria or AIDS (Webb et al. 2012; Afolabi et al. 2021). Coinfection between gastrointestinal helminths and malaria has attracted attention in the last decades not only to understand if and how helminths might aggravate the severity of malaria symptoms (Degarege et al. 2016; Kwan et al. 2018), but also to forecast the public health effect of deworming campaigns on malaria incidence and severity (Fenton 2013; Budischak et al. 2018), or to assess the potential effect of helminth infection on the efficacy of malaria vaccines (Su et al. 2006; Noland et al. 2010; Coelho et al. 2019). However, despite considerable effort, epidemiological studies overall report an inconsistent pattern where coinfection can both result in protection or increased severity of malaria symptoms (Nacher 2011; Salazar-Castañón et al. 2014; Degarege and Erko 2016). This inconsistency can be explained by several factors related, for instance, to the species of coinfecting helminths (Adegnika and Kremsner 2012; Hürlimann et al. 2019). Work conducted using animal models has also reported inconsistent results, depending on mouse strains and parasite species (Segura et al. 2009; Knowles 2011; Kolbaum et al. 2012). Here, we suggest that a previously underestimated factor shaping the outcome of the coinfection might also be the order and timing of the infection with the two parasites (Salazar-Castañón et al. 2018; Karvonen et al. 2019; Billet et al. 2020). In most epidemiological studies, the order of infection is not taken into account because this information is typically not available and might contribute to inflate the heterogeneity of the reported results.

The rationale underlying the hypothesis that order of infection is an important driver of the coinfection dynamics rests on the observation that, when coinfecting parasites are regulated through top-down effects, the polarization of the host immune response following the first infection (Hartgers and Yazdanbakhsh 2006; Kotepui et al. 2023) makes the host more susceptible to a subsequent infection with the second parasite (Su et al. 2005; Craig and Scott 2017). We explored this question using a model of coinfection with the gastrointestinal nematode *Heligmosomoides polygyrus* (hereafter Hp) and *Plasmodium yoelii* (hereafter Py) in mice. Hp is a model species in immunoparasitology (Reynolds et al. 2012). This nematode is a natural parasite of rodents (Gregory et al. 1992) which elicits a canonical Th2 immune response, orchestrated by Th2 cytokines, such as IL-4, IL-5 and IL-13 (Reynolds et al. 2012). Mice are infected when ingesting infective larvae (L3), and adult worms live in the lumen of the small intestine where they feed on the host’s intestinal tissue (Bansemir and Sukhdeo 1994). Py is also a model species for the study of the pathogenesis of malaria and the associated immune response (Olatunde et al. 2022). The strain 17XNL of Py produces self-resolving infections, and although the parasitemia can reach high level, all mice recover from the infection in about four weeks. The cycle in the vertebrate host includes an early hepatic-stage that is followed by the asexual replication within the red blood cells (RBCs) (blood-stage); the lysis of RBCs induces anemia, the typical symptom of malaria infection. Infection with Py stimulates the production of pro-inflammatory cytokines, such as the interferon-gamma (IFN-γ), and therefore polarizes the immune response towards Th1 effectors (Olatunde et al. 2022).

Therefore, given that Hp and Py do not share the same ecological niche and do not share common resources for their replication (RBCs for Py), or survival/reproduction (intestinal lumen for Hp), any interference between the two parasites is unlikely to be driven by bottom-up regulation mechanisms. On the contrary, given that the two parasites elicit different immune effectors, and that Th1 and Th2 cytokines have reciprocal inhibitory effects (Paludan 1998), top-down mechanisms are likely candidates to drive the outcome of the coinfection in this system.

We conducted a series of experiments with the aim of exploring to what extent the order and timing of the infection with the two parasites drives the infection dynamics and the severity of the disease. We also explored the possible mechanisms underlying the increased disease severity in coinfected hosts. Based on the rationale reported above, we predicted that if the polarization of the immune response towards Th1 (in the case of Py infection) or Th2 (in the case of Hp infection) makes the host more vulnerable to the infection with the other parasite, coinfection should aggravate the symptoms of the disease, compared to single infections.

## Materials and Methods

### Ethical statement

Experiments were approved by the Comité d’éthique Grand Campus Dijon of the University of Burgundy and were authorized by the French Ministry of Higher Education and Research under the numbers 33492 and 47573.

### Experimental animals and infections

C57BL/6JRj female mice (7-10 weeks old) were purchased from Janvier Labs (Le Genest-Saint-Isle, France), housed in cages containing 5 individuals under pathogen-free conditions, and maintained under a constant temperature of 24°C and a photoperiod of 12h:12h light:dark with *ad libitum* access to water and standard chow diet (A03-10, Safe, Augny, France). All mice were acclimatized to the housing conditions during, at least, 7 days prior to the start of the experiments, were monitored twice a day to check health status, and euthanized by cervical dislocation under anesthesia with isoflurane either if they reached previously defined end points (and considered as mortality events in the survival analysis) or at specific days post-infection (p.i.) for terminal collection of blood and organs (and considered as censored events in the survival analysis).

Mice were infected with *Plasmodium yoelii* 17XNL by intra-peritoneal (i.p.) injection with 5×10^5^ infected red blood cells (iRBC) suspended in 0.1 ml of PBS and/or *Heligmosomoides polygyrus bakeri* by oral gavage with L3 larvae (350 larvae suspended in 0.2 ml of drinking water).

### Experimental groups

Mice were randomly assigned to 8 experimental groups as described in figure 1. A control group (group 1) of mice was sham infected (i.p. injection of 0.1 ml of PBS and oral gavage with a 0.2 ml of drinking water). Two groups (groups 2 and 3) received single infections, either with Hp or Py. Five additional groups were coinfected with the two parasites according to specific orders and timings of infection. In one group, mice were simultaneously (same day) infected with the two parasites (group 4); in two groups, mice were infected first with Hp and subsequently with Py either at day 7 post Hp infection, or at day 28 post Hp infection (groups 5 and 6); in two groups, mice were first infected with Py and subsequently with Hp either at day 14 post Py infection, or at day 28 post Py infection (groups 7 and 8). The timings of infection in the coinfection groups correspond to specific stages in the life cycles of the two parasites. Day 7 and 28 post Hp infection correspond to the stage that just precedes the emergence of the adult worms in the intestinal lumen and when the infection has reached a chronic stage, respectively. Day 14 and day 28 post Py infection approximately correspond to the parasitemia peak and the end of the acute phase infection, respectively. The timings of the sequential coinfection groups also correspond to the timing when the host immune response has been polarized towards a Th2/Treg or a Th1 profile. For the simultaneous infection group, the two types of immune response overlap during the early stage of the infection, with epithelial alarmin cytokines involved in the Th2 response (Pelly et al. 2016, Xing et al. 2024) and NK and γδ T cells producing Th1 cytokines (IFN-γ, TNF-α) (Choudhury et al. 2000). The whole experiment was repeated twice (n_1_ and n_2_) with 40 mice per experimental group (n_1_ = 20 and n_2_ = 20), except for group 6 (n_1_ = 17 and n_2_ = 19) and group 8 (n_1_ = 20 and n_2_ = 15). However, not all traits were measured in all mice of the two replicates.

**Figure 1.**
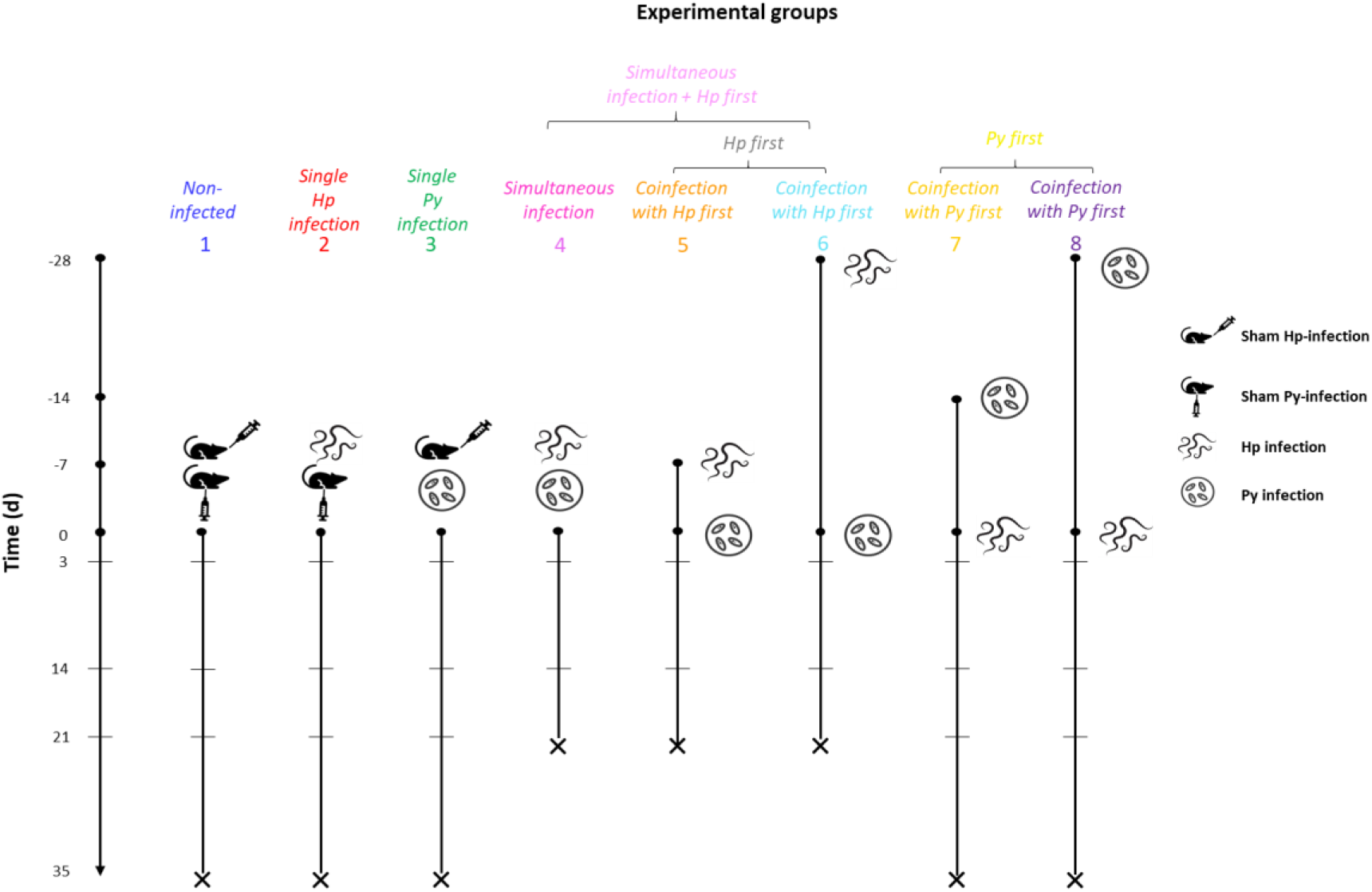
Schematic view of the experimental setup and groups. Non-infected (group 1) and single (Hp, Py) infected mice (groups 2 and 3) were monitored during 35 days post-infection. One group of coinfected mice (group 4) was simultaneously (same day) infected with Py and Hp. The other coinfection groups were either infected first with Hp (at day −7 or day −28) and then with Py at day 0 (groups 5 and 6), or infected first with Py (at day −14 or day −28) and then with Hp at day 0 (groups 7 and 8). Groups 4, 5 and 6 were monitored during 21 days post last exposure. We used a color code to distinguish the different experimental groups: 1 = blue, 2 = red, 3 = green, 4 = magenta, 5 = dark orange, 6 = deep sky blue, 7 = gold, 8 = violet. These color codes were consistently used in all subsequent figures. In addition, certain experimental groups were lumped together in the statistical models: the simultaneous infection plus the groups where hosts were first infected with Hp; the groups where hosts were first infected with Hp; the groups where host were first infected with Py. In this case, a different color code was consistently used: simultaneous infection and Hp first = pink; Hp first = grey; Py first = yellow.

### Erythropoietin administration

Mice (n = 20) were infected with Hp and at day 28 p.i. they were infected with Py. Mice were then split randomly into two groups, one received an i.p. injection of 1 µg of recombinant mouse erythropoietin (rEPO) (959-ME, Bio-Techne, Minneapolis, Minnesota, USA) in 200 µl of PBS at days 0, 2 and 4 post Py infection; the other group was injected with 200 µl of PBS only, for control. Erythropoietin is the hormone that regulates the production of RBCs and this experiment served to investigate if rEPO administration buffered to severity of the disease symptoms.

### Monitoring of health status, hematological and plasma parameters, and Py parasitemia

Mice were regularly weighed (at least three times a week) and body mass was used as an integrative marker of host health. Hematological parameters were assessed by collecting 10 µl of blood from the tail tip in 0.5 ml tubes with 1 µl of heparin (Héparine Choay® 25 000 UI/5ml, Sanofi-Aventis, Gentilly, France). These samples were immediately read with a SCIL Vet abc Plus+ hematology analyzer (Horiba Medical, Montpellier, France) to assess RBC counts and mean globular volume (MGV). RBC counts inform us on the possible anemia induced by the infection, whereas MGV is a proxy of reticulocyte production and inform us on the replacement of RBCs lost during the infection. For mice with Py infection, a drop of blood was smeared on a slide and fixed with methanol. Slides were subsequently stained with a Giemsa RS solution (Carlo Erba Reagents, Val de Reuil, France) (10% v/v in phosphate buffer) for 15 minutes and rinsed with water. Infected RBCs were counted using an optical microscope under magnification x1000, and parasitemia expressed as the proportion of iRBCs.

At day 3, 14, 21 and 35 p.i. (post last infection for the coinfection groups), five mice per group were euthanized by cervical dislocation under isoflurane anesthesia, for terminal collection of total peripheral blood and organs. Given the severity of the disease symptoms, the simultaneous infection group and the two coinfection groups first infected with Hp (groups 4, 5, 6) were only monitored up to day 21 post last infection. Plasma was separated from total blood by centrifugation (7 min 3000 rpm 4°C), aliquoted and stored at −80°C.

Plasma levels of aspartate aminotransferase (AST) and heme oxygenase-1 (HO-1) were assessed with ab263882 and ab229431 commercial ELISA kits from Abcam (Cambridge, UK), respectively, with the aim of investigating infection-induced liver damage (AST) and the scavenging of free heme groups released during the lysis of RBCs (HO-1). Plasma levels of mouse erythropoietin (EPO) were assessed with EM28RB commercial ELISA kit from Invitrogen™ (Carlsbad, California, USA), with the aim of assessing the production of the hormone that regulates the production of RBCs. Plates were read with a SpectraMax iD3 multi-mode microplate reader (Molecular Devices, San Jose, California, USA).

### Organ collection

Spleen was removed and weighed because splenomegaly (i.e., the enlargement of the spleen) is a prominent symptom of the infection. About one third of the spleen was cut and immediately frozen in liquid nitrogen for RT-qPCR, and the remaining was kept on ice in PBS before the flow cytometry staining procedure. A liver slice (left lobe) was also immediately frozen in liquid nitrogen for RT-qPCR.

### RNA extraction and qPCR for Hmox-1, IFN-γ, IL-10 and TGF-β1 gene expression

Tissues were homogenized in TRIzol™ Reagent (Invitrogen™, Carlsbad, California, USA) under strong agitation using 0.5 mm glass beads and a Precellys® 24 Touch homogenizer (Bertin Technologies, Montigny-le-Bretonneux, France). RNA extraction was performed following the manufacturer’s instructions. RNA concentration was measured with a NanoPhotometer® N50 (Implen, Munich, Germany). Reverse transcription was performed with a High-Capacity cDNA Reverse Transcription Kit (Applied Biosystems™, Foster City, California, USA) from 1.5 µg total RNA. Quantitative PCR was performed with PowerUp™ SYBR™ Green Master Mix (Applied Biosystems™) on a QuantStudio™ 3 Real-Time PCR System (Applied Biosystems™). We used two housekeeping genes (β-actin and GAPDH); β-actin provided more consistent values (less intra-group variability) and therefore we used it as the reference gene (i.e., the level of expression of the genes of interest relative to the expression of the housekeeping gene). Primer sequences are reported in the supplementary material (table S1), and were checked for specificity using the Primer-BLAST tool (NCBI). Melt curve analysis was also performed in all samples to check for the presence of nonspecific amplification products.

### Assessment of immune cell populations through spectral flow cytometry

Spleens were homogenized with a 70 µm cell strainer and washed twice with PBS to obtain a single-cell suspension. RBCs were removed from suspensions with eBioscience™ RBC Lysis Buffer (Invitrogen™) (3 min incubation at RT). Cells were counted with viability trypan blue dye and dispatched in 96-well V bottom plates to obtain 1×10^7^ live cells/well. Cells were then stained with LIVE/DEAD™ Fixable Blue Dead Cell Stain Kit (Invitrogen™) in PBS for 30 min at 4°C, incubated in stain buffer with BD Fc Block™ (BD Pharmingen™, Franklin Lakes, New Jersey, USA) for 15 min at RT (0.5 µg/well), and stained with surface antibodies in brilliant stain buffer (see table S2 for details on the markers used) for 30 min at 4°C. Cells were then fixed and permeabilized with eBioscience™ Foxp3 / Transcription Factor Staining Buffer Set (Invitrogen™) prior intracellular staining in brilliant stain buffer (table S2) for 30 min at RT. Cells were resuspended in stain buffer, filtered with 100 µm nylon mesh and analyzed on a 4-lasers (UV/V/B/R) Aurora™ (Cytek® Biosciences, Fremont, California, USA) spectral flow cytometer in the ImaFlow Facility, part of the US58 BioSanD. Events were gated and analyzed with SpectroFlo® v3.0.0 software (Cytek® Biosciences) (see figures S1 and S2 for the gating strategy).

### Statistical analyses

We used general and generalized linear mixed models (GLMM) to analyze data where each individual mouse was repeatedly sampled, using either a normal distribution or a beta distribution (i.e., for proportions) of errors. Mouse ID was included in these models as a random intercept. Variables do not necessarily vary linearly with time over the course of the infection. We therefore included squared (and cubic) time p.i. to capture these nonlinear relationships. However, polynomial regression might provide a poor fit to the data. Therefore, we also ran general additive models (GAM) where the degrees of freedom of the smoothing function were chosen based on the generalized cross-validation (GCV) optimization criterion. Some variables had an asymmetrical distribution and were log-transformed to reduce skewness and meet the model assumptions.

Differences in survival between groups were analyzed using the Kaplan Meier estimator and significance assessed using a Log-Rank test. For coinfected mice, any mortality that had occurred before the infection with the second parasite was considered as caused by single infections (and therefore included as single infected individuals in the model); any effect of coinfection on survival was, thus, assessed only after the infection with the second parasite. To increase the statistical power, the survival analysis also included additional mice that were followed for a longer period of time (up to day 99 p.i.) to assess the long-term effect of coinfection on the infection dynamics. These mice, that involved all the infection groups (excluding therefore the non-infected individuals), were housed in the same conditions and received the same infection protocols (dose, timing of the infection), the only difference being that they were not euthanized at specific time points (unless they reached the end points). Therefore, the survival analysis included up to four replicated datasets for the infection groups and two replicated datasets for the non-infected group.

For variables that were measured only once per individual mouse (spleen mass, AST, EPO, gene expressions, HO-1, and immune cell population by flow cytometry), we ran general or generalized linear models (GLM), depending on the distribution of errors (normal or beta distribution). Post-hoc comparisons between groups were done using LS-means with Bonferroni correction of p values.

Differences in sample size across groups reflect missing values, either due to the reaching of the end point or to technical issues during the processing of the sample.

To simplify the interpretation of the results, we ran different sets of models, comparing different groups. First, we compared non-infected mice versus single infections (groups 1, 2 and 3). These models provided us with the baseline information on the intrinsic effect of each parasite when infecting the host alone. Then, we compared single infections versus coinfection groups. This was done separately for Py and Hp by including in the model the corresponding coinfection groups where Py and Hp infected mice that had been previously infected with the other parasite (i.e., groups 3, 4, 5, 6 in one model, and groups 2, 7, 8 in another model). The simultaneous infection group (group 4) was always included in the Py model. Given that the timing of the infection did appear to have minor effects on the response variables, we also ran models where the different coinfection groups were clustered together (groups 4, 5 and 6 together; groups 7 and 8 together) and we compared non-infected, single infected and coinfected hosts in the same model. Finally, we ran a model that was restricted to the sequential coinfection groups (excluding the simultaneous infection group) where we could assess the effect of the order of the infection [either Hp first (groups 5 and 6), or Py first (groups 7 and 8)]. All the statistical analyses and figures were done using SAS Studio.

## Results

### Severity of the disease in single and coinfected hosts

#### Body mass

We analyzed body mass as percent change relative to the body mass at the day of the sham-infection for the control group (group 1) and the day of the infection for single and simultaneous infection groups (groups 2, 3 and 4) or the day of the last infection for the sequential coinfection groups (groups 5, 6, 7 and 8) (corresponding to t = 0 in fig. 1). We first compared changes in body mass over time between non-infected and single (Hp, Py) infected mice (groups 1, 2, 3). The model showed that body mass increased linearly over time, at a similar rate for non-infected and single infected mice (table 1, fig. 2a).

**Figure 2.**
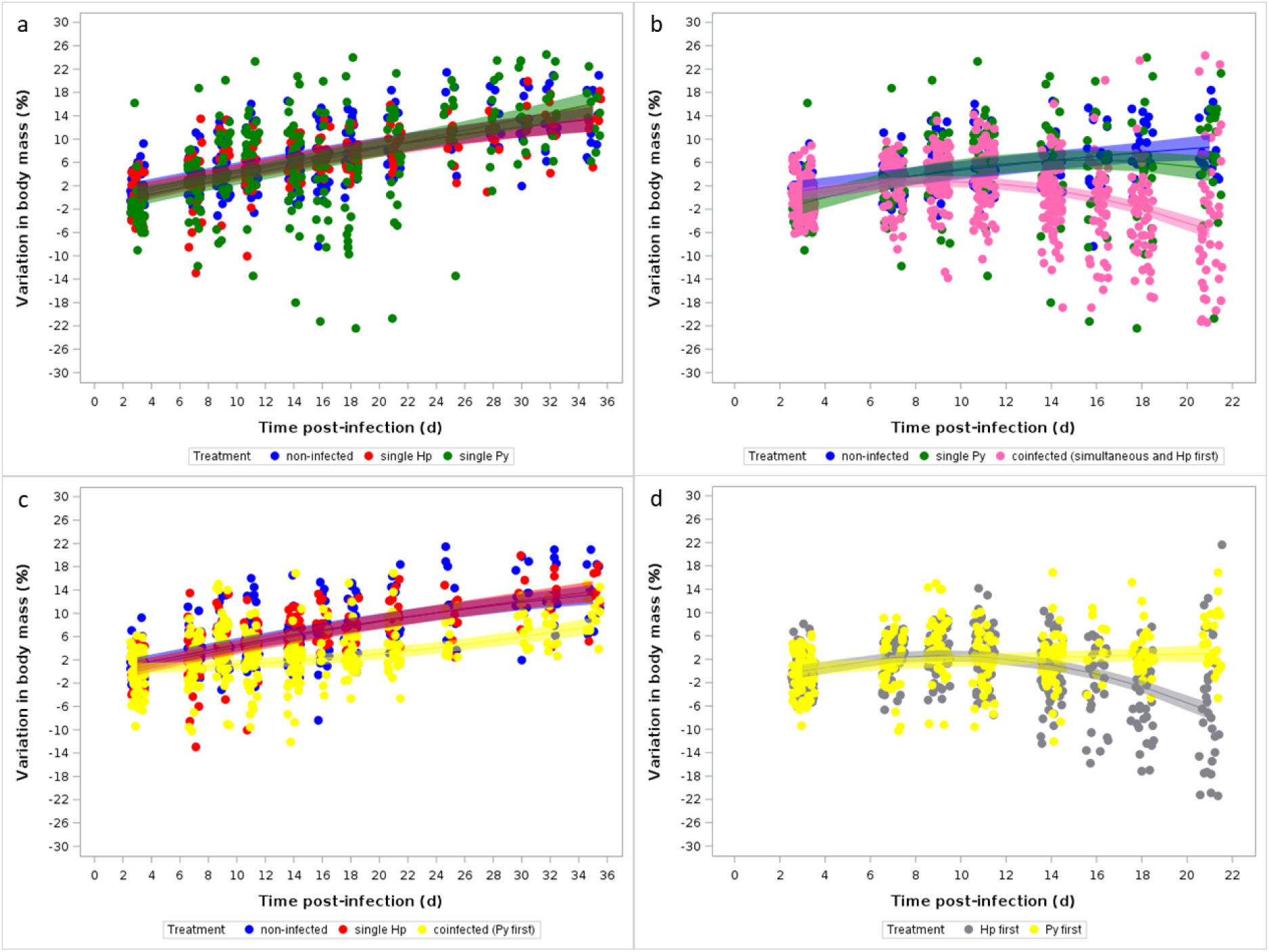
Changes in body mass over the course of the infection. Body mass is expressed as percent change with respect to body mass at day 0 [for the non-infected, single (Hp, Py) infected, and simultaneous infection groups] or at the day of the last infection for the sequential coinfection groups. a) Changes in body mass for non-infected and single (Hp, Py) infected mice; b) Changes in body mass for non-infected, single Py infected and coinfected (simultaneous infection and Hp infecting first) groups; c) Changes in body mass for non-infected, single Hp infected and coinfected (Py infecting first) groups; d) Changes in body mass for coinfected groups depending on the order of the infection (Hp infecting first or Py infecting first). Dots represent the raw data, the lines represent the fit of a GLMM, and the shaded area around the line the 95% CI.

**Table 1.** General linear mixed model investigating the changes in body mass (percent change with respect to day 0) in non-infected and single (Hp and Py) infected mice. We report the parameter estimates with the 95% CI, degrees of freedom (df), F and p values. Mouse ID was included as a random effect to take into account the non-independence of observations for the same individual over time. N = 120 individuals and 774 observations.

| Fixed effects |  | Estimate | 95% CI | df | F | p |
| --- | --- | --- | --- | --- | --- | --- |
| Treatment |  |  |  | 2,648 | 0.24 | 0.7828 |
|  | non-infected | 0 |  |  |  |  |
|  | single Hp | -0.47 | -3.13/2.20 |  |  |  |
|  | single Py | -0.95 | -3.60/1.71 |  |  |  |
| Time |  | 0.54 | 0.35/0.74 | 1,648 | 83.44 | <0.0001 |
| Squared time |  | -0.004 | -0.01/0.001 | 1,648 | 2.80 | 0.095 |
| Time*Treatment |  |  |  | 2,648 | 0.12 | 0.8827 |
|  | non-infected | 0 |  |  |  |  |
|  | single Hp | 0.007 | -0.27/0.29 |  |  |  |
|  | single Py | -0.06 | -0.33/0.22 |  |  |  |
| Squared Time*Treatment |  |  |  | 2,648 | 0.88 | 0.4141 |
|  | non-infected | 0 |  |  |  |  |
|  | single Hp | 0.0003 | -0.01/0.01 |  |  |  |
|  | single Py | 0.005 | -0.003/0.01 |  |  |  |
| Random effect |  | Estimate | SE | z | P |  |
| ID |  | 16.49 | 2.49 | 6.62 | <0.0001 |  |

We then compared changes in body mass in mice that were infected with Py alone and mice that were simultaneously infected or were first infected with Hp and then with Py (groups 4, 5, 6). Although mice in the coinfection groups lost body mass from day 14 post Py infection onwards (table S3, fig. S3a), the squared time by treatment interaction was not statistically significant (table S3). Given that there was no apparent difference in the dynamics of body mass change between the coinfection groups, we ran an additional model where we compared the changes in body mass between non-infected, single Py infected and coinfected groups (groups 4, 5 and 6 together). This model showed a strong effect of coinfection on changes in body mass (table 2, fig. 2b), with coinfected mice losing body mass over the course of the infection.

**Table 2.**
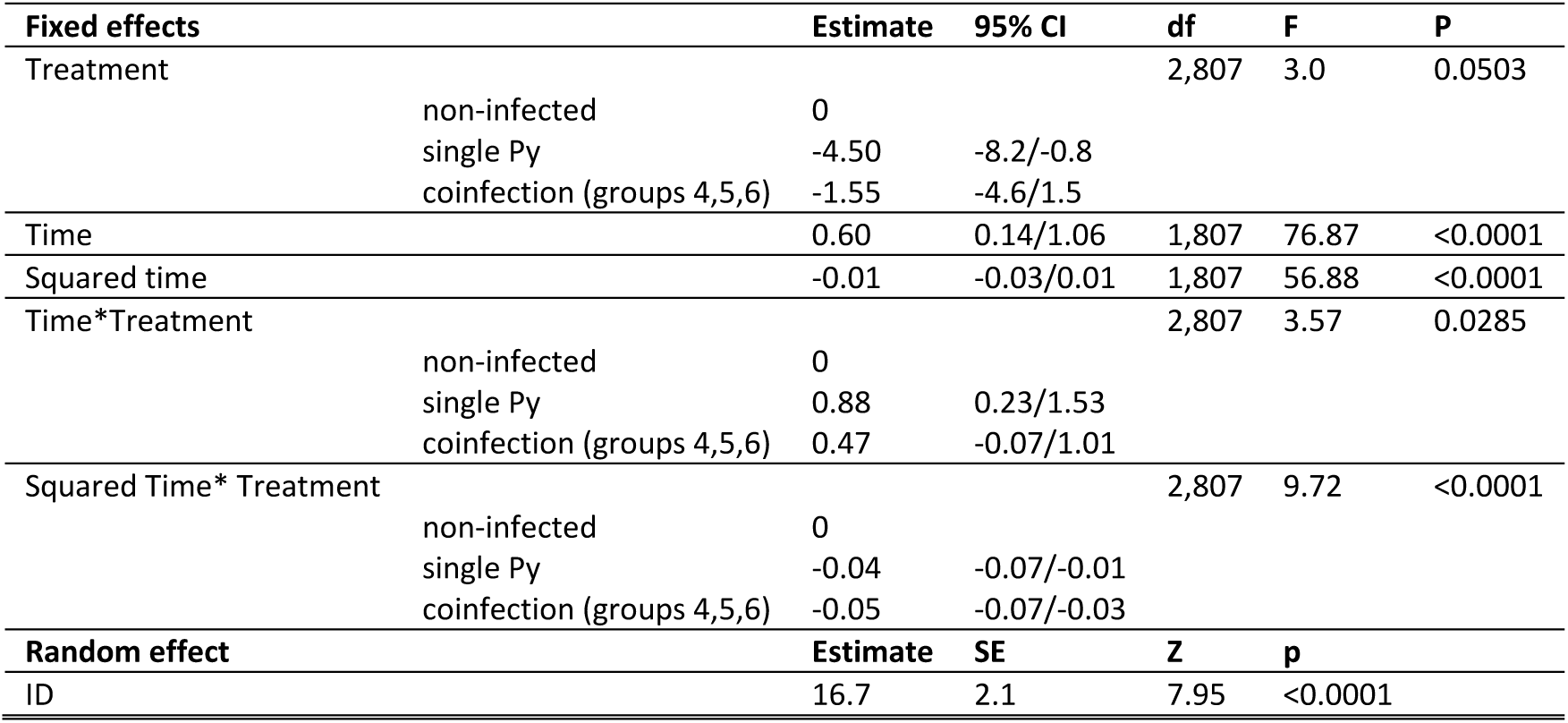
General linear mixed model investigating the changes in body mass (percent change with respect to day 0) in non-infected, single Py infected and coinfected (simultaneous infection and Hp infecting first) mice. We report the parameter estimates with the 95% CI, degrees of freedom (df), F and p values. Mouse ID was included as a random effect to take into account the non-independence of observations for the same individual over time. N = 196 individuals and 1009 observations.

| Fixed effects |  | Estimate | 95% CI | df | F | P |
| --- | --- | --- | --- | --- | --- | --- |
| Treatment |  |  |  | 2,807 | 3.0 | 0.0503 |
|  | non-infected | 0 |  |  |  |  |
|  | single Py | -4.50 | -8.2/-0.8 |  |  |  |
|  | coinfection (groups 4,5,6) | -1.55 | -4.6/1.5 |  |  |  |
| Time |  | 0.60 | 0.14/1.06 | 1,807 | 76.87 | <0.0001 |
| Squared time |  | -0.01 | -0.03/0.01 | 1,807 | 56.88 | <0.0001 |
| Time*Treatment |  |  |  | 2,807 | 3.57 | 0.0285 |
|  | non-infected | 0 |  |  |  |  |
|  | single Py | 0.88 | 0.23/1.53 |  |  |  |
|  | coinfection (groups 4,5,6) | 0.47 | -0.07/1.01 |  |  |  |
| Squared Time* Treatment |  |  |  | 2,807 | 9.72 | <0.0001 |
|  | non-infected | 0 |  |  |  |  |
|  | single Py | -0.04 | -0.07/-0.01 |  |  |  |
|  | coinfection (groups 4,5,6) | -0.05 | -0.07/-0.03 |  |  |  |
| Random effect |  | Estimate | SE | Z | p |  |
| ID |  | 16.7 | 2.1 | 7.95 | <0.0001 |  |

A different picture emerged when comparing changes in body mass in mice that were infected first with Py and then with Hp (groups 7 and 8). In this case, coinfection slowed down the rate of body mass gain, but did not produce a loss in body mass (table S4, fig. S3b). As the timing of infection did not have any effect on the changes in body mass, we grouped them and ran an overall model comparing non-infected, single Hp infected and coinfected groups (7 and 8 together). This model confirmed the slight effect of coinfection on the dynamics of body mass change, with coinfected mice having a slower rate of increase in body mass, but not losing mass over time (table 3, fig. 2c). Finally, we ran a model that was only restricted to the four sequential coinfection groups (groups 5, 6, 7 and 8), excluding the simultaneous infection group. This model confirmed to importance of the order of infection in shaping the dynamics of body mass (table S5, fig. 2d).

**Table 3.** General linear mixed model investigating the changes in body mass (percent change with respect to day 0) in non-infected, single Hp infected and coinfected (Py infecting first) mice. We report the parameter estimates with the 95% CI, degreed of freedom (df), F and p values. Mouse ID was included as a random effect to take into account the non-independence of observations for the same individual over time. N = 155 individuals and 869 observations.

| Fixed effects |  | Estimate | 95% CI | df | F | p |
| --- | --- | --- | --- | --- | --- | --- |
| Treatment |  |  |  | 2,708 | 1.06 | 0.3477 |
|  | non-infected | 0 |  |  |  |  |
|  | single Hp | -0.52 | -2.61/1.57 |  |  |  |
|  | coinfection (groups 7,8) | 0.8 | -1.042.64 |  |  |  |
| Time |  | 0.54 | 0.40/0.68 | 1,708 | 86.46 | <0.0001 |
| Squared time |  | -0.004 | -0.008/-0.0005 | 1,708 | 0.58 | 0.4468 |
| Time* Treatment |  |  |  | 2,708 | 22.19 | <0.0001 |
|  | non-infected | 0 |  |  |  |  |
|  | single Hp | 0.007 | -0.2/0.21 |  |  |  |
|  | coinfection (groups 7,8) | -0.52 | -0.70/-0.34 |  |  |  |
| Squared Time* Treatment |  |  |  | 2,708 | 9.50 | <0.0001 |
|  | non-infected | 0 |  |  |  |  |
|  | single Hp | 0.0007 | -0.005/0.006 |  |  |  |
|  | coinfection (groups 7,8) | 0.01 | 0.005/0.015 |  |  |  |
| **Random effect** |  | **Estimate** | **SE** | **z** | **P** |  |
| ID |  | 11.96 | 1.62 | 7.37 | <0.0001 |  |

Therefore, coinfection aggravated one of the most integrative symptoms of disease severity, but the magnitude of the aggravating effect was strongly dependent on the order of the infection, being greatest in mice that were infected with Hp first.

#### Splenomegaly

We measured spleen mass at day 3, 14, 21 and 35 p.i. and confirmed that single infection induced splenomegaly of different magnitude depending on the parasite, with larger spleen mass in Py infected mice, especially at day 14 – 21 p.i. (table S6, fig. S4a). When comparing groups infected first with Hp and then with Py (plus the simultaneous infection, groups 4, 5, 6), coinfection only had a mild initial effect on splenomegaly at day 3 post Py infection compared to single Py infected hosts (Table S7), but by day 14 p.i. the differences vanished (fig. S5a). This result was corroborated when comparing non-infected, single Py infected and coinfected groups together (Table S8, fig. S4b).

When comparing the effect of coinfection in groups that were first infected by Py (groups 7 and 8), we found only a moderate increase in spleen mass, essentially as a consequence of the residual enlargement due to the previous Py infection, since the difference between single Hp infection and coinfection groups tended to vanish over time (table S9 and table S10, fig. S4c, fig. S5b).

The asymmetric effect of infection order on splenomegaly was best captured by the model on the four sequential coinfection groups (groups 5, 6, 7 and 8) where mice were either infected first with Hp or with Py. This model showed that spleen mass increased over the course of the infection only when mice were first infected with Hp (table S11, fig. S4d).

Therefore, as for body mass, coinfection affected splenomegaly in an asymmetric way, depending on infection order, with larger spleens in mice infected first with Hp and subsequently with Py.

#### Marker of liver damage

We compared a marker of liver damage (aspartate aminotransferase, AST) in plasma at day 21 p.i. in non-infected and single (Hp, Py) infected mice, and found that Py infected mice had moderately higher levels of AST compared to non-infected hosts (GLM, F_2,23_ = 4.45, p = 0.0218, n = 26 individuals; fig. S6a). Coinfection did aggravate liver damage in the groups where Py followed the Hp infection (plus simultaneous infection, groups 4, 5, 6) (GLM, F_3,36_ = 4.28, p = 0.0111, n = 40; fig. S7a), and this effect was confirmed when analyzing non-infected, single Py infected and coinfected mice together in the same model (GLM, F_2,45_ = 4.22, p = 0.0209, n = 48; fig. S6b).

We found rather similar effects of coinfection when analyzing the groups where Hp follows Py (groups 7 and 8), with coinfection groups having higher AST levels compared to non-infected and single Hp infected groups (GLM, F_2,29_ = 4.96, p = 0.0141, n = 32; fig. S6c, fig. S7b). As a consequence, there was no effect of infection order in the coinfection groups (groups 5, 6, 7 and 8) on AST level at day 21 p.i. (GLM, F_1,35_ = 0.28, p = 0.5976, n = 37; fig. S6d).

Therefore, coinfection tended to aggravate liver damage with no effect of infection order.

#### Survival

In agreement with the pattern of change in body mass over the course of the infection, we did not find any difference in the survival rate between non-infected and single infected mice (Log-Rank test χ²_2_ = 0.92, p = 0.6315, n = 229; fig. 3a). Single Py infected mice had higher survival compared to the simultaneous infection and coinfection groups (groups 4, 5 and 6) (Log-Rank test χ²_3_ = 22.81, p < 0.0001, n = 242, fig. S8), but there was no difference according to the timing of coinfection (all sidak-adjusted p values > 0.6. We therefore compared the coinfection groups together with the non-infected and single Py infected and found a strong effect of coinfection (Log-Rank test χ²_2_ = 26.02, p < 0.0001, n = 282; fig. 3b).

**Figure 3.**
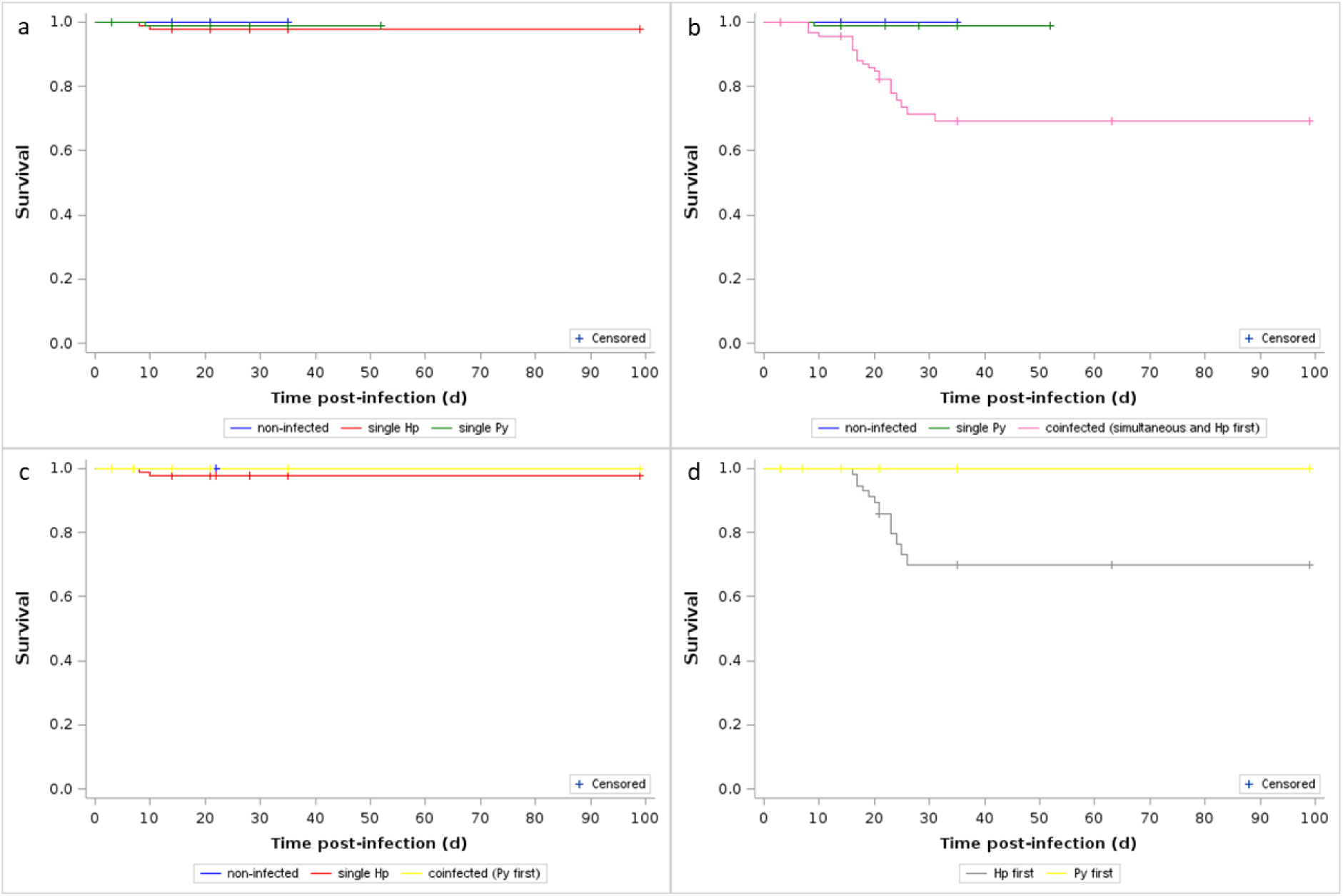
Effect of infection and coinfection on host survival. a) Survival curves for non-infected and single (Hp, Py) infected mice; b) Survival curves for non-infected, single Py infected and coinfected (simultaneous infection and Hp first) mice; c) Survival curves for non-infected, single Hp infected and coinfected (Py infecting first) mice; d) Survival curves for coinfected mice depending on the order of infection (Hp infecting first or Py infecting first). Crosses indicate censored observations.

There was no host mortality in the coinfection groups where Hp followed Py (groups 7 and 8). Therefore, there was no difference in the mortality rate of non-infected, single Hp infected and coinfected groups (groups 7 and 8) (Log-Rank test χ²_2_ = 2.34, p = 0.3099, n = 229; fig. 3c). As a consequence, order of infection was a strong predictor of host mortality in the sequential coinfection groups (groups 5, 6, 7 and 8) (Log-Rank test χ²_1_ = 13.47, p = 0.0002, n = 189; fig. 3d).

Overall, these results show that order of infection has a paramount effect on the severity of the symptoms and therefore on parasite virulence (infection-induced damage), with Hp infection producing mild symptoms whatever the order of the infection, and Py becoming substantially more virulent when infecting hosts already harboring the nematode. However, the timing of infection had virtually no effect on disease severity.

### Parasitemia

If previous infection with Hp makes the host more susceptible to a subsequent Py infection, we should expect higher parasitemia in coinfected hosts (groups 4, 5, 6) compared to single Py infection. During single Py infection, parasitemia reached a peak at around day 16 p.i., and then the infection resolved by day 30 (fig. 4a; fig. S9). Given that the coinfection groups (groups 4, 5 and 6) were stopped at day 21 p.i., we could only compare the rising and the early declining stages of the parasitemia. The model showed that coinfected groups reached higher parasitemia and that, contrary to the single Py infection group, the parasitemia of coinfected mice kept increasing up to day 21 p.i. (table S12 and table 4, fig. S10 and fig. 4b).

**Figure 4.**
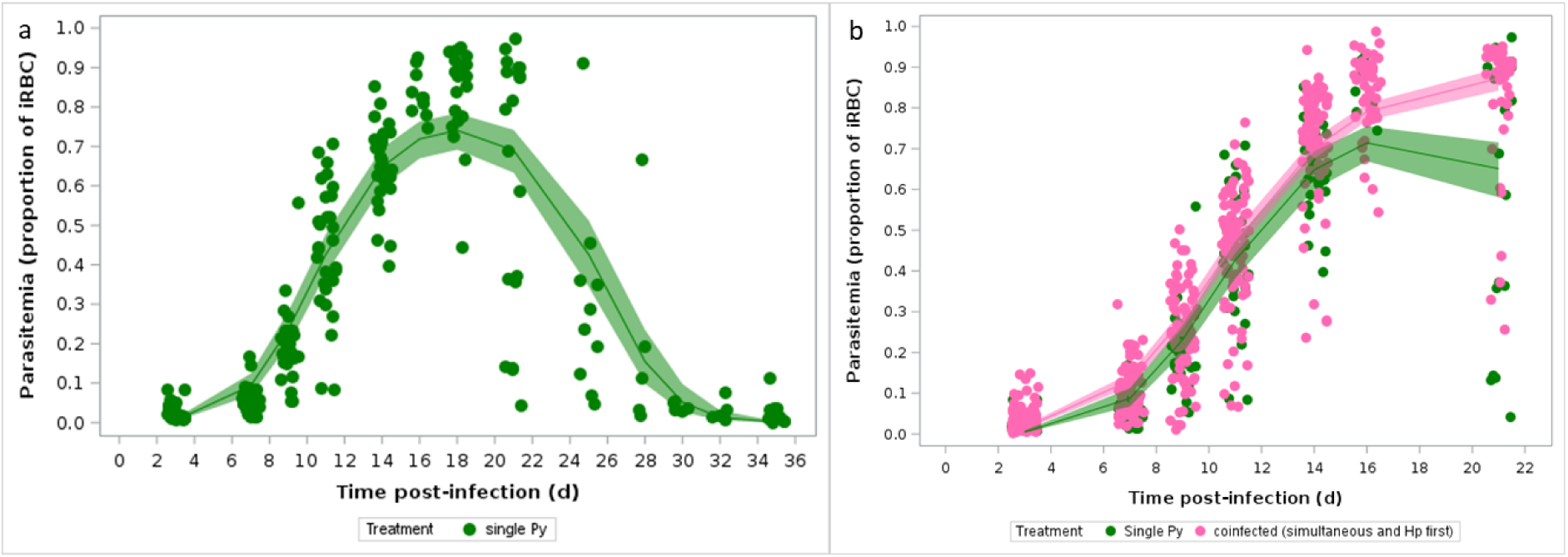
Changes in parasitemia over the course of the infection. a) Changes in parasitemia (proportion of iRBCs counted using optical microscopy) in single Py infected mice over time; b) Changes in parasitemia (proportion of iRBCs) in single Py infected and coinfected (simultaneous infection and Hp first) mice. Dots represent the raw data, the lines represent the fit of the GLMM, and the shaded area around the line the 95% CI.

**Table 4.**
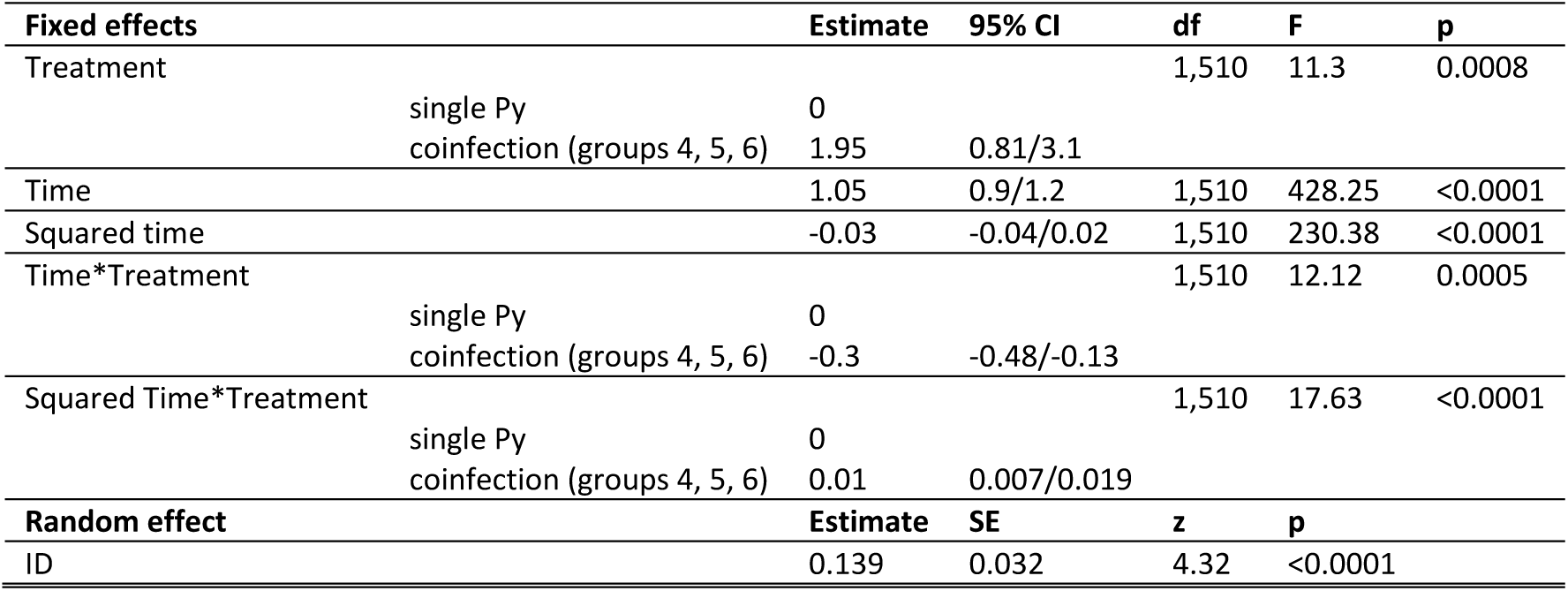
Generalized linear mixed model investigating the changes in parasitemia (proportion of iRBC) in single Py infected and coinfected (simultaneous infection and Hp first) hosts. We report the parameter estimates with the 95% CI, degrees of freedom (df), F and p values. Mouse ID was included as a random effect to take into account the non-independence of observations for the same individual over time. The distribution of errors was modelled using a beta distribution. N = 151 individuals and 665 observations.

Therefore, as expected, previous or simultaneous infection with Hp increased host susceptibility to a subsequent infection, since Py replicated more and for longer time in coinfected mice. Enhanced persistent parasitemia might thus explain the observed increase of Py virulence in coinfected hosts through specific costs incurred during the asexual reproduction of Py.

### Specific costs of Py infection

The principal cost induced by Py infection is related to the anemia caused by the lysis of RBCs during the asexual reproduction of the parasite (Olatunde et al. 2022). We therefore assessed the changes in RBC counts over the course of the infection. First, we compared changes in RBC counts between non-infected and single (Hp, Py) infections. As expected, Py infection produced a sharp decline in the number of RBCs, while there was no difference between non-infected and single Hp infection (table 5, fig. 5a, fig. S11), corroborating the idea that Py and Hp do not compete for common resources.

**Figure 5.**
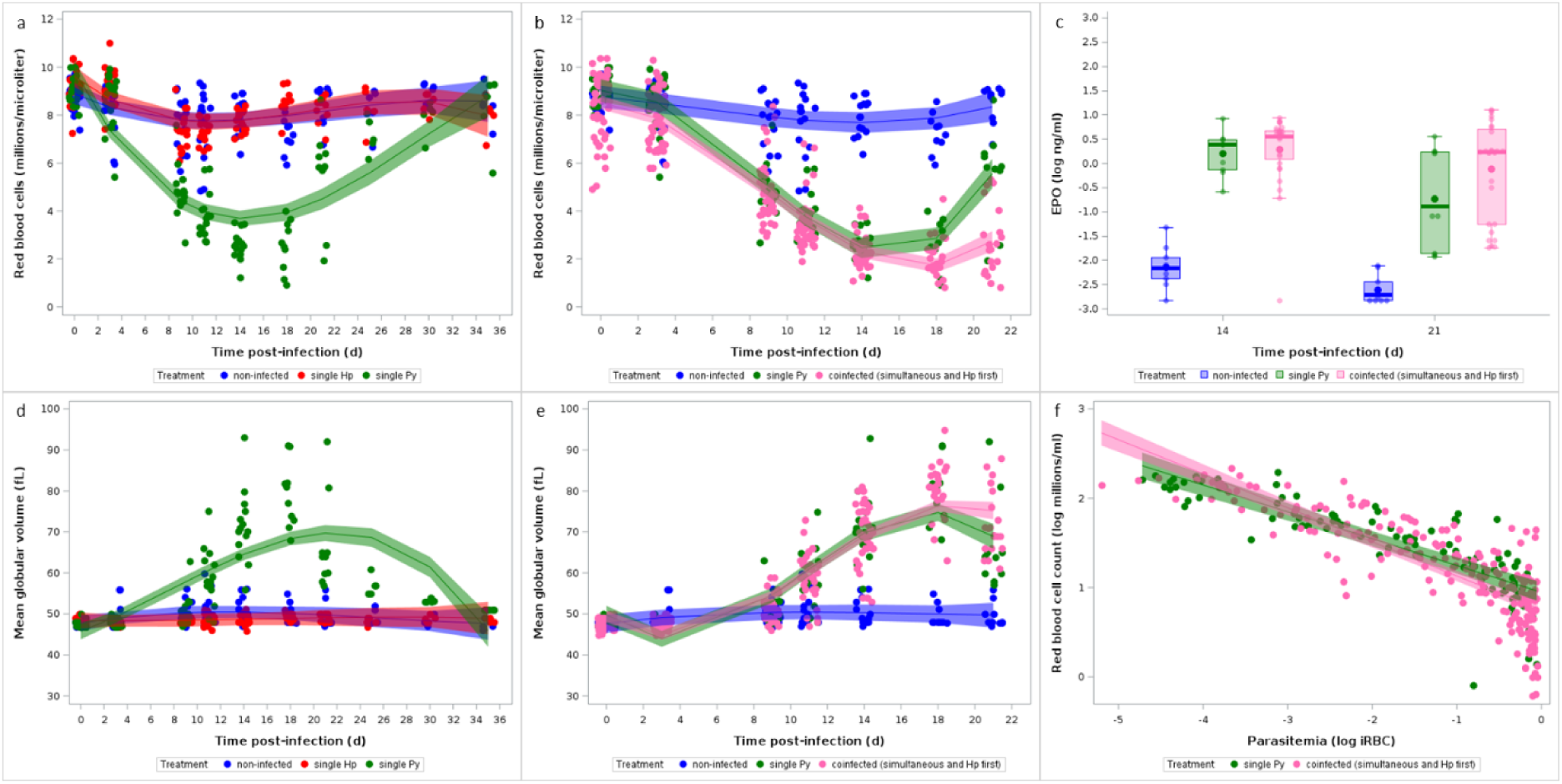
Effect of infection and coinfection on RBCs. a) Changes in RBC counts measured by hematology analyzer for non-infected and single (Hp, Py) infected mice; b) Changes in RBCs for non-infected, single Py infected and coinfected (simultaneous infection and Hp first) groups. Dots represent the raw data, the lines represent the fit of the GLMM, and the shaded area around the line the 95% CI. c) Levels of erythropoietin (EPO, log ng/ml) in the plasma assessed by ELISA at day 14 and 21 p.i. in non-infected, single Py infected and coinfected (simultaneous infection and Hp first) mice. Dots represent the raw data, the boxes represent the interquartile range (IQR), the horizontal lines the median, and whiskers the range of data within 1.5 the IQR; d) Changes in mean globular volume (MGV, fL) assessed by hematology analyzer for non-infected and single (Hp, Py) infected mice; e) Changes in MVG for non-infected, single Py infected and coinfected (simultaneous infection and Hp first) groups. Dots represent the raw data, the lines represent the fit of the GLMM, and the shaded area around the line the 95% CI; f) Log-log plot of RBC counts on parasitemia for single Py infected and coinfected (simultaneous infection and Hp first) groups. Dots represent the raw data, the lines represent the fit of the GLMM, and the shaded area around the line the 95% CI.

**Table 5.**
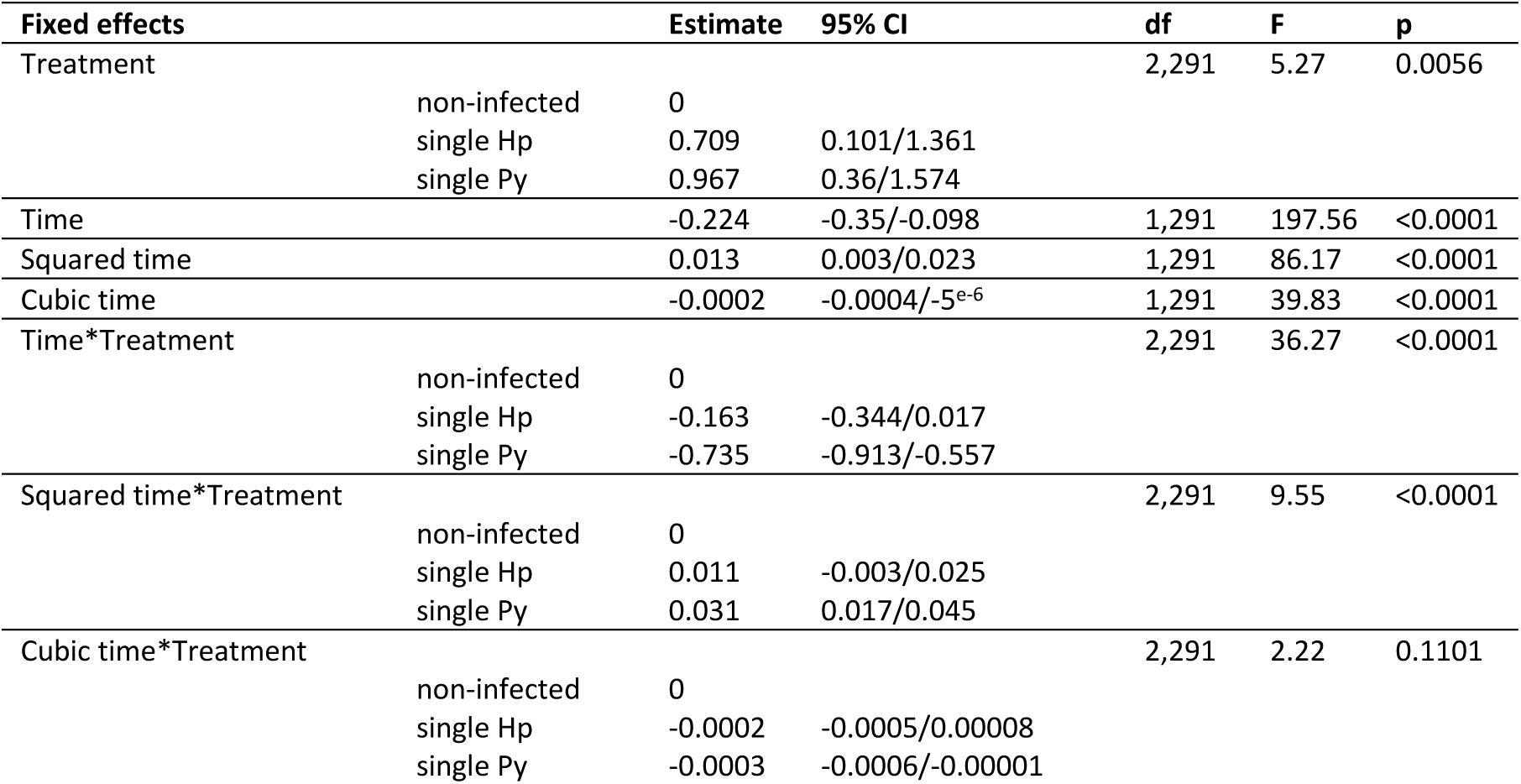
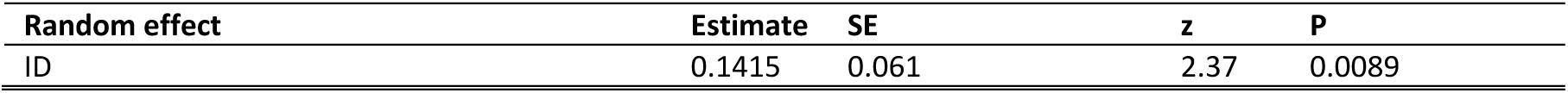
General linear mixed model investigating the changes in RBC counts (10^6^/µl) in non-infected and single (Hp, Py) infected mice. We report the parameter estimates with the 95% CI, degrees of freedom (df), F and p values. Mouse ID was included as a random effect to take into account the non-independence of observations for the same individual over time. N = 60 individuals and 360 observations.

Coinfected groups (groups 4, 5, 6) had a similar drop in the count of RBCs (table S13, fig. S12a,b), and we clustered them and compared the changes in RBC counts between non-infected, single Py infected and coinfected mice (groups 4, 5, 6). The model showed that coinfected mice still had low levels of RBCs at day 21 p.i. compared to single Py infected mice that had already started the recovery phase (table 6, fig. 5b, fig. S12c).

**Table 6.**
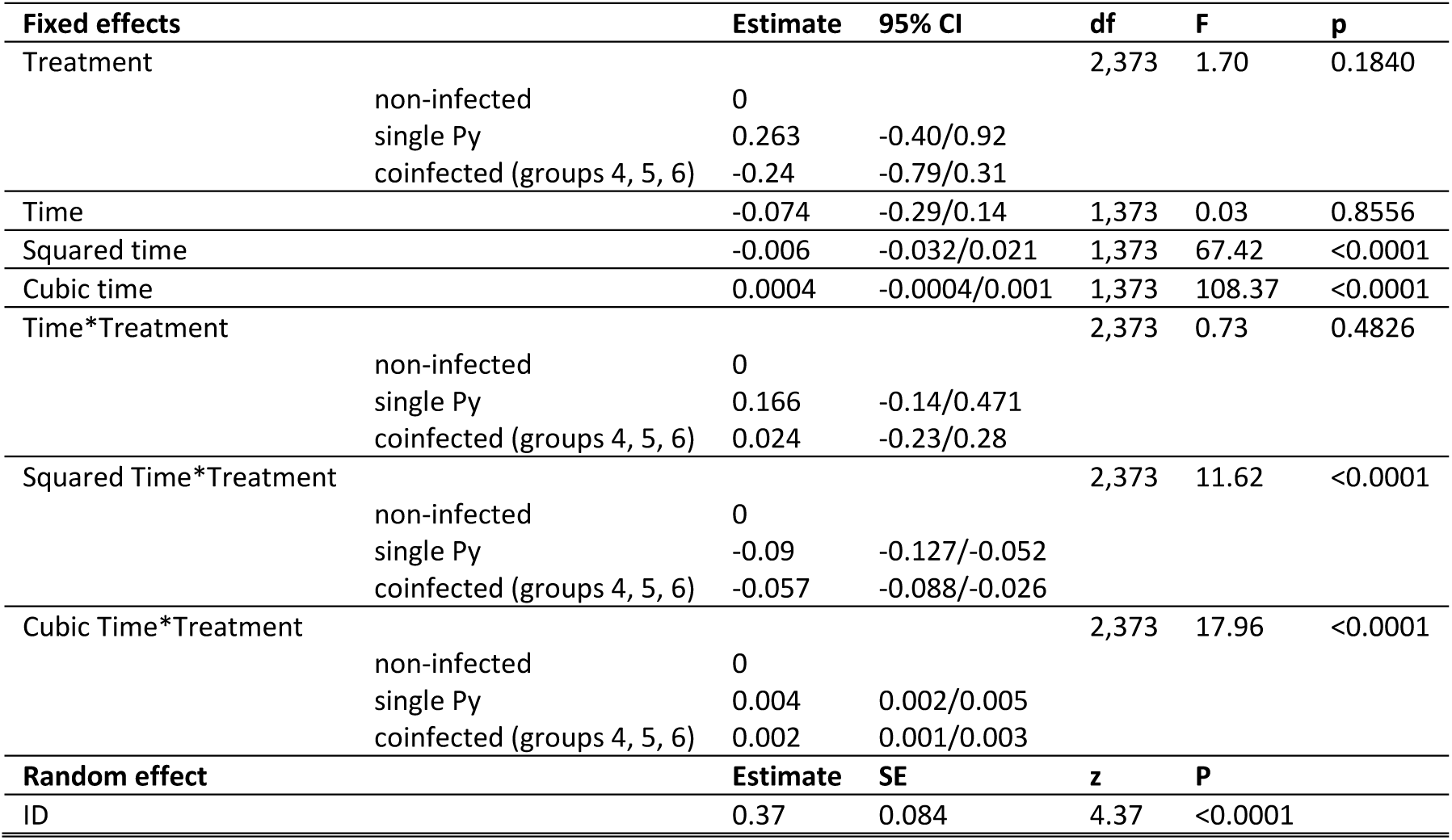
General linear mixed model investigating the changes in RBC count (10^6^/µl) in non-infected, single Py infected and coinfected (simultaneous infection and Hp first) mice. We report the parameter estimates with the 95% CI, degreed of freedom (df), F and p values. Mouse ID was included as a random effect to take into account the non-independence of observations for the same individual over time. N = 94 individuals and 476 observations.

Infection-induced anemia stimulates erythropoiesis with the aim of restoring homeostatic RBC counts. We therefore investigated whether coinfection impaired the host capacity to repair the damage (here the loss of RBCs) caused by the infection. To this purpose, we assessed the production of erythropoietin (EPO) (at day 14 and 21 p.i.) and the mean globular volume that, in the case of malaria-induced anemia, reflects the proportion of reticulocytes among the total RBCs.

Mice with single Py infection produced EPO in response to the loss of RBCs compared to non-infected individuals (GLM, F_1,34_ = 113.53, p < 0.0001, n = 38; fig. 5c). However, despite having lower RBC counts, coinfected hosts (groups 4, 5, 6) did not produce more EPO compared to single Py infected mice, neither when including each coinfection group separately in the model (GLM, F_3,67_ = 2.24, p = 0.0917, n = 75; fig. S13), nor when grouping them (post hoc comparison of LS-means between single infection and coinfection with Bonferroni correction, p = 0.3096, n = 94; fig. 5c)

This result was mirrored by the change of MGV over the course of the infection. Py infected mice had a sharp increase in the MGV between day 11 and 21 p.i., compared to non-infected and Hp infected mice, indicating a higher proportion of reticulocytes among RBCs (table 7, fig. 5d, fig. S14). However, changes in MGV were similar between single Py infection and the coinfection groups (groups 4, 5, 6) (table 8, fig. 5e, table S14, fig. S15).

**Table 7.**
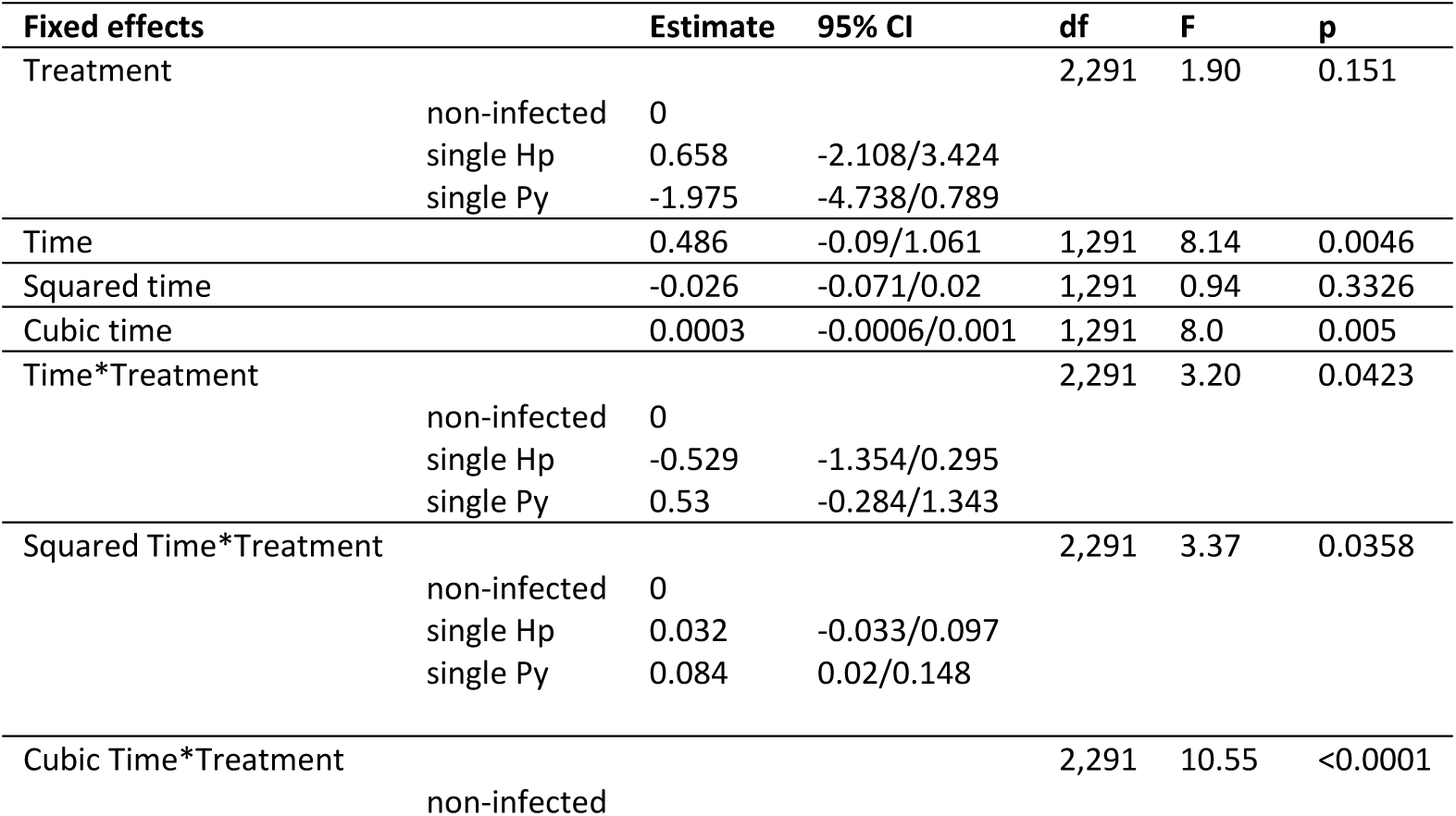
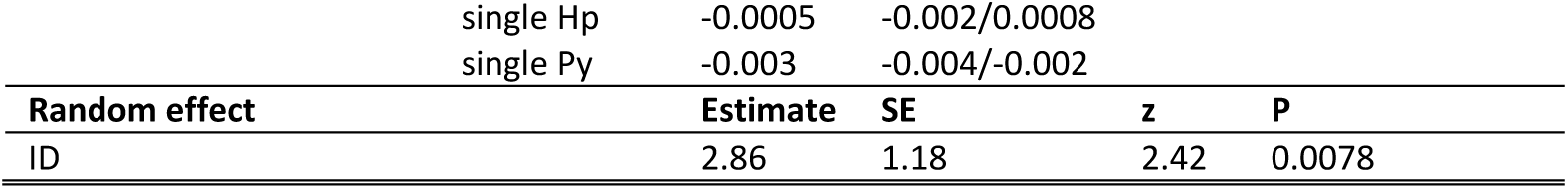
General linear mixed model investigating the changes in mean globular volume (fL) in non-infected and single (Hp, Py) infected mice. We report the parameter estimates with the 95% CI, degrees of freedom (df), F and p values. Mouse ID was included as a random effect to take into account the non-independence of observations for the same individual over time. N = 60 individuals and 360 observations.

**Table 8.** General linear mixed model investigating the changes in mean globular volume (fL) in non-infected, single Py infected and coinfected (simultaneous infection and Hp first) mice. We report the parameter estimates with the 95% CI, degrees of freedom (df), F and p values. Mouse ID was included as a random effect to take into account the non-independence of observations for the same individual over time. N = 94 individuals and 476 observations.

| Fixed effects |  | Estimate | 95% CI | Df | F | P |
| --- | --- | --- | --- | --- | --- | --- |
| Treatment |  |  |  | 2,373 | 1.05 | 0.3518 |
|  | non-infected | 0 |  |  |  |  |
|  | single Py | 2.1 | -1.05/5.25 |  |  |  |
|  | coinfection (groups 4, 5, 6) | 1.7 | -0.91/4.31 |  |  |  |
| Time |  | 0.6 | -0.54/1.75 | 1,373 | 45.87 | <0.0001 |
| Squared time |  | -0.04 | -0.18/0.1 | 1,373 | 111.07 | <0.0001 |
| Cubic time |  | 0.0008 | -0.004/0.005 |  | 109.01 |  |
| Time*Treatment |  |  |  | 2,373 | 17.09 | <0.0001 |
|  | non-infected | 0 |  |  |  |  |
|  | single Py | -4.36 | -5.98/-2.73 |  |  |  |
|  | coinfection (groups 4, 5, 6) | -3.56 | -4.92/-2.21 |  |  |  |
| Squared Time*Treatment |  |  |  | 2,373 | 30.02 | <0.0001 |
|  | non-infected | 0 |  |  |  |  |
|  | single Py | 0.71 | 0.51/0.90 |  |  |  |
|  | coinfection (groups 4, 5, 6) | 0.58 | 0.41/0.74 |  |  |  |
| Cubic Time*Treatment |  |  |  | 2,373 | 27.42 | <0.0001 |
|  | non-infected | 0 |  |  |  |  |
|  | single Py | -0.02 | -0.03/0.016 |  |  |  |
|  | coinfection (groups 4, 5, 6) | -0.02 | -0.02/0.012 |  |  |  |
| Random effect |  | Estimate | SE | Z | p |  |
| ID |  | 4.28 | 1.25 | 3.44 | 0.0003 |  |

Given that coinfected hosts had a higher proportion of iRBCs than single infected hosts, a similar rate of renewal of lost RBCs might result in a reduction in tolerance in coinfected mice. We defined tolerance as the slope of the regression of RBC counts on parasitemia, the steeper the slope the weaker the tolerance (i.e., the capacity to cope with the damage induced by the infection). We found that coinfected hosts (groups 4, 5, 6) were less tolerant (i.e., steeper slope) to Py infection compared to single infected mice (table 9; fig. 5f).

**Table 9.** General linear mixed model investigating the relationship between RBC counts (log 10^6^/µl) and parasitemia (log proportion iRBC) in single Py infected and coinfected (simultaneous infection and Hp first) mice. We report the parameter estimates with the 95% CI, degrees of freedom (df), F and p values. Mouse ID was included as a random effect to take into account the non-independence of observations for the same individual over time. N = 48 individuals and 298 observations.

| Fixed effects | Estimate | 95% CI | Df | F | P |
| --- | --- | --- | --- | --- | --- |
| Treatment |  |  | 1,248 | 8.71 | 0.0035 |
| single *Py* | 0 |  |  |  |  |
| Coinfection (groups 4, 5, 6) | -0.199 | -0.331/-0.066 |  |  |  |
| Parasitemia | -0.300 | -0.341/-0.259 | 1,248 | 646.17 | <0.0001 |
| Treatment *Parasitemia |  |  | 1,248 | 9.14 | 0.0028 |
| single *Py* | 0 |  |  |  |  |
| Coinfection (groups 4, 5, 6) | -0.081 | -0.134/-0.028 |  |  |  |
| Random effect | Estimate | SE | Z | p |  |
| Mouse ID | 0.015 | 0.006 | 2.51 | 0.0061 |  |

If coinfected hosts suffer from more severe disease symptoms because they cannot effectively restore RBCs lost during the infection, we might expect that experimentally inducing erythropoiesis, at least partially, dampen the cost of coinfection. To explore this hypothesis, we ran an experiment where mice were first infected with Hp and after 28 days they were infected with Py. Mice were then treated with a rEPO or left as control. We assessed the severity of the symptoms using an integrative marker of health (body mass), a specific marker of Py infection (changes in RBCs), and any mortality (reaching of the end points) that might have occurred during the course of the infection.

Treating coinfected mice with rEPO reduced the severity of the disease symptoms, since body mass remained stable over the course of the infection, while mice in the control group lost body mass from day 14 p.i. onwards (table 10, fig. 6a). rEPO treatment only slightly affected anemia (GLMM, time p.i.*treatment, F_1,129_ = 4.12, p = 0.0443, n = 20 individuals and 153 observations; fig. 6b, fig. S16), and although mortality was higher in the control group, the difference was not statistically significant (Log-rank test, χ²_1_ = 1.059, p = 0.3034, n = 20; fig. 6c), suggesting that other aspects of the pathophysiology of coinfection might explain the observed increase in Py virulence (although it should be acknowledged that larger sample size might have produced a statistically significant difference between groups). Nevertheless, the rEPO administration improved the tolerance to Py infection, since coinfected mice treated with rEPO had similar slope relating RBC counts to parasitemia as single Py infected hosts (GLMM, treatment*parasitemia, F_1,111_ = 0.11, p = 0.7363, n = 25 individuals and 138 observations; fig. 6d). Therefore, coinfection reduced the tolerance to Py infection and administering rEPO partially restored it.

**Figure 6.**
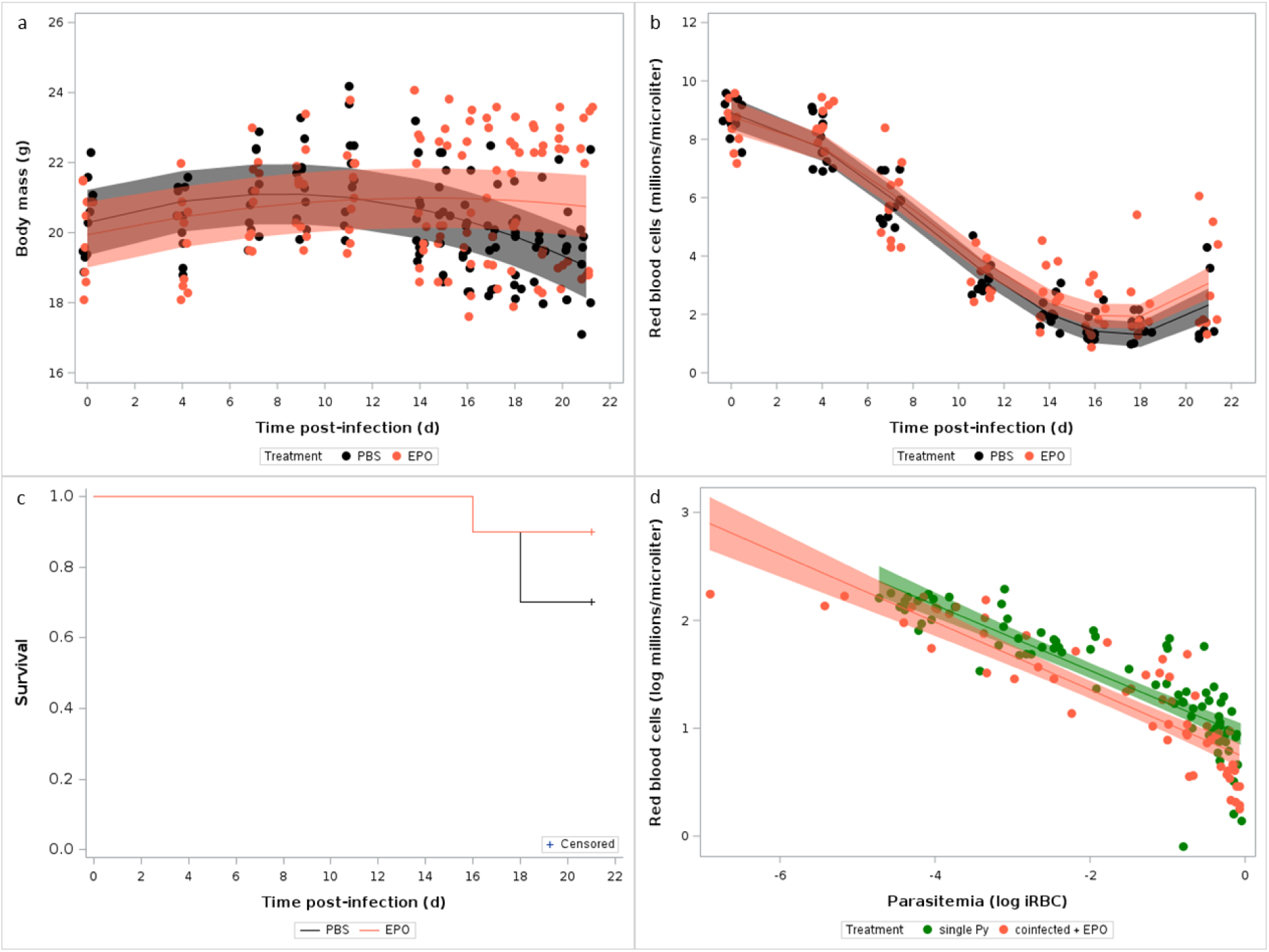
Effect of rEPO administration on the symptoms of the coinfection. a) Changes in body mass (g) over the course of the infection in mice that were infected with Hp at day −28 and with Py at day 0, and treated with rEPO or left as control (PBS); b) Changes in RBC counts assessed by hematology analyzer in coinfected mice treated with EPO or left as control (PBS). Dots represent the raw data, the lines represent the fit of the GLMM, and the shaded area around the line the 95% CI; c) Survival of coinfected mice treated with rEPO or left as control (PBS); d) Log-log plot of RBC counts on parasitemia for single Py infected mice and coinfected (infected with Hp at day −28 and with Py at day 0) mice treated with rEPO. Dots represent the raw data, the lines represent the fit of the GLMM, and the shaded area around the line the 95% CI.

**Table 10.**
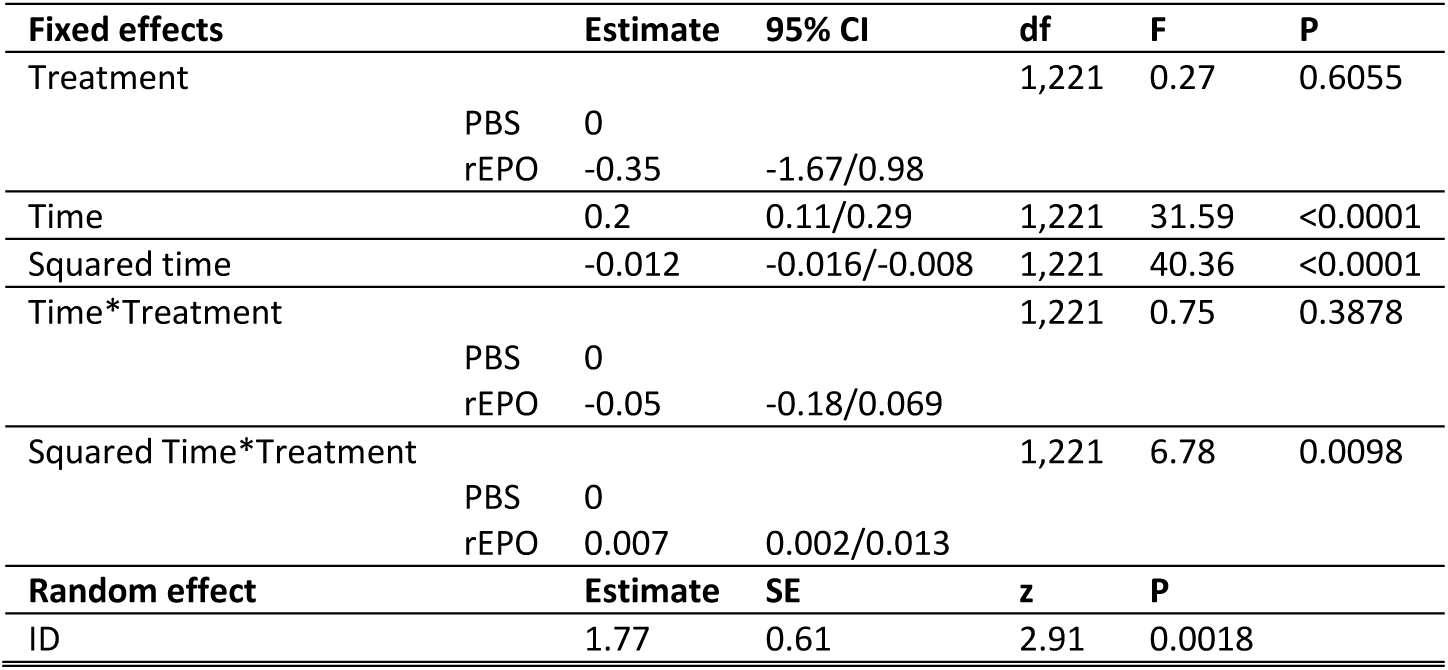
General linear mixed model investigating the changes in body mass (g) in coinfected (Hp-28+Py) mice treated with rEPO or with PBS. We report the parameter estimates with the 95% CI, degrees of freedom (df), F and p values. Mouse ID was included as a random effect to take into account the non-independence of observations for the same individual over time. N = 20 individuals and 245 observations.

In addition to inducing anemia, the lysis of RBCs releases free heme groups that have pro-oxidant and pro-inflammatory properties, and stimulates the production of enzymes whose role is to scavenge them. We therefore hypothesized that the increased virulence of Py in coinfected hosts (when Py follows Hp) might stem from an impaired capacity to detoxify free heme groups. To this purpose, we assessed the expression of the *Hmox1* gene in splenocytes at day 14 p.i. and the amount of circulating heme oxygenase-1 (HO-1) in plasma at day 14 and 21 post last infection. *Hmox1* was strongly upregulated in Py infected mice compared to non-infected individuals (GLM, F_1,8_ = 96.92, n = 10; p < 0.0001; fig. 7a, fig. S17a), and it was also upregulated in coinfected groups (groups 4, 5, 6) compared to single Py infected hosts (post-hoc comparison of LS-means, p = 0.0074, n = 28; fig. 7a, fig. S17a,b). However, although single Py infected mice also had much higher plasma levels of the enzyme HO-1 at both day 14 and 21 p.i. compared to non-infected individuals (GLM, F_1,24_ = 82.98, p < 0.0001, n = 28; fig. 7b), coinfected groups (groups 4, 5, 6) did not differ from single Py infection (when considering the different coinfection groups separately, GLM, F_3,68_ = 2.49, p = 0.0677, n = 76; fig. S17c; when considering the coinfection groups together, post-hoc comparison of LS-means, p = 1, n = 86; fig. 7b). Therefore, although *Hmox1* gene expression was upregulated in the coinfection groups, the level of the enzyme in the blood did not differ between single infection and coinfection.

**Figure 7.**
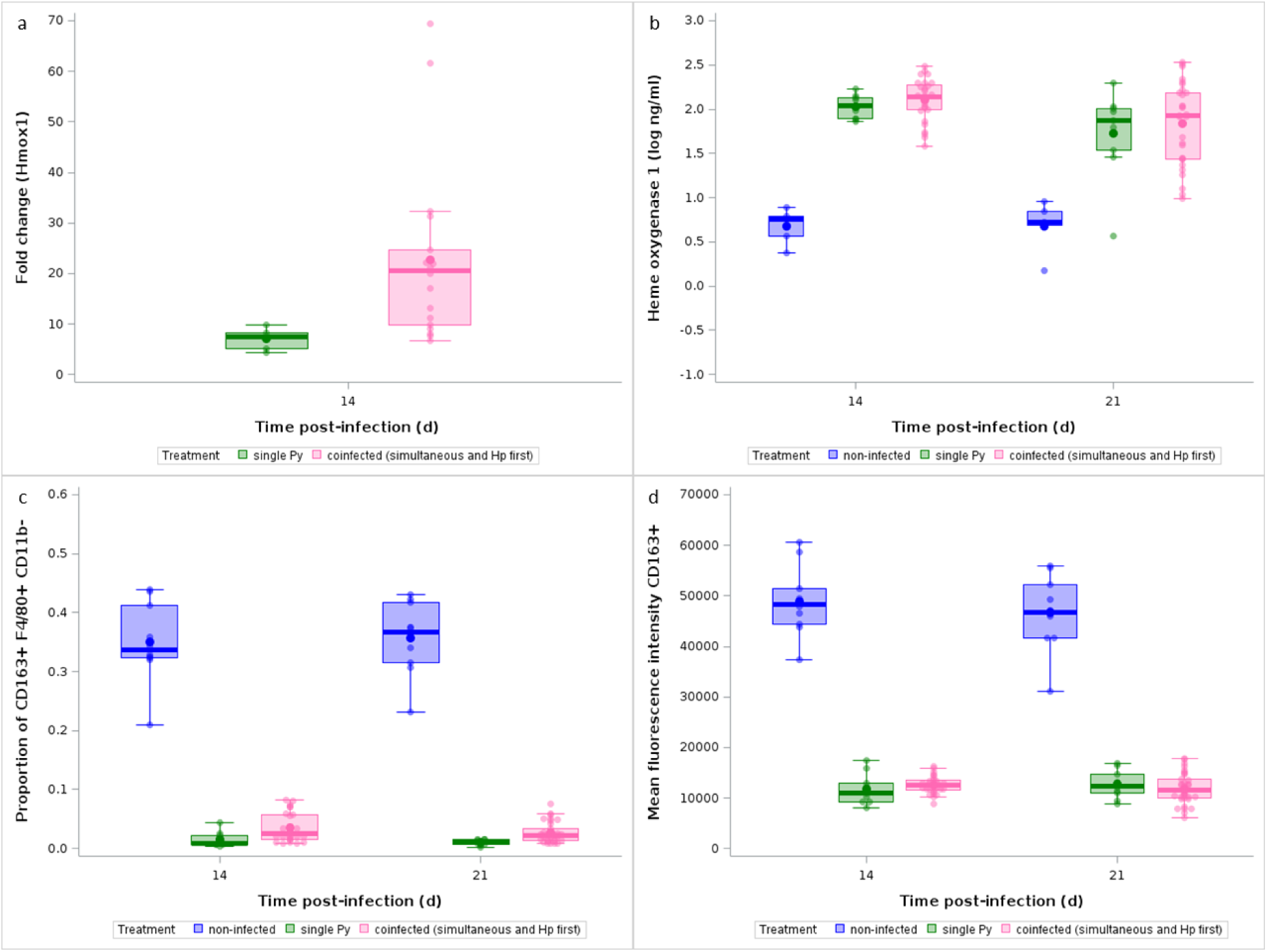
Effect of infection and coinfection on heme detoxification. a) Fold change relative to non-infected mice of Hmox1 mRNA in spleen at day 14 p.i. in single Py infected and coinfected (simultaneous infection and Hp first) groups; b) Levels of Heme oxygenase-1 in the plasma (log ng/ml) at day 14 and 21 p.i. in non-infected, single Py infected and coinfected (simultaneous infection and Hp first) groups; c) Proportion of CD163^+^ within F4/80^+^ CD11b^−^ cells analyzed by flow cytometry at day 14 and 21 p.i. in non-infected, single Py infected and coinfected (simultaneous infection and Hp first) groups; d) Mean fluorescence intensity (arbitrary fluorescence units) of CD163 at day 14 and 21 p.i. in non-infected, single Py infected and coinfected (simultaneous infection and Hp first) groups. Dots represent the raw data, the boxes represent the interquartile range (IQR), the horizontal lines the median, and whiskers the range of data within 1.5 the IQR.

Another important function related to the clearance of free hemoglobin is provided by the scavenger receptor CD163 expressed on red pulp macrophages (RPM, F4/80^+^ CD11b^−^) (Kristiansen et al. 2001). We therefore assessed the proportions of RPM that were CD163^+^, by flow cytometry, and compared them between non-infected and single Py infected and found that the population of CD163^+^ RPM drastically dropped during the infection (generalized linear model with a beta distribution of errors, F_1,35_ = 344.88, p < 0.0001, n = 39; fig. 7c). However, the proportion of RPM that were CD163^+^ was similarly reduced in coinfected mice (groups 4, 5, 6) compared to single Py infection, at both day 14 and 21 p.i. (LS-means comparison with Bonferroni correction, day 14 p.i., p = 0.1648, day 21 p.i., p = 0.1959, n = 100; fig. 7c, fig. S17d). The receptor CD163 was also less expressed in F4/80^+^ CD11b^−^ of Py infected mice, as shown by the drastic drop in mean fluorescence intensity (MFI), with no difference between single and coinfected mice (LS-means comparison with Bonferroni correction, day 14 p.i., p = 1, day 21 p.i., p = 1, n = 100; fig. 7d). Therefore, the population of CD163^+^ RPM which are a key resident cell population responsible for clearing free hemoglobin released during the lysis of RBCs similarly decreased in single and coinfected mice.

### Immune predictors of Py virulence

*Plasmodium* infection elicits proinflammatory effectors that might cause immunopathology if regulatory mechanisms fail to restore homeostasis. Therefore, we speculated that exacerbated disease severity in coinfected hosts (when Py follows Hp) might stem from an overreacting, non-regulated, inflammatory response. To address this question, we first assessed the gene expression of pro- (IFN-γ) and anti-inflammatory (IL-10, TGF-β) cytokines in splenocytes at day 3 and 14 post-infection. IFN-γ, IL-10 and TGF-β1 genes were all overexpressed at both 3 and 14 days p.i. in single Py infected hosts compared to non-infected mice (GLM, IFN-γ, F_1,34_ = 24.44, p < 0.001; IL-10, F_1,34_ = 49.86, p < 0.001; TGF-β, F_1,34_ = 20.62, p < 0.001, n = 38; fig. 8; fig. S18a,b,c). However, coinfection (groups 4, 5, 6) did not induce a change in the expression of any of the genes compared to single Py infection (LS-means comparison with Bonferroni correction, all p’s = 1, fig. 8, fig. S18d,e,f).

**Figure 8.**
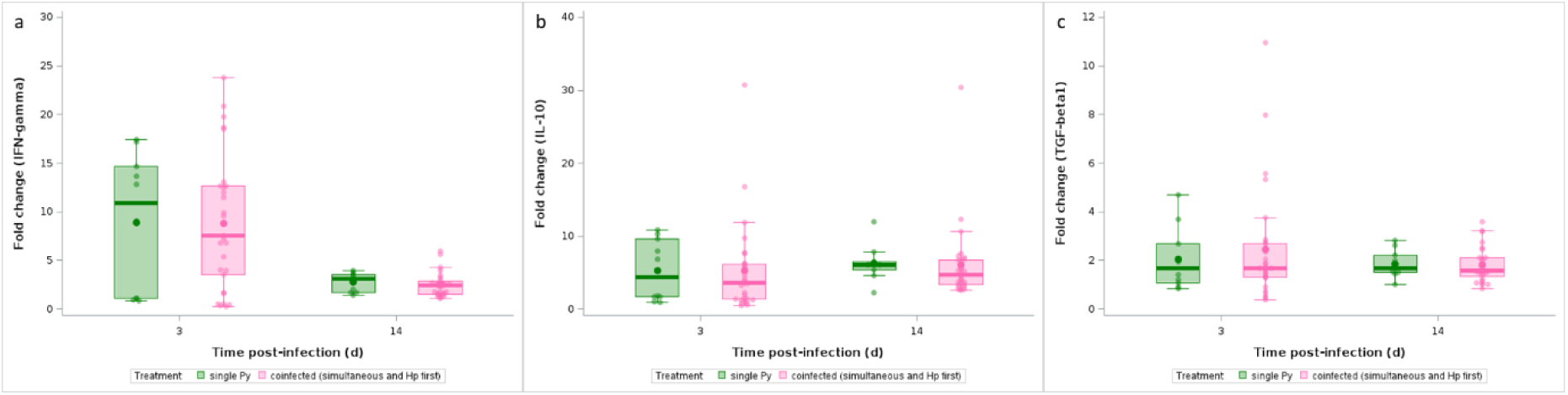
Effect of infection and coinfection on cytokine expression. Fold change (relative to non-infected hosts) of a) IFN-γ, b) IL-10, c) TGF-β1 mRNA levels in spleen at day 3 and 14 p.i. in single Py infected and coinfected (simultaneous infection and Hp first) groups. Dots represent the raw data, the boxes represent the interquartile range (IQR), the horizontal lines the median, and whiskers the range of data within 1.5 the IQR.

Previous work conducted on the same model system has shown that coinfected mice have a higher activation of regulatory T cells (Treg), possibly accounting for their impaired anti-*Plasmodium* response (Tetsutani et al. 2009). Therefore, we assessed the proportions of CD4^+^ T cells that were FoxP3^+^ by flow cytometry and compared them, over the course of the infection, between non-infected, single Py infected and coinfected mice (groups 4, 5, 6); we found no differences between single Py infection and coinfection groups (LS-means comparison with Bonferroni correction, p = 1; n = 100; fig. 9a, table S15; fig. S19a). The expression of the immune checkpoint CTLA-4 on Treg has been shown to impede immunity to the acute infection with Py (Kurup et al. 2017), by enhancing their immunosuppressive activity. We thus assessed the proportions of CD4^+^ T cells that were FoxP3^+^ and CTLA-4^+^ by flow cytometry and compared them between non-infected, single Py infected and coinfected mice (groups 4, 5, 6); we found that coinfected mice had a higher proportion of CTLA-4^+^ Treg compared to non-infected and single Py infection at day 14 p.i. (table 11; fig. 9b; table S16; fig. S19b). Therefore, although coinfection did not produce an expansion of the Treg population compared to single Py infected hosts, the immunosuppressive power of Treg was upregulated in coinfected hosts.

**Figure 9.**
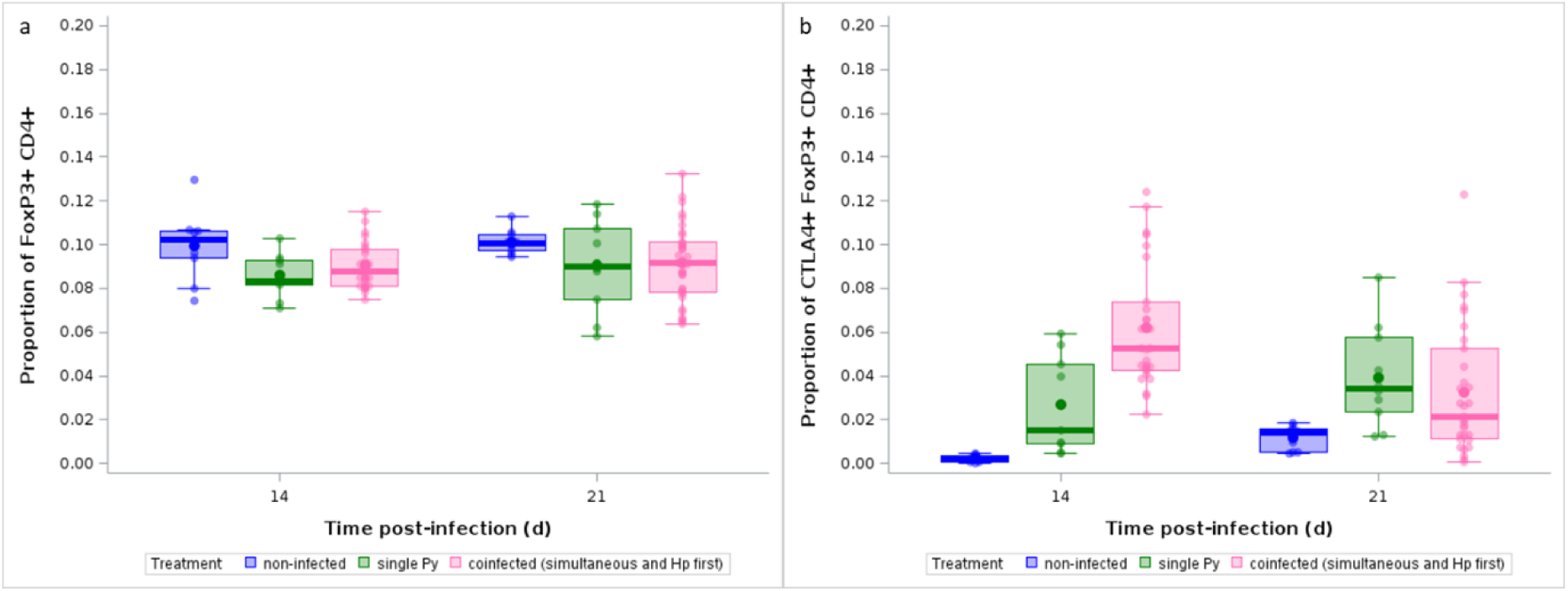
Effect of infection and coinfection on Treg expansion and immunosuppressive function. a) Proportion of FoxP3^+^ within CD4^+^ T cells analyzed by flow cytometry over the course of the infection in non-infected, single Py infected and coinfected (simultaneous infection and Hp first) mice; b) Proportion of CTLA-4^+^ within FoxP3^+^ CD4^+^ T cells analyzed by flow cytometry over the course of the infection in non-infected, single Py infected and coinfected (simultaneous infection and Hp first) mice. Dots represent the raw data, the boxes represent the interquartile range (IQR), the horizontal lines the median, and whiskers the range of data within 1.5 the IQR.

**Table 11.** Generalized linear model with a beta distribution of errors investigating the changes in the proportion of CTLA-4^+^ within FoxP3^+^ CD4^+^ T cells in non-infected, single Py infected and coinfected (simultaneous infection and Hp first) mice at day 14 and 21 post-infection. We report the degrees of freedom (df), and the F and p values. N = 96 individuals.

| Source of variation | df | F | P |
| --- | --- | --- | --- |
| Treatment | 2,90 | 19.01 | <0.0001 |
| Time | 1,90 | 0.79 | 0.3755 |
| Time*Treatment | 2,90 | 13.89 | <0.0001 |

Using flow cytometry, we also immunophenotyped mice using a large panel of markers in the spleen (table S2), at day 3, 14 and 21 post last infection. We identified 20 cell populations (table S17) and their proportions were included as variables in a principal component analysis (PCA), with a PCA per sampling day. The first two components explained 29.02% and 18.79%, 39.65% and 12.86%, 37.63% and 15.92% of the total variance, at day 3, 14, and 21 p.i., respectively. The correlation circles showed that many of the variables included in the PCA had high absolute loadings (table S17, fig. S20a, b, c). At day 3 p.i., the scores along the first principal component (PC1) strongly differed between the three experimental groups (GLM, F_2,45_ = 28.56, n = 48; p < 0.0001), with coinfected mice (groups 4, 5, 6) having higher scores compared to both non-infected and single Py infection (LS-means comparisons with Bonferroni correction, both p’s < 0.0001; fig. S20d). At day 14 and 21 p.i., there was a complete separation, along the PC1, between the three groups, with single Py infected lying in between non-infected and coinfected individuals (groups 4, 5, 6) (day 14 p.i.: GLM, F_2,43_ = 313.37, p < 0.0001, n = 46; day 21 p.i.: GLM, F_2,51_ = 257.50, p < 0.0001, n = 54; fig. S20d). The PC2 had a much less discriminant power among groups (day 3 p.i.: GLM, F_2,45_ = 3.02, p = 0.0588, n = 48; day 14 p.i.: GLM, F_2,43_ = 1.27, p = 0.2902, n = 46; day 21 p.i.: GLM, F_2,51_ = 7.57, p = 0.0013, n = 54; fig. S20e).

To go further, we focused on the variables that had strong absolute loadings on the PC1 and compared them between the groups, with the aim to see if there was a consistent difference between single Py infected and coinfected hosts (groups 4, 5, 6). The proportion of Th2 (proportion of CD4^+^ T cells that were GATA-3^+^) differed between single and coinfected hosts at the three sampling dates (GLM with a beta distribution of errors, F_2,139_ = 53.91, p < 0.0001, n = 148; LS-means comparison between single and coinfected with Bonferroni correction, day 3, p = 0.0002; day 14, p = 0.0102; day 21, p < 0.0001; fig. 10a). Among the other variables, only CD8^+^ T cells consistently discriminated between single Py and coinfected hosts (groups 4, 5, 6), with the proportion of CD8^+^ T cells (proportion of CD3^+^ that were CD8^+^) sharply decreasing over time and especially so in coinfected hosts (GLM with a beta distribution of errors, F_2,139_ = 347.41, p < 0.0001, n = 148; LS-means comparison between single and coinfected with Bonferroni correction, day 3, p = 1; day 14, p < 0.0001; day 21, p < 0.0001, fig. 10b).

**Figure 10.**
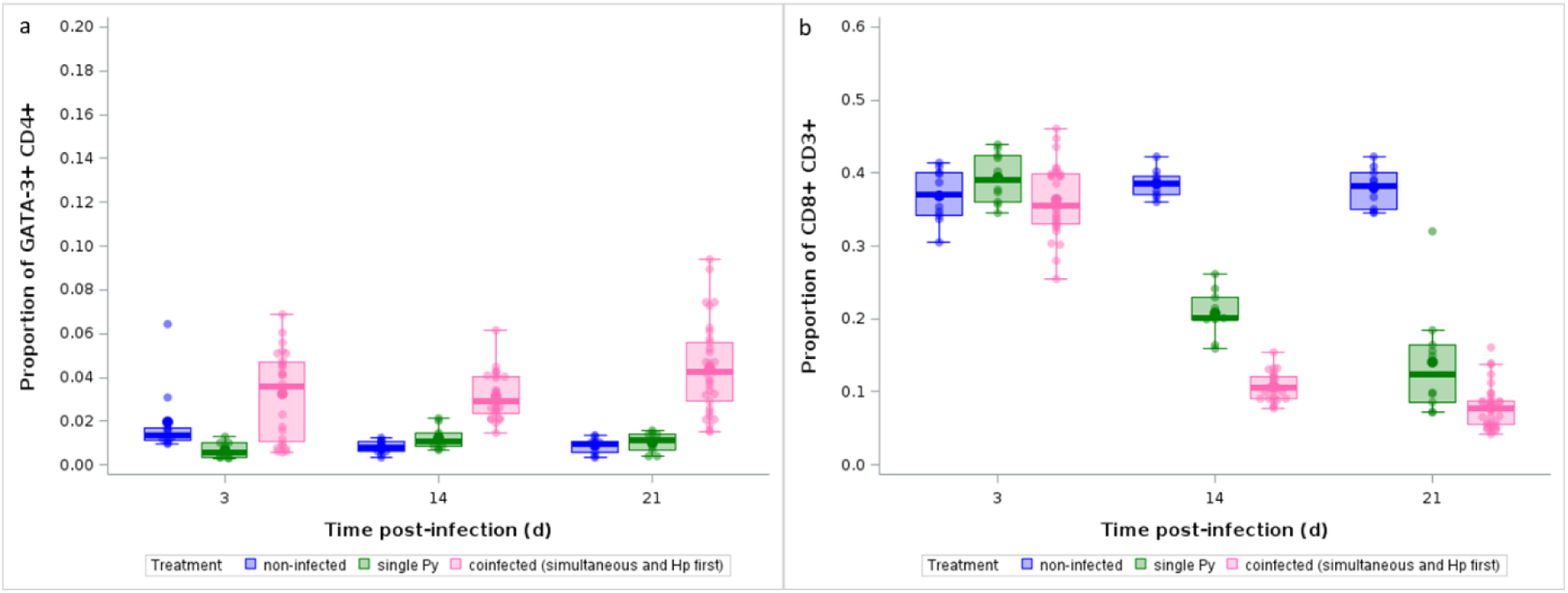
Effect of infection and coinfection on Th2 and CD8^+^ T cells. a) Proportion of GATA-3^+^ within CD4^+^ T cells and b) proportion of CD8^+^ within CD3^+^ T cells analyzed by flow cytometry over the course of the infection in non-infected, single Py infected and coinfected (simultaneous infection and Hp first) hosts. Dots represent the raw data, the boxes represent the interquartile range (IQR), the horizontal lines the median, and whiskers the range of data within 1.5 the IQR.

Therefore, Th2 and CD8^+^ T cells were the two populations that best discriminated between Py infected and coinfected hosts (when Py follows Hp). While the effect of coinfection on the rise of Th2 lymphocytes can easily be understood as the direct consequence of Hp induction of Th2 immunity, we wished to better investigate the reasons underlying the decrease of CD8^+^ T cells in coinfected mice. To this purpose, we assessed the proportion of CD8^+^ T cells that expressed three immune checkpoints in non-infected, single Py infected and coinfected hosts (groups 4, 5 and 6), and predicted that coinfected mice should have higher proportions of CD8^+^ T cells expressing the immune checkpoints compared to single Py infection. Py infection enhanced the proportion of CTLA-4^+^ CD8^+^ T cells (proportion of CD8^+^ that were CTLA-4^+^) but there was no difference between single infection and coinfection (groups 4, 5 and 6) at any of the sampling day (GLM with a beta distribution of errors: day 14 p.i, t = −0.52, df = 90, p = 1; day 21 p.i., t = −1.19, df = 90, p = 1; fig. 11a). The proportions of LAG-3^+^ CD8^+^ and PD-1^+^ CD8^+^ cells (proportion of CD8^+^ that were LAG-3^+^ and PD-1^+^, respectively) were also increased in infected mice compared to non-infected individuals, however, at day 14 p.i. there was also a higher proportion of LAG-3^+^ and PD-1^+^ CD8^+^ T cells in coinfected mice compared to single Py infection (GLM with a beta distribution of errors: LAG-3, day 14 p.i.: t = 3.38, df = 91, p = 0.0163; day 21 p.i., t = 2.36, df = 91, p = 0.3040; PD-1, day 14 p.i.: t = 3.94, df = 90, p = 0.0024; day 21 p.i., t = 2.26, df = 91, p = 0.3948, fig. 11b,c).

**Figure 11.**
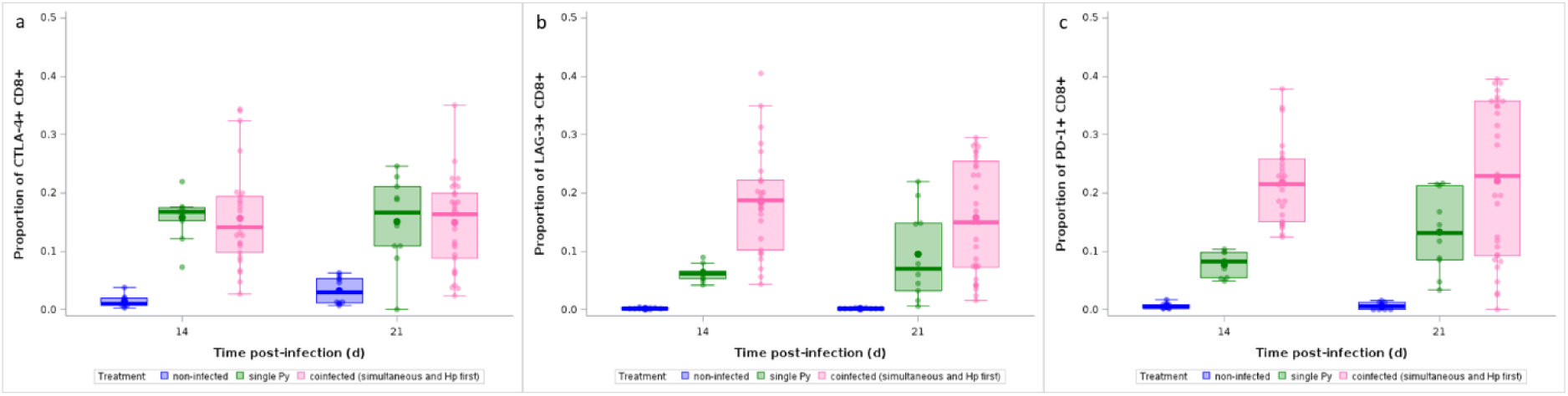
Effect of infection and coinfection on CD8^+^ T cell exhaustion. a) Proportion of CTLA-4^+^ within CD8^+^ T cells, b) LAG-3^+^ within CD8^+^ T cells, c) PD-1^+^ within CD8^+^ T cells analyzed by flow cytometry at day 14 and 21 p.i. in non-infected, single Py infected and coinfected (simultaneous infection and Hp first) hosts. Dots represent the raw data, the boxes represent the interquartile range (IQR), the horizontal lines the median, and whiskers the range of data within 1.5 the IQR.

Persistent antigenic stimulation and high levels of inhibitory receptors (immune checkpoints) can lead to the exhaustion of CD8^+^ T cells, characterized by a loss of effector functions and proliferative potential (McLane et al. 2019), and to the increase in the CD4^+^/CD8^+^ ratio above the homeostatic value (between 1.5 and 2.5) (McBride and Striker 2017). Deviations from this homeostatic value have been associated with severe symptoms in a few infectious diseases (e.g., Pascual-Dapena et al. 2022). We therefore wished to investigate if high values of the CD4^+^/CD8^+^ ratio also predicted mortality in our system. We could not investigate this point at the individual level because data on the proportion of immune cells were gathered on mice that were euthanized at specific days post last infection. Nevertheless, we could correlate the CD4^+^/CD8^+^ ratio at day 14 and 21 p.i. with the mortality rate (the proportion of mice reaching the end points) within each experimental group and found that indeed the higher the ratio the higher the mortality risk (spearman correlation, Rs = 0.766, n = 100, p < 0.0001, fig. 12).

**Figure 12.**
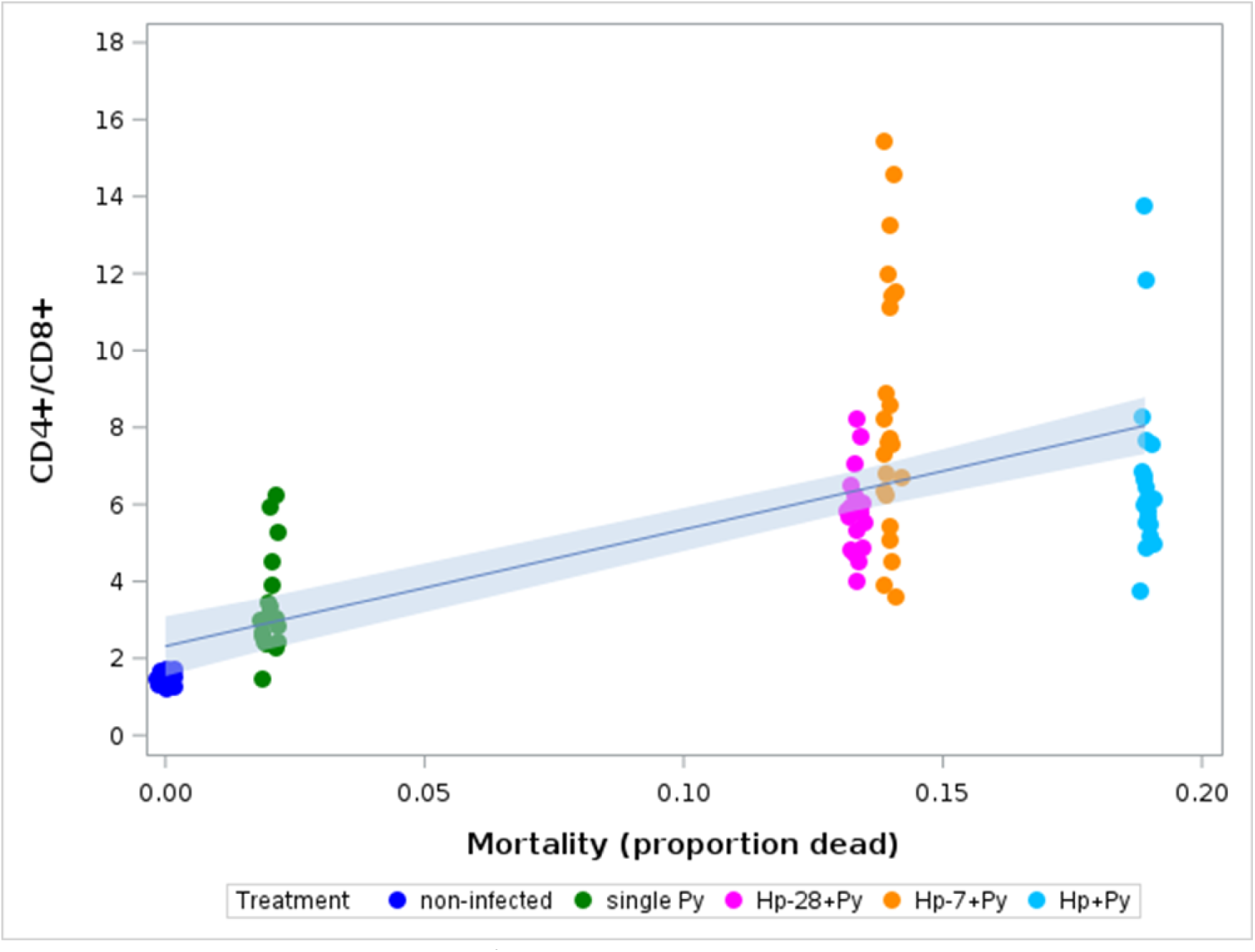
Correlation between the CD4^+^/CD8^+^ ratio at day 14 and 21 p.i. and the mortality rate in non-infected, single Py infected and coinfected (simultaneous infection and Hp first) hosts. Mortality refers to the proportion of hosts reaching the experimental end points within groups. Dots represent the raw data, the line represents a linear regression fit and the band the 95% CI of the slope.

## Discussion

We showed that the order of infection plays a paramount role in determining the outcome of the coinfection between an intestinal nematode and a *Plasmodium* parasite. In particular, Py virulence increased when infecting hosts that were previously or simultaneously infected with Hp, compared both to hosts with single infections and to hosts that were infected with Py first and then with Hp. Contrary to Py, virulence of Hp was minimally affected by the coinfection. These results are partially in agreement with our initial predictions. We hypothesized that previous infection with either Py or Hp would polarize the immune response towards Th1 or Th2 effectors making hosts more susceptible to a subsequent infection. This prediction was met when Py infected mice harboring Hp, but not when Hp infected mice harboring Py. Therefore, there is a strong asymmetry in the effect of coinfection between Py and Hp, depending on the order of the infection. The increase in Py virulence in mice previously infected with Hp is consistent with previous results on this system (Tetsutani et al. 2009), suggesting that previous Hp infection impairs the subsequent immune response to Py (Noland et al. 2008; Tetsutani et al. 2009).

The possible evolutionary reasons underlying this asymmetry are manifold. Intestinal nematodes usually produce chronic infections where the hosts keep shedding parasite eggs for months or even years depending on the species (Morand 1996). Inter-host transmission success and parasite fitness is tightly associated with the duration of egg shedding (patent period), therefore any parasite trait that compromises host health and survival is likely counter selected in these systems. In agreement with this view, nematode infections are usually well tolerated and host damage is usually observed only when parasite burden is high (Brooker 2010; Lippens et al. 2016). Since reinfection was not possible in our experiment, parasite burden was constrained by the initial number of infective larvae given to the host. It might be interesting to investigate to what extent the lack of increase in Hp virulence in coinfected hosts could change when using higher infective doses (Lippens et al. 2016), or allowing hosts to be reinfected. Although the transmission success of Py cannot be directly assessed from the proportion of iRBC, longer persistence of high parasite density, as observed in coinfected hosts, should increase the likelihood of gametocyte production, as reported for other *Plasmodium* species (see Tadesse et al. 2017). In agreement with this view, Py transmission success to the dipteran vector was enhanced when mice were coinfected with the helminth *Echinostoma caproni* (Noland et al. 2007). Therefore, the cost of virulence is probably higher in Hp than in Py. Alternatively, increased Py virulence might be maladaptive if host overexploitation and death jeopardize parasite fitness through reduced transmission success.

Whatever the benefit or disadvantage of changes in virulence for Py, from the host perspective, coinfection did incur costs both in terms of general health and specific markers of Py infection. In particular, while single infected hosts still gained mass during the 5 weeks p.i., at a rate that was similar to the non-infected hosts, coinfection induced a loss of body mass (simultaneous infection and when Py infected mice previously infected with Hp) or temporarily slowed down the gain in body mass (when Hp infected mice previously infected with Py). In agreement with the results on changes in body mass, host mortality (reaching of the end points) was only reported in coinfection groups where Py followed the Hp infection (and the simultaneous infection). Interestingly, there was no difference according to the timing of the infection, suggesting that the importance of the order of the infection is conserved and relatively consistent over different coinfection scenarios. In another coinfection model, involving the cestode *Taenia crassiceps* and *Plasmodium yoelii* 17XL, Salazar-Castañón et al. (2018) found that coinfected mice had relatively longer survival time compared to single Py infected hosts and that the gain in survival was related to the timing of the infection (the longer the delay between the infection, the better the gain in survival time), suggesting that the effect of timing of infection might be model-dependent.

Our prediction of increased virulence in coinfected hosts was based on the rationale that previous infection with nematodes polarizes the immune system towards a Th2 response (Reynolds et al. 2012). In agreement with this hypothesis, we found that Th2 cells discriminated the three experimental groups, with higher percentage of GATA-3^+^ within CD4^+^ T cells in coinfected hosts (when Py follows Hp) compared to both single Py infected and non-infected mice. Accordingly, we also predicted that coinfected mice should have an impaired capacity to deal with Py infection and indeed we found that parasitemia was still high at day 21 p.i. in coinfected hosts compared to single Py infected mice. Therefore, direct damage produced by Py multiplication might account for the more severe disease symptoms observed in coinfected hosts. In agreement with this view, changes in RBC counts over the course of the infection mirrored changes in parasitemia, with coinfected groups still suffering from anemia at day 21 p.i., when single Py infected mice had already started to recover.

The fitness consequences of the damage caused by Py multiplication also depend on the host capacity to repair damaged tissues, or in other terms on the capacity to tolerate the infection (Medzhitov et al. 2012). In particular, RBCs are constantly renewed during hematopoiesis and therefore hosts that can restore lost RBCs at a faster pace should be able to better tolerate the infection (i.e., to pay a smaller infection cost) compared to hosts with a slower rate of RBC renewal (Chang et al. 2004). Given that coinfected hosts suffered from higher anemia, compensating for the higher rate of RBC loss would have needed a higher production of reticulocytes. However, we found that coinfected hosts had similar plasma levels of EPO and similar proportions of reticulocytes, as assessed by the MGV, compared to single infected hosts. Therefore, despite higher parasitemia and higher anemia, coinfected hosts only replaced RBCs at a similar rate of single infected hosts. This finding suggests that any improvement of the capacity to renew lost RBCs should reduce the cost of coinfection. Experimental administration of rEPO to coinfected hosts partially improved the symptoms of the disease, as also previously reported (Kaiser et al. 2006), and improved host tolerance to the infection. However, the magnitude of the effects was small, suggesting that other aspects of the physiopathology of the coinfection might be important determinants of the coinfection costs. Moreover, there are physiological constraints that set a limit to the maximum daily rate of RBC production, including the number of hematopoietic stem cells that can be committed to become reticulocytes or the saturation of EPO receptors (Suzuki et al. 2002, Comazzetto et al. 2019). In addition, the production of reticulocytes also replenishes the reservoir of cells that can be exploited by Py for its own replication, which can also contribute to maintain parasitemia over longer period of time and possibly exacerbates disease severity (Chang et al. 2004; Tsubata et al. 2005).

Massive release of free heme groups has the potential to damage host tissues through their prooxidant, proinflammatory and cytotoxic effects (Kumar and Bandyopadhyay 2005; Immenschuh et al. 2017). Previous studies have shown that the effectiveness of the mechanisms of detoxification of free heme groups are key determinants of host survival (Pamplona et al. 2007; Seixas et al. 2009) and liver function restoring once the parasite has been cleared (Dey et al. 2014), although the anti-inflammatory properties of HO-1 might also promote liver infection (Epiphanio et al. 2008). We therefore assessed the expression of the *Hmox1* gene in splenocytes and the amount of the enzyme HO-1 in plasma. As for the renewal of lost RBCs, given that coinfected hosts had higher amount of free hemoglobin compared to single infection (due to the higher RBC lysis), we expected to find a higher expression of *Hmox1* and more HO-1 in plasma of coinfected hosts. However, although the gene was overexpressed in coinfected mice (the gene was highly overexpressed in infected mice compared to non-infected hosts), the amount of enzyme was similar in coinfected and single Py infected hosts, possibly due to the degradation of the enzyme through the ubiquitin-proteasome system (Lin et al. 2008; Chau 2015). We also found that another mechanism allowing the host to deal with free hemoglobin (the scavenger receptor CD163) (Kristiansen et al. 2001) was downregulated in RPM of coinfected and single infected hosts.

Free heme group is also recognized as a DAMP by the immune system and can induce an overreacting inflammatory response leading to host damage (Fortes et al. 2012). Coinfection between Hp and another model species of murine malaria (*Plasmodium chabaudi* AS) has been reported to increase host mortality (Su et al. 2005), due to liver pathology involving hepatic IFN-γ, IL-17 and IL-22 (Helmby 2009). Although infected hosts overexpressed IFN-γ, IL-10 and TGF-β1 compared to non-infected mice, there was no difference between coinfected and single Py infected mice. Similarly, we did not find any difference in the percentage of FoxP3^+^ within CD4^+^ T cells between single Py infected and coinfected mice; however, the percentage of CTLA-4^+^ within FoxP3^+^ CD4^+^ T cells was higher in coinfected mice compared to single Py infected hosts during the peak of the acute infection. Therefore, although we did not find an expansion of Treg in Py infected mice (as previously shown, Tetsutani et al. 2009), the immunosuppressive capacities of Treg were upregulated in single Py infected mice, and even more so in coinfected hosts through the CTLA-4 immune checkpoint (see Kurup et al. 2017). Therefore, coinfected mice appeared to have an upregulated immunosuppressive function, possibly accounting for the failure to control Py multiplication.

We went a step further by identifying 20 immune cell populations in the spleen by spectral flow cytometry using a 27-color panel. The results of the PCA clearly showed that the three experimental groups clustered in different areas of the space defined by the first two components, and especially so at day 14 and 21 p.i., with the single Py infection group lying in between the non-infected and the coinfection groups. The cell population that best discriminated between single Py infected and coinfected hosts was the CD8^+^ T cells that got exhausted more in coinfected mice. Several Py epitopes are recognized by CD8^+^ T cells during the blood stage of the infection (Howland et al. 2015). Moreover, previous work has shown that CD8^+^ T cells play a role in the protective immunity against blood-stage infection with Py (Imai et al. 2010), and chronic infection with the nematode *Litosomoides sigmodontis* interferes with the anti-*Plasmodium* CD8^+^ T cell response (Kolbaum et al. 2012). Infection with Hp has also been shown to suppress CD8^+^ T cells in mice subsequently infected with the protozoan *Toxoplasma gondii* (Khan et al. 2008). Exhaustion of CD8^+^ cells results from the overexpression of immune checkpoints whose function is to avoid immunopathology during persistent antigenic exposure (Chandele et al. 2010; Wykes and Lewin 2018; McLane et al. 2019). In agreement with this, we found that the proportion of both PD-1^+^ and LAG-3^+^ within CD8^+^ T cells was higher in coinfected hosts compared to single Py infection at day 14 post last infection. Mechanisms supposed to protect the hosts from immunopathology (Howland et al. 2015) might thus actually make them more susceptible to parasite exploitation (Scholzen et al. 2009; Frimpong et al. 2019), and especially so when hosts are coinfected by parasites with different exploitation strategies.

One of the consequences of the CD8^+^ exhaustion is an increase of the ratio between CD4^+^ and CD8^+^ T cells. Deviations of the CD4^+^/CD8^+^ ratio from the homeostatic values (inverted ratio below 1, or increased values above 2.5) have been investigated in a few host-pathogen systems, and found to correlate with poor host survival prospects (McBride and Striker 2017). For instance, high values of the CD4^+^/CD8^+^ ratio have been shown to be predictors of patient mortality during SARS-CoV-2 infection (De Zunai et al. 2021; Pascual-Dapena et al. 2022). We found similar results, although the correlation between the CD4^+^/CD8^+^ ratio and mortality could only be assessed at the population (inter-group) level. The increase of the CD4^+^/CD8^+^ ratio probably reflects a global, temporary, dysfunction of the immune response, which favors the replication rate of Py in coinfected hosts. In agreement with this view, BALB/c mice infected with Py and depleted of CD8^+^ T cells suffered from greater parasitemia and higher mortality compared to control individuals during the period of peak parasitemia (Imai et al. 2015).

This work provides some relevant findings for the epidemiology of the coinfection between nematodes and *Plasmodium* and possibly for the evolution of virulence when parasites exploit the same host. In terms of the epidemiology of the disease, a key question is whether infection order is random or if it intrinsically depends on the life cycle of the parasite. In the specific case of the coinfection between gastrointestinal nematodes and *Plasmodium*, hosts might have higher chances to suffer from ordered infections where *Plasmodium* infects hosts already harboring worms. This suggestion is based on the observation that gastrointestinal nematodes usually establish chronic, steady state, infections that can last for months (or even years for some human helminths) while *Plasmodium* produces acute infections (lasting 2-4 weeks) that can either be cleared or lead to subsequent relapses (again depending on the parasite and the host species). However, this hypothesis rests on the assumption that hosts have equal probabilities to be infected by the two parasites per unit of time, or in other words that the probability of infection is not age-dependent. Actually, age-dependent infection might depend on the local epidemiological conditions. In areas of malaria hyperendemicity (high transmission rate and high R_0_), it is plausible that children are first infected with *Plasmodium* (Natama et al. 2018, Buchwald et al. 2019, Trape et al. 2024) and only at later ages by gastrointestinal nematodes. However, this pattern is likely to change in areas of low malaria transmission (low R_0_) where children might acquire helminth infection first (Blouin et al. 2018, Chis Ster et al. 2021), and still harbor them in case of subsequent *Plasmodium* infection. It should also be reminded that our experimental work only considered primary infections (reinfection was not allowed). However, in the real world, hosts are repeatedly exposed to multiple parasites and when there is no sterilizing immunity, previous infection does not fully prevent subsequent reinfection. The complexity due to age-dependent probability of infection and repeated exposure/infection might therefore blur the importance of the infection order as a driver of disease severity in natural settings.

Our results also suggest that the campaigns of deworming that have been implemented in several countries with the aim of reducing the negative effects of helminthiases on infant physical and cognitive development, might also have indirect benefits in terms of reduced severity of malaria symptoms, although the epidemiological surveys that have been published so far have reported been inconclusive results (Afolabi et al. 2021, Dila et al. 2022). However, we should keep in mind that interacting organisms exert reciprocal selection pressures and any prediction on the effect of deworming should also consider the microevolutionary response of the coinfecting pathogen. With this respect, our results might also be relevant to predict how selection might shape the optimal level of virulence (defined as infection-induced host damage) when parasites exploit hosts that harbor coinfecting parasites that elicit different immune effectors. A key prediction here is that species-specific cost of virulence might constraint the evolution of host damage in different ways in coinfecting parasites, and shape the optimal strategy of immunomodulation in species that rely on host immune suppression for long-term persistence. Experimental evolution and theoretical modelling would be valuable tools to explore these questions. A final thought is that, although we considered that the coinfection dynamics with *Plasmodium* and nematodes is regulated by top-down mechanisms involving the host immune response, we cannot ignore that both parasites (and nematodes in particular) can also interfere with the host immune response for their own benefit. This paves the way for a scenario where the top-down mechanisms might also be under parasite control.

## Conclusion

To conclude, our integrative approach provided several lines of evidence showing that the order of infection is an important factor shaping the severity of disease symptoms in nematode/malaria coinfected hosts. From a more mechanistic perspective, our results suggest that this order-dependent disease severity stems from the trade-off between specific immune effectors that provide protection against either nematodes or *Plasmodium*. We hope that these results will stimulate further research on both the importance of infection order under natural settings, and the potential epidemiological, and public health consequences of large-scale helminth removal through deworming campaigns.

## Acknowledgments

We are grateful to Valérie Saint Giorgio and all the staff of the animal facility for taking care of the animals.

## Author contribution

GS, BF, MR and BR conceived the study and gathered the funding; AD, LB, NP, GS and BF collected the data; AD, LB, EG performed the lab work; NP assisted with the spectral flow cytometry analysis; GS analyzed the data; GS, AD and LB wrote the first draft of the manuscript; all authors revised and approved the final version.

## Data availability statement

All the data and codes are available in DRYAD (DOI: 10.5061/dryad.0gb5mkmcb).

