## supplementary tables and figures for "Disease severity in coinfected hosts: the importance of infection order"

Table S1. Primer sequences used for the RT-qPCR.

| Target | Forward sequence (5' to 3') | Reverse sequence (5' to 3') |
|---|---|---|
| β-Actin | ATGGAGGGGAATACAGCCC | TTCTTTGCAGCTCCTTCGTT |
| GAPDH | CTCCCACTCTTCCACCTTCG | GCCTCTCTTGCTCAGTGTCC |
| Hmox-1 | GCCGAGAATGCTGAGTTCATG | TGGTACAAGGAAGCCATCACC |
| IFN-γ | GAGCTCATTGAATGCTTGGC | GCGTCATTGAATCACACCTG |
| IL-10 | AAGGCTTGGCAACCCAAGTAACCC | TGCACTACCAAAGCCACAAGGCAG |
| TGF-β1 | CCCAGTCTCCATACATTAACCC | CACCATCACCTTGAACTCTGAC |

Table S2. 27-color spectral flow cytometry panel used for the staining of mouse splenocytes.

| Emission (nm) | Target | Fluorophore | Clone | Supplier | Cat. ID |
|---|---|---|---|---|---|
| *Viability* | | | | | |
| 473 | / | LIVE/DEAD Blue | / | Invitrogen | L23105 |
| *Surface markers* | | | | | |
| 388 | CD45 | BUV395 | 30-F11 | BD Bioscience | 564279 |
| 496 | CD80 | BUV496 | 16-10A1 | BD Bioscience | 741091 |
| 563 | F4/80 | BUV563 | T45-2342 | BD Bioscience | 749284 |
| 615 | CD19 | BUV615 | 1D3 | BD Bioscience | 751213 |
| 661 | CD8α | BUV661 | 53-6.7 | BD Bioscience | 750023 |
| 737 | LAG-3 | BUV737 | C9B7W | BD Bioscience | 741820 |
| 805 | CD11b | BUV805 | M1/70 | BD Bioscience | 741934 |
| 421 | CTLA-4 | BV421 | UC10-4B9 | BioLegend | 106312 |
| 451 | Ly6G | Pacific Blue | 1A8 | BioLegend | 127612 |
| 506 | PD-1 | eFluor506 | J43 | ThermoFisher | 69-9985-82 |
| 570 | Ly6C | BV570 | HK1.4 | BioLegend | 128030 |
| 605 | CD3 | BV605 | 17A2 | BD Bioscience | 564009 |
| 650 | CD11c | BV650 | N418 | BioLegend | 117339 |
| 711 | CD127 | BV711 | A7R34 | BioLegend | 135035 |
| 750 | NkP46 | BV750 | 29A1.4 | BD Bioscience | 746875 |
| 786 | CD49d | BV786 | R1-2 | BD Bioscience | 564397 |
| 521 | CD163 | FITC | TNKUPJ | ThermoFisher | 11-1631-82 |
| 710 | PD-L1 | PerCP-eFluor 710 | MIH5 | ThermoFisher | 46-5982-82 |
| 660 | CD11a | APC | M17/4 | BioLegend | 101120 |
| 700 | CD4 | AF700 | RM4-5 | BD Bioscience | 557956 |
| 780 | MHCII | APC-Cy7 | M5/114.15.2 | BioLegend | 107628 |
| *Intracellular markers* | | | | | |
| 480 | ki67 | BV480 | B56 | BD Bioscience | 566109 |
| 576 | Tbet | PE | 4B10 | BD Bioscience | 561265 |
| 610 | RORγt | PE-CF594 | Q31-378 | BD Bioscience | 562684 |
| 693 | GATA3 | BB700 | L50-823 | BD Bioscience | 566642 |
| 774 | FOXP3 | PE-Cy7 | FJK-16s | Invitrogen | 25-5773-82 |

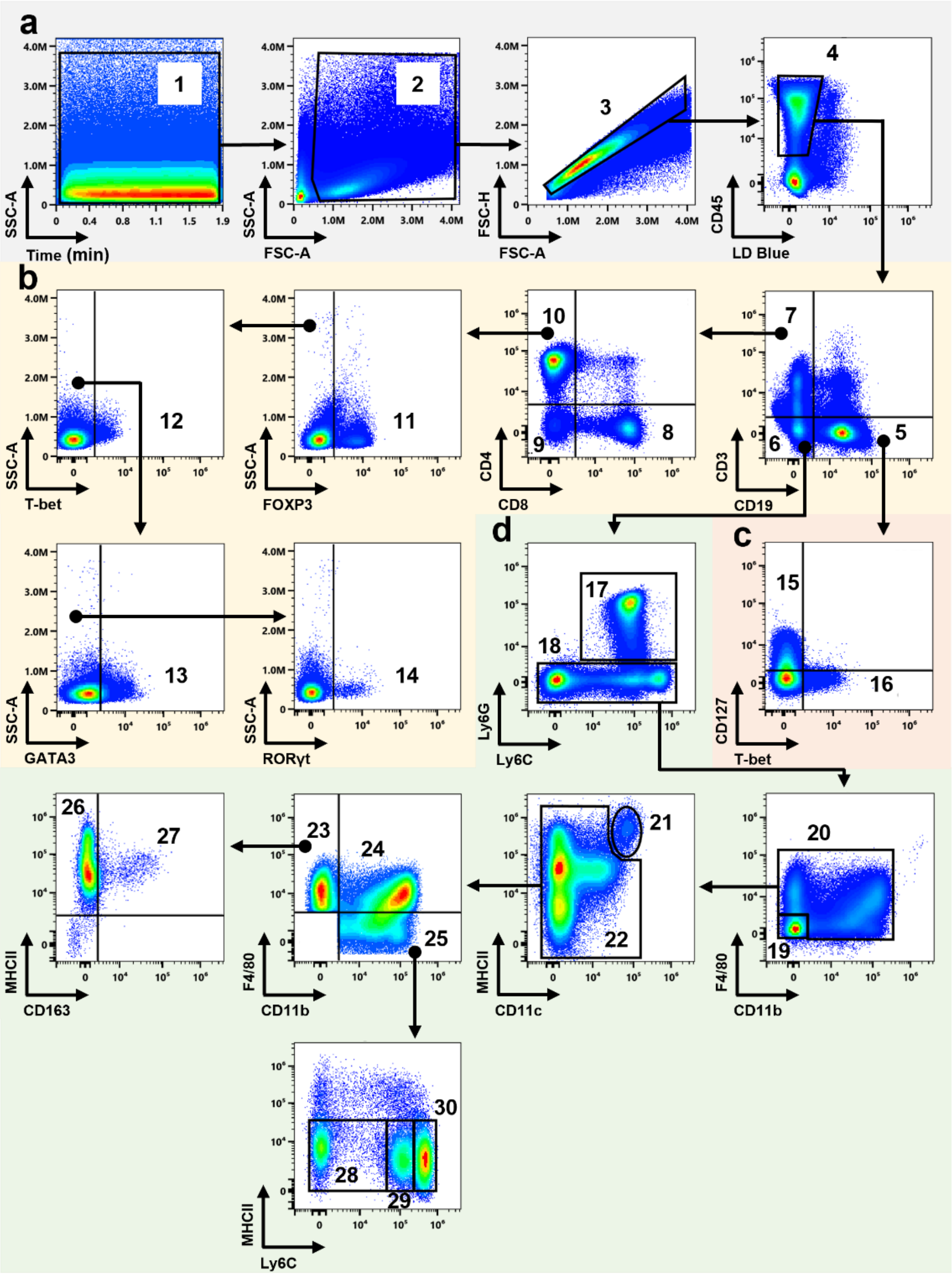


Figure S1. Gating strategy used for the spectral flow cytometry analysis to separate mouse splenocyte subsets. a) Gating of live immune cells. (1) Time parameter to exclude microfluidic fluctuations. Selection of (2) splenocytes without debris and (3) singlets. Selection of (4) live LD-Blue$^{-}$ and immune CD45$^{+}$ cells. b) Gating of T lymphocyte subsets. Selection of (5) CD19$^{+}$CD3$^{-}$ B cells, (6) LIN$^{-}$ cells and (7) CD19$^{-}$CD3$^{+}$ T cells. Selection of (8) CD8$^{+}$ T cells, (9) CD4$^{-}$CD8$^{-}$ other T cells and (10) CD4$^{+}$ T cells. Selection of (11) FoxP3$^{+}$ Treg, (12) Tbet$^{+}$ Th1, (13) GATA-3$^{+}$ Th2 and (14) RORγt$^{+}$ Th17 cell subsets within CD4$^{+}$ T lymphocytes. c) Gating of B lymphocyte subsets. Selection of (15) CD127$^{+}$ immature B cells and (16) Tbet$^{+}$ memory B cells. d) Gating of LIN$^{-}$ cells. Separation of (17) Ly6G$^{+}$ neutrophils from (18) other LIN$^{-}$ cells. Selection of (19) non-myeloid LIN$^{-}$ cells and (20) other myeloid cells. Separation of (21) MHCII$^{high}$CD11c$^{high}$ dendritic cells from (22) non-dendritic other myeloid cells. Selection of (23) F4/80$^{+}$CD11b$^{-}$ red pulp macrophages, (24) F4/80$^{+}$CD11b$^{+}$ macrophages and (25) F4/80$^{-/low}$CD11b$^{+}$ monocytes. Selection of (26) MHCII$^{+}$CD163$^{-}$ red pulp macrophages and (27) MHCII$^{+}$CD163$^{+}$ scavenger red pulp macrophages. Selection of (28) MHCII$^{-/low}$Ly6C$^{-/low}$, (29) MHCII$^{-/low}$Ly6C$^{int}$ and (30) MHCII$^{-/low}$Ly6C$^{high}$ monocyte subsets.

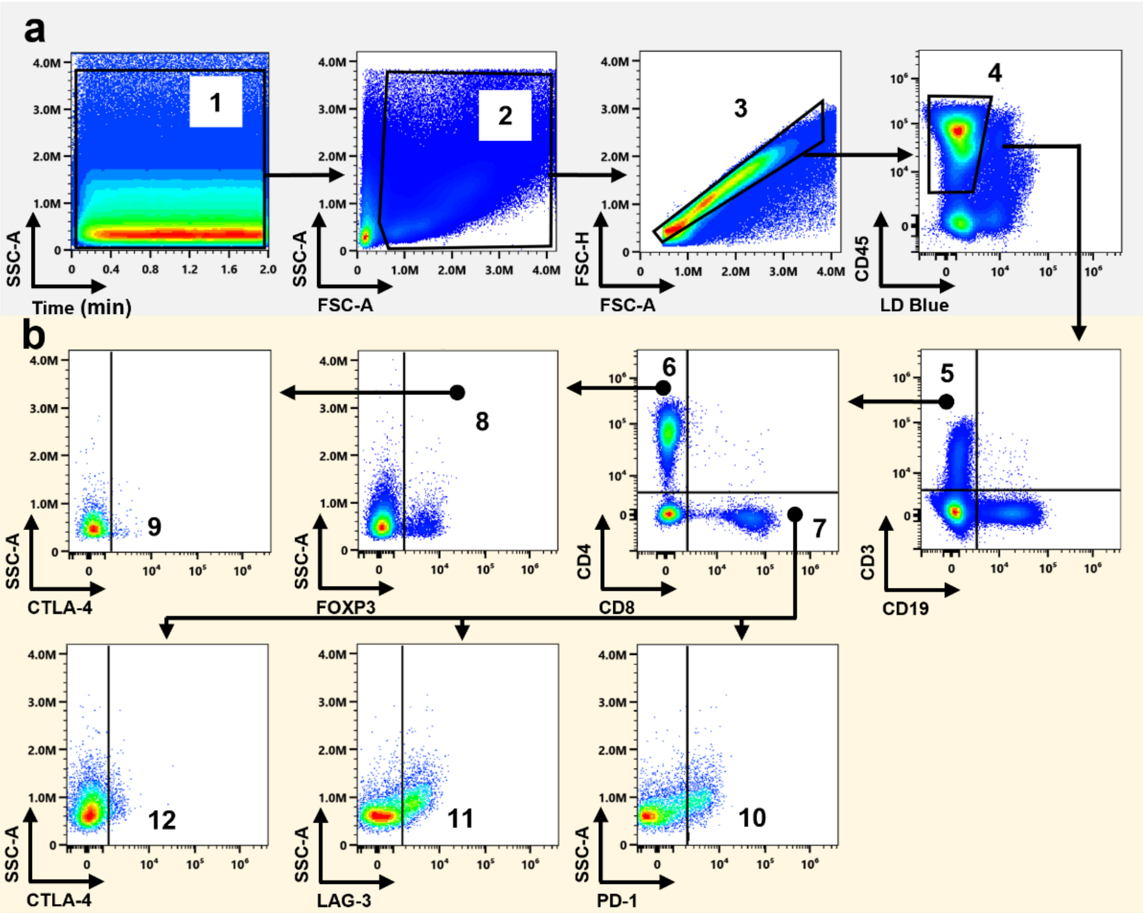


Figure S2. Gating strategy used for the spectral flow cytometry analysis to identify immune checkpoint markers within mouse splenocyte subsets. a) Gating of live immune cells. (1) Time parameter to exclude microfluidic fluctuations. Selection of (2) splenocytes without debris and (3) singlets. Selection of (4) live LD-Blue$^-$ and immune CD45$^+$ cells. b) Gating of T lymphocyte subsets. Selection of (5) CD19$^-$ CD3$^+$ T cells. Selection of (6) CD4$^+$ T cells and (7) CD8$^+$ T cells. Selection of (8) FoxP3$^+$ Treg. Identification of (9) CTLA-4$^+$ Treg cells. Identification of (10) PD-1$^{+,}$ (11) LAG-3$^+$ and (12) CTLA-4$^+$ cells from CD8$^+$ T lymphocyte subset.

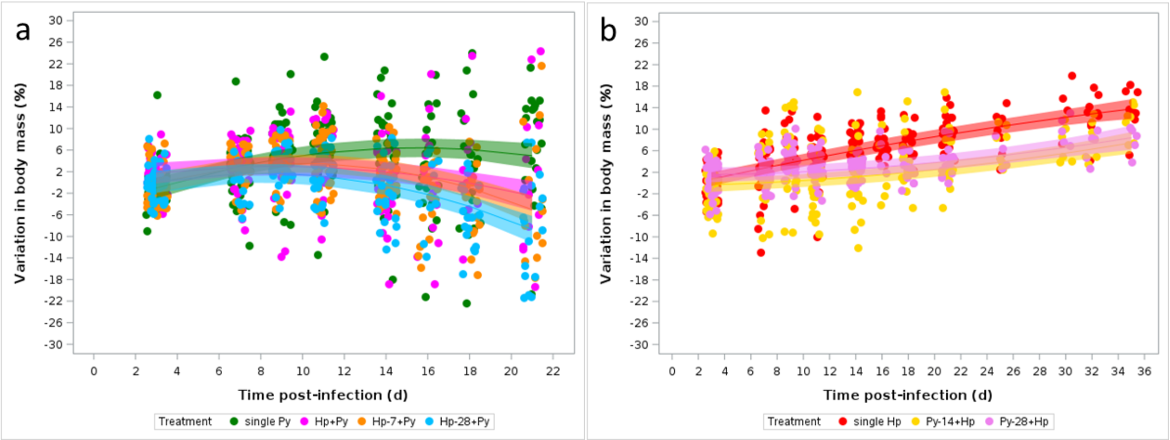


Figure S3. Changes in body mass over the course of the infection. Body mass is expressed as percent change with respect to body mass at day 0 for the single infection and the simultaneous infection or the day of the last infection for the coinfection groups. a) Changes in body mass for single Py infected mice and coinfected (Hp+Py, Hp-7+Py, Hp-28+Py) mice; b) Changes in body mass for single Hp infected mice and coinfected (Py-14+Hp, Py-28+Hp) mice. Dots represent the raw data, the lines represent the fit of a GLMM, and the shaded area around the line the 95% CI.

Table S3. General linear mixed model investigating the changes in body mass (percent change with respect to day 0) in single Py infected and coinfected (Hp+Py, Hp-7+Py, Hp-28+Py) mice. We report the parameter estimates with the 95% CI, df (degrees of freedom), F and p values. Mouse ID was included as a random effect to take into account the non-independence of observations for the same individual over time. N = 156 individuals and 789 observations.

| **Fixed effects** | | **Estimate** | **95% CI** | **df** | **F** | **p** |
|---|---|---|---|---|---|---|
| Treatment | | | | 3,625 | 2.32 | 0.0742 |
| | Py | 0 | | | | |
| | Hp+Py | 5.2 | 1.14/9.25 | | | |
| | Hp-7+Py | 1.28 | -2.74/5.3 | | | |
| | Hp-28+Py | 2.78 | -1.33/6.9 | | | |
| Time | | 0.544 | 0.35/0.74 | 1,625 | 73.10 | <0.0001 |
| Squared time | | -0.004 | -0.01/0.001 | 1,625 | 90.37 | <0.0001 |
| Time*Treatment | | | | 3,625 | 2.19 | 0.0882 |
| | Py | 0 | | | | |
| | Hp+Py | -0.84 | -1.58/-0.1 | | | |
| | Hp-7+Py | -0.04 | -0.76/0.68 | | | |
| | Hp-28+Py | -0.49 | -1.22/0.24 | | | |
| Squared Time*Treatment | | | | 3,625 | 1.70 | 0.1661 |
| | Py | 0 | | | | |
| | Hp+Py | 0.01 | -0.02/0.04 | | | |
| | Hp-7+Py | -0.02 | -0.05/0.007 | | | |
| | Hp-28+Py | -0.01 | 0.04/0.017 | | | |
| **Random effect** | | **Estimate** | **SE** | **z** | **p** | |
| ID | | 17.7 | 2.58 | 6.87 | <0.0001 | |

Table S4. General linear mixed model investigating the changes in body mass (percent change with respect to day 0 or the day of the last infection for the coinfection groups) in single Hp infected and coinfected (Py-14+Hp, Py-28+Hp) mice. We report the parameter estimates with the 95% CI, df (degrees of freedom), F and p values. Mouse ID was included as a random effect to take into account the non-independence of observations for the same individual over time. N = 115 individuals and 609 observations.

| **Fixed effects** | | **Estimate** | **95% CI** | **df** | **F** | **p** |
|---|---|---|---|---|---|---|
| Treatment | | | | 2,488 | 2.47 | 0.086 |
| | Hp | 0 | | | | |
| | Py-14+Hp | 0.37 | -1.79/2.53 | | | |
| | Py-28+Hp | 2.36 | 0.15/4.57 | | | |
| Time | | 0.55 | 0.40/0.70 | 1,488 | 16.88 | <0.0001 |
| Squared time | | -0.003 | -0.008/0.0004 | 1,488 | 2.86 | 0.0916 |
| Time*Treatment | | | | 2,488 | 13.93 | <0.0001 |
| | Hp | 0 | | | | |
| | Py-14+Hp | -0.50 | -0.73/-0.28 | | | |
| | Py-28+Hp | -0.54 | -0.78/-0.31 | | | |
| Squared Time*Treatment | | | | 2,488 | 5.41 | 0.0047 |
| | Hp | 0 | | | | |
| | Py-14+Hp | 0.009 | 0.003/0.015 | | | |
| | Py-28+Hp | 0.009 | 0.003/0.016 | | | |
| **Random effect** | | **Estimate** | **SE** | **z** | **p** | |
| ID | | 11.50 | 1.9 | 6.07 | <0.0001 | |

Table S5. General linear mixed model investigating the changes in body mass (percent change with respect to the day of the last infection) in coinfected mice depending on the order of infection (either Hp first or Py first). We report the parameter estimates with the 95% CI, df (degrees of freedom), F and p values. Mouse ID was included as a random effect to take into account the non-independence of observations for the same individual over time. N = 151 individuals and 720 observations

| **Fixed effects** | | **Estimate** | **95% CI** | **df** | **F** | **p** |
|---|---|---|---|---|---|---|
| Treatment | | | | 1,565 | 5.40 | 0.0205 |
| | Py first | 0 | | | | |
| | Hp first | -2.79 | -5.14/-0.43 | | | |
| Time | | 0.26 | -0.05/0.57 | 1,565 | 47.44 | <0.0001 |
| Squared time | | -0.005 | -0.018/0.009 | 1,565 | 60.10 | <0.0001 |
| Time*Treatment | | | | 1,565 | 20.63 | <0.0001 |
| | Py first | 0 | | | | |
| | Hp first | 1.0 | 0.57/1.43 | | | |
| Squared Time*Treatment | | | | 1,565 | 45.74 | <0.0001 |
| | Py first | 0 | | | | |
| | Hp first | -0.06 | -0.08/-0.05 | | | |
| **Random effect** | | **Estimate** | **SE** | **z** | **p** | |
| ID | | 10.42 | 1.67 | 6.24 | <0.0001 | |

Table S6. General linear model investigating the changes in spleen mass (log mg) in non-infected, and single (Hp, Py) infected mice. We report the type III sum of squares, degrees of freedom (df), the F and p values. The $R^2$ of the model is 0.92. N = 117 individuals.

| **Source of variation** | **Type III SS** | **df** | **F** | **p** |
|---|---|---|---|---|
| Treatment | 9.02 | 2,105 | 328.25 | <0.0001 |
| Time | 4.32 | 3,105 | 104.82 | <0.0001 |
| Treatment*Time | 3.92 | 6,105 | 47.49 | <0.0001 |

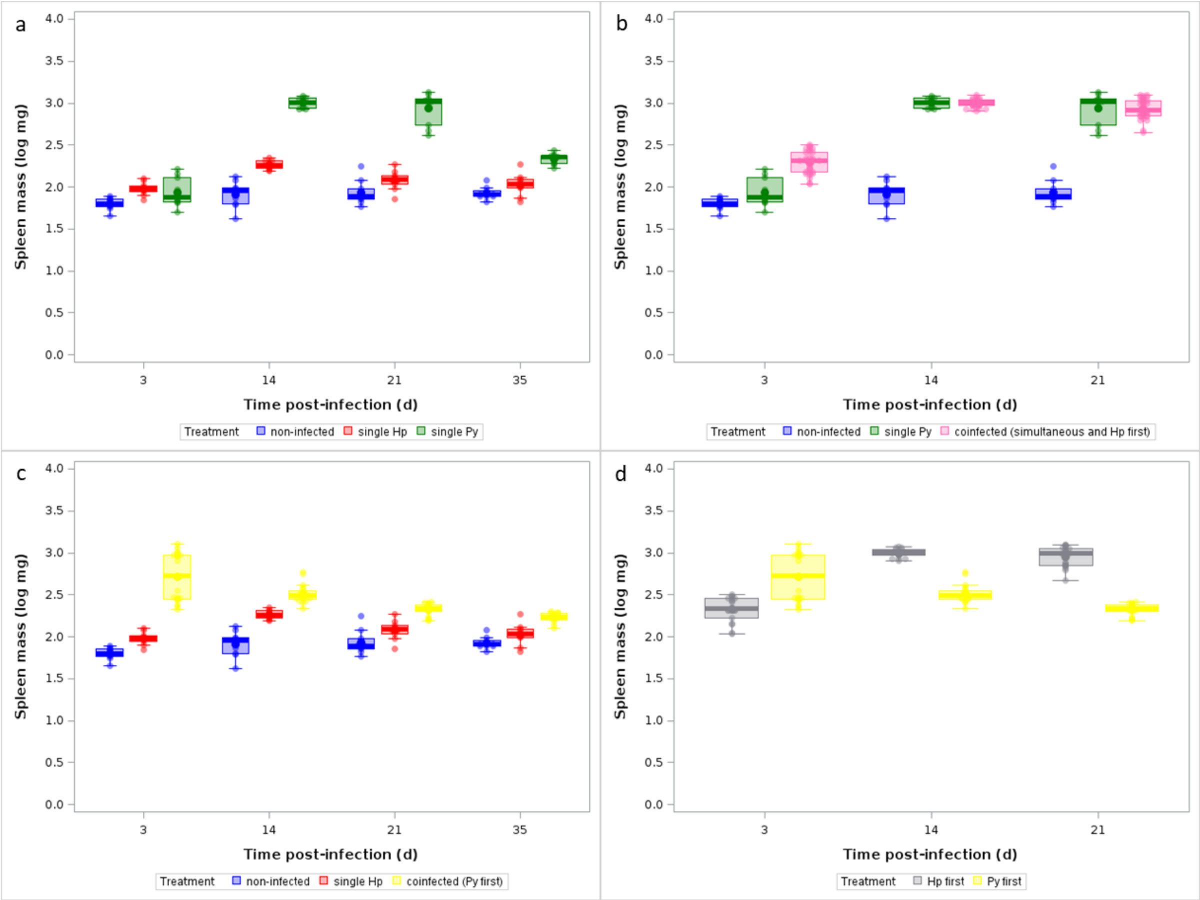


Figure S4. Changes in $log_{10}$ transformed spleen mass (mg) over the course of the infection. a) Changes in spleen mass for non-infected and single (Hp, Py) infected mice; b) Changes in spleen mass for non-infected, single Py infected and coinfected (simultaneous infection and Hp first) groups; c) Changes in spleen mass for non-infected, single Hp infected and coinfected (Py first) groups; d) Changes in spleen mass for coinfected groups depending on the order of the infection (Hp infecting first or Py infecting first). Dots represent the raw data, the boxes represent the interquartile range (IQR), the horizontal lines the median, and whiskers the range of data within 1.5 the IQR.

Table S7. General linear model investigating the changes in spleen mass (log mg) in single Py infected and coinfected (Hp+Py, Hp-7+Py, Hp-28+Py) mice. We report the type III sum of squares, degrees of freedom (df), the F and p values. The $R^2$ of the model is 0.92. N = 118 individuals.

| **Source of variation** | **Type III SS** | **df** | **F** | **p** |
|---|---|---|---|---|
| Treatment | 0.49 | 3,106 | 12.47 | <0.0001 |
| Time | 15.08 | 2,106 | 580.71 | <0.0001 |
| Treatment*Time | 0.76 | 6,106 | 9.73 | <0.0001 |

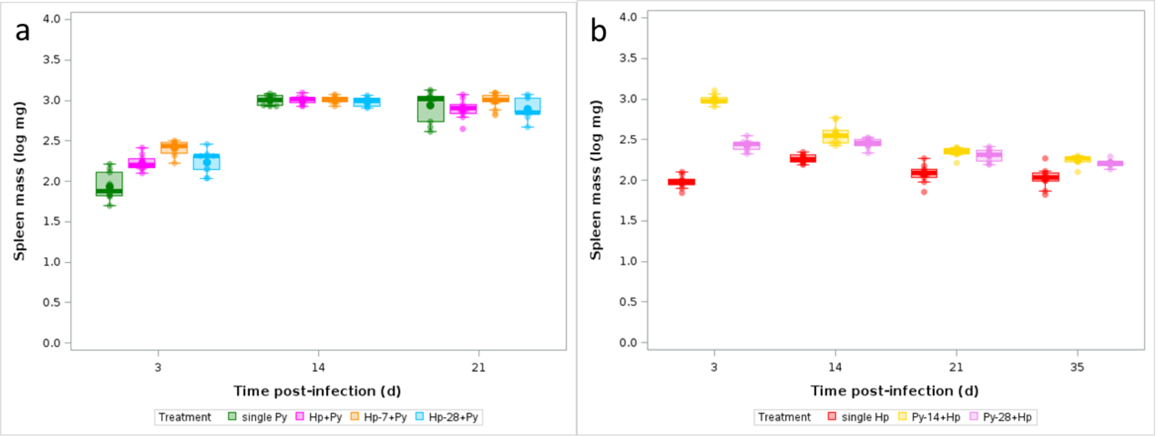


Figure S5. Changes in $log_{10}$ transformed spleen mass (mg) over the course of the infection (for the coinfection groups, time p.i. refers to the time post last infection); a) Changes in spleen mass for single Py infected mice and coinfected (Hp+Py, Hp-7+Py, Hp-28+Py) mice; b) Changes in spleen mass for single Hp infected and coinfected (Py-14+Hp, Py-28+Hp) mice. Dots represent the raw data, the boxes represent the interquartile range (IQR), the horizontal lines the median, and whiskers the range of data within 1.5 the IQR.

Table S8. General linear model investigating the changes in spleen mass (log mg) in non-infected, single Py infected and coinfected (simultaneous infection and Hp first) mice. We report the type III sum of squares, degrees of freedom (df), the F and p values. The $R^2$ of the model is 0.94. N = 148 individuals.

| **Source of variation** | **Type III SS** | **df** | **F** | **P** |
|---|---|---|---|---|
| Treatment | 16.74 | 2,139 | 558.07 | <0.0001 |
| Time | 9.49 | 2,139 | 316.5 | <0.0001 |
| Time* Treatment | 2.86 | 4,139 | 47.65 | <0.0001 |

Table S9. General linear model investigating the changes in spleen mass (log mg) in single Hp infected and coinfected (Py-14+Hp, Py-28+Hp) mice. We report the type III sum of squares, degrees of freedom (df), the F and p values. The $R^2$ of the model is 0.92. N = 108 individuals.

| **Source of variation** | **Type III SS** | **df** | **F** | **p** |
|---|---|---|---|---|
| Treatment | 3.66 | 2,96 | 259.23 | <0.0001 |
| Time | 1.61 | 3,96 | 75.93 | <0.0001 |
| Time*Treatment | 2.13 | 6,96 | 50.44 | <0.0001 |

Table S10. General linear model investigating the changes in spleen mass (log mg) in non-infected, single Hp infected and coinfected (Py first) mice. We report the type III sum of squares, degrees of freedom (df), the F and p values. The $R^2$ of the model is 0.82. N = 148 individuals.

| **Source of variation** | **Type III SS** | **df** | **F** | **p** |
|---|---|---|---|---|
| Treatment | 8.26 | 2,136 | 202.54 | <0.0001 |
| Time | 0.51 | 3,136 | 8.42 | <0.0001 |
| Time*Treatment | 2.00 | 6,136 | 16.36 | <0.0001 |

Table S11. General linear model investigating the changes in spleen mass (log mg) in coinfected mice depending on the order of infection (Hp first or Py first). We report the type III sum of squares, degrees of freedom (df), the F and p values. The $R^2$ of the model is 0.76. N = 117 individuals.

| **Source of variation** | **Type III SS** | **df** | **F** | **p** |
|---|---|---|---|---|
| Treatment | 1.69 | 1,111 | 70.43 | <0.0001 |
| Time | 1.04 | 2,111 | 21.60 | <0.0001 |
| Time*Treatment | 5.87 | 2,111 | 122.29 | <0.0001 |

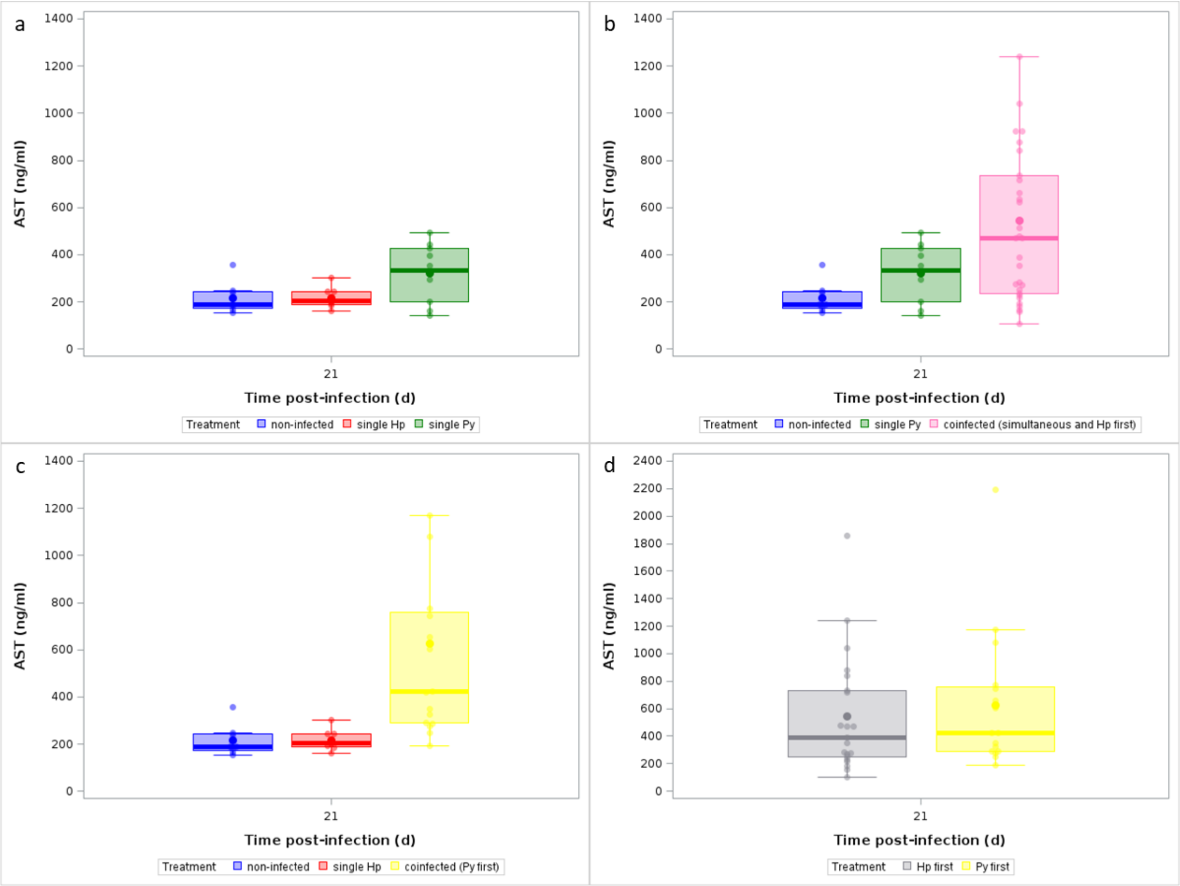


Figure S6. Effect of infection and coinfection on liver damage. a) Levels of aspartate aminotransferase (AST, ng/ml) in the plasma at day 21 p.i. in non-infected and single (Hp, Py) infected mice; b) AST level in plasma of non-infected, single Py infected and coinfected (simultaneous infection and Hp first) groups at day 21 p.i.; c) AST level in plasma of non-infected, single Hp infected and coinfected (Py first) groups at day 21 p.i.; d) AST level in plasma of coinfected mice, depending on the order of the infection (Hp infecting first or Py infecting first) at day 21 post-infection. Dots represent the raw data, the boxes represent the interquartile range (IQR), the horizontal lines the median, and whiskers the range of data within 1.5 the IQR.

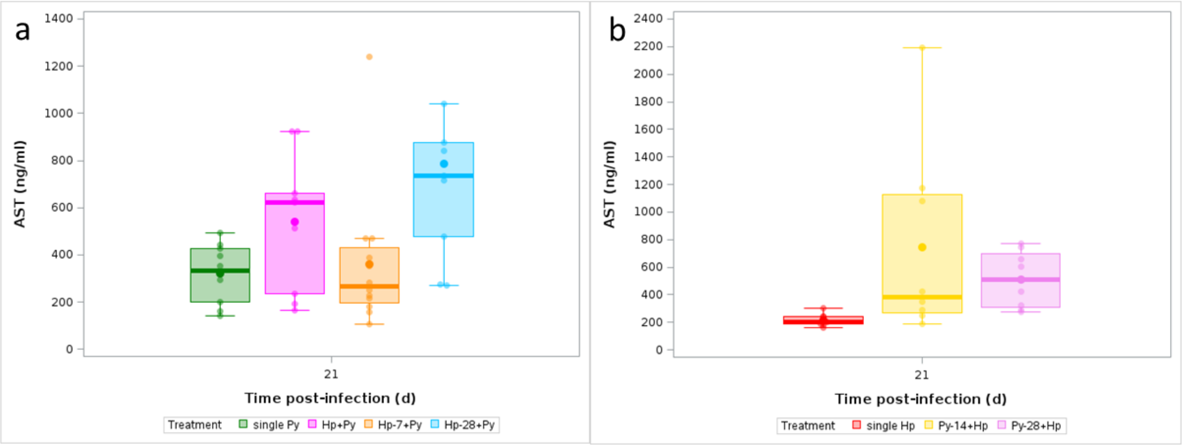


Figure S7. a) Levels of aspartate aminotransferase (AST, ng/ml) in the plasma at day 21 p.i. in single Py infected and coinfected (Hp+Py, Hp-7+Py, Hp-28+Py) mice; b) AST level in plasma of single Hp infected and coinfected (Py-14+Hp, Py-28+Hp) mice. Dots represent the raw data, the boxes represent the interquartile range (IQR), the horizontal lines the median, and whiskers the range of data within 1.5 the IQR.

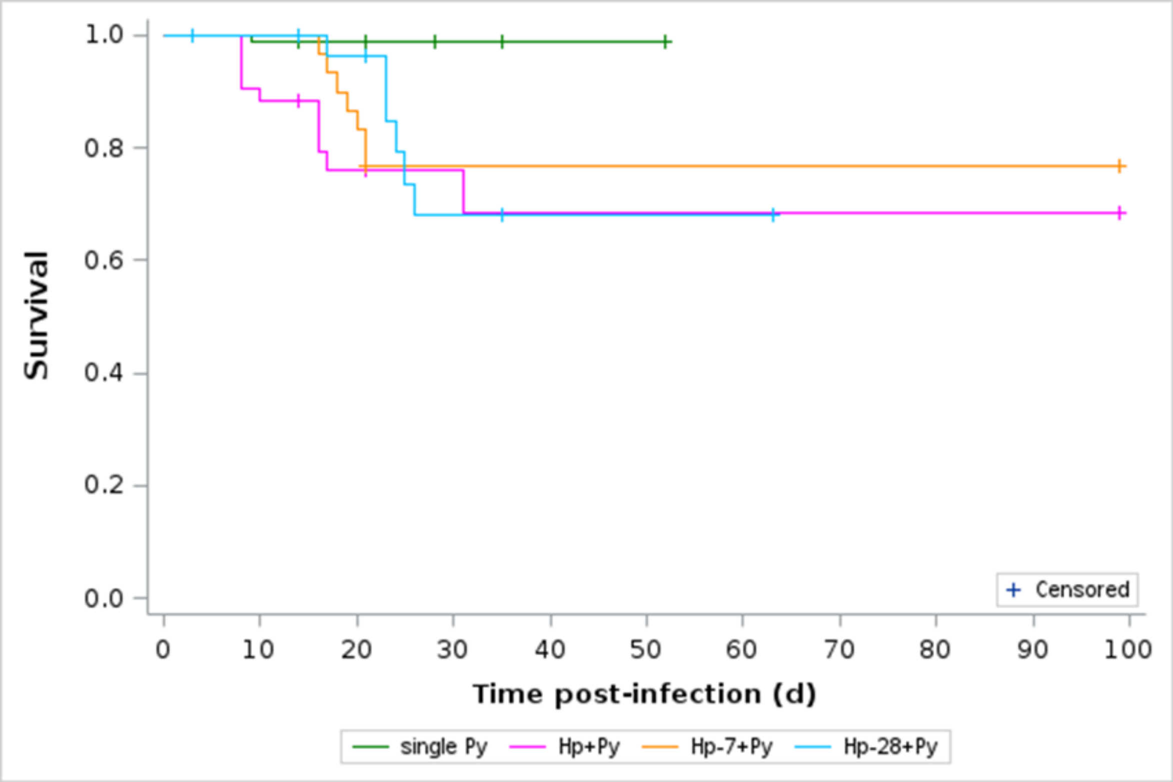


Figure S8. Survival curves for single Py infected mice and coinfected (Hp+Py, Hp-7+Py, Hp-28+Py) mice. Crosses indicate censored observations.

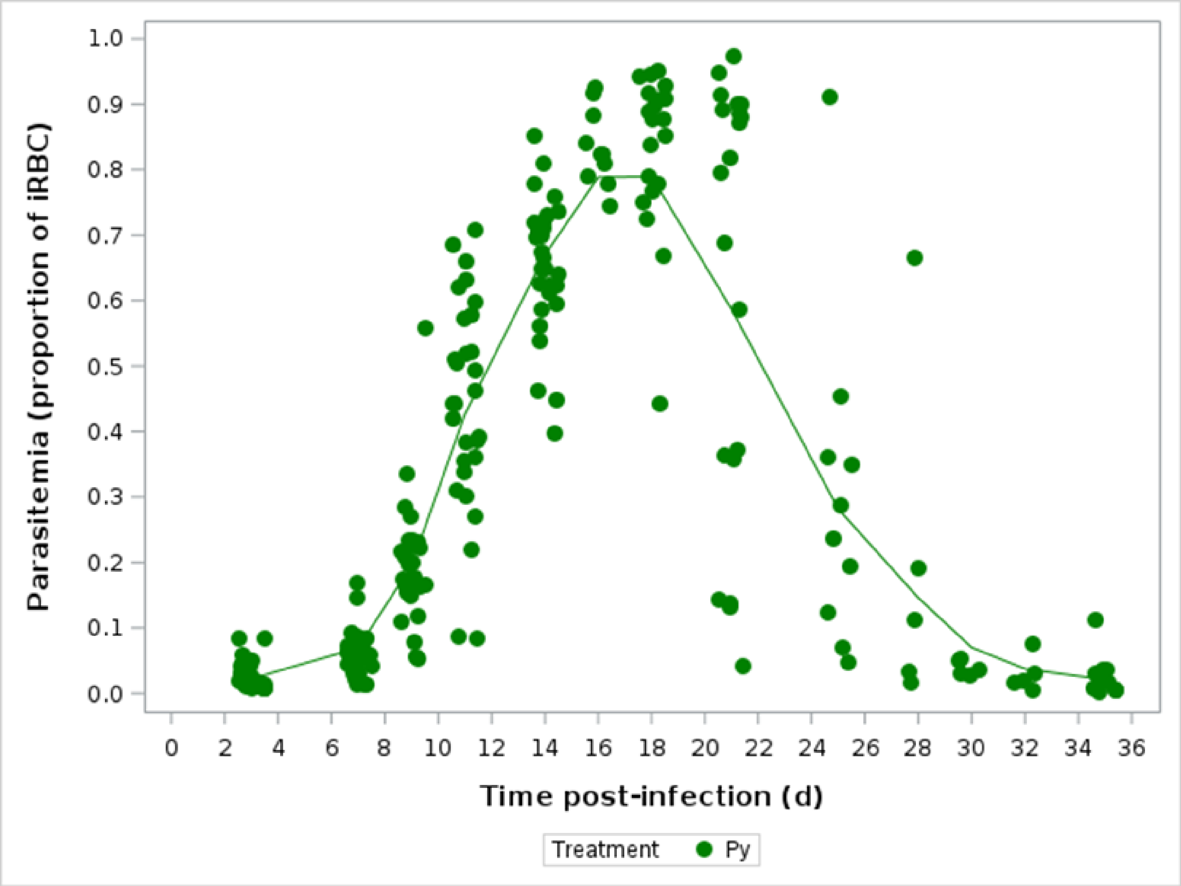


Figure S9. Changes in parasitemia (proportion of iRBCs) in single Py infected mice over time. Dots represent the raw data, the line corresponds to the fit of a GAM where the level of smoothing was determined based on the GCV criterion.

Table S12. General linear mixed model investigating the changes in parasitemia (proportion of iRBCs) in single Py infection and coinfected (Hp+Py, Hp-7+Py, Hp-28+Py) mice. We report the parameter estimates with the 95% CI, df (degrees of freedom), F and p values. Mouse ID was included as a random effect to take into account the non-independence of observations for the same individual over time. The distribution of errors was modelled using a beta distribution. N = 151 individuals and 665 observations.

| **Fixed effects** | | **Estimate** | **95% CI** | **df** | **F** | **p** |
|---|---|---|---|---|---|---|
| Treatment | | | | 3,506 | 8.22 | <0.0001 |
| | Py | 0 | | | | |
| | Hp+Py | 0.94 | -0.44/2.32 | | | |
| | Hp-7+Py | 2.91 | 1.66/4.16 | | | |
| | Hp-28+Py | 0.96 | -0.64/2.57 | | | |
| Time | | 1.05 | 0.9/1.2 | 1,506 | 486.96 | <0.0001 |
| Squared time | | -0.03 | -0.04/0.02 | 1,506 | 226.59 | <0.0001 |
| Time*Treatment | | | | 3,506 | 6.48 | 0.0003 |
| | Py | 0 | | | | |
| | Hp+Py | -0.13 | -0.34/0.08 | | | |
| | Hp-7+Py | -0.42 | -0.61/-0.22 | | | |
| | Hp-28+Py | -0.22 | -0.46/0.01 | | | |
| Squared Time*Treatment | | | | 3,506 | 7.51 | <0.0001 |
| | Py | 0 | | | | |
| | Hp+Py | 0.006 | -0.001/0.013 | | | |
| | Hp-7+Py | 0.02 | 0.01/0.024 | | | |
| | Hp-28+Py | 0.012 | 0.003/0.02 | | | |
| **Random effect** | | **Estimate** | **SE** | **z** | **p** | |
| ID | | 0.128 | 0.031 | 4.19 | <0.0001 | |

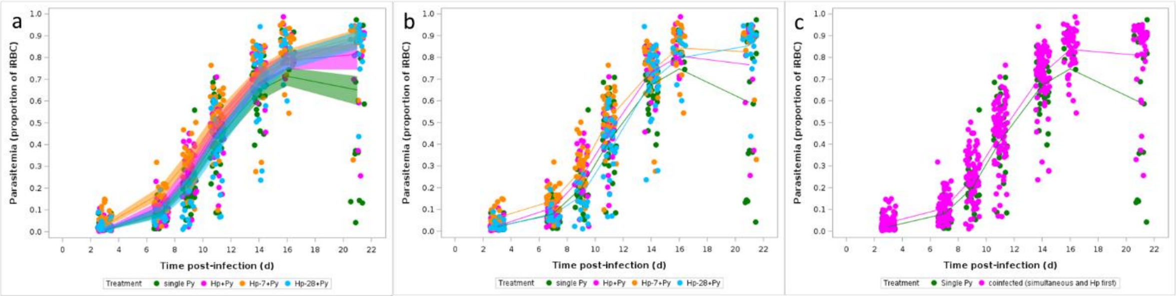


Figure S10. a) Changes in parasitemia (proportion of iRBCs) in single Py infected mice and coinfected (Hp+Py, Hp-7+Py, Hp-28+Py) mice over time. Dots represent the raw data, the lines represent the fit of the GLMM, and the shaded area around the line the 95% CI; b) As in a) but the lines represent the fit of a GAM where the level of smoothing was determined based on the GCV criterion; c) Changes in parasitemia (proportion of iRBCs) in single Py infected and coinfected (simultaneous infection and Hp first) mice. Dots represent the raw data, the lines correspond to the fit of a GAM where the level of smoothing was determined based on the GCV criterion.

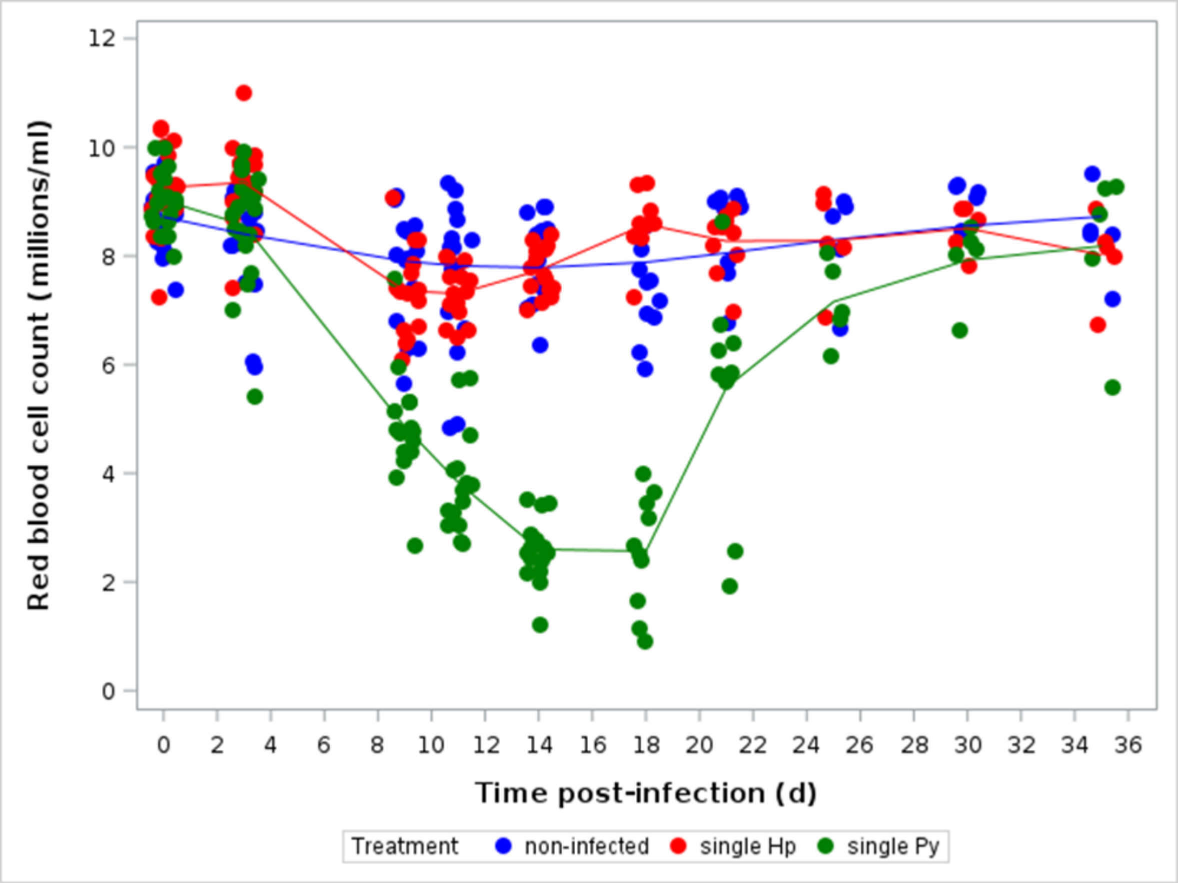


Figure S11. Changes in RBC counts ($10^6/\mu l$) over the course of the infection for non-infected and single (Hp, Py) infected mice. Dots represent the raw data, the lines correspond to the fit of a GAM where the level of smoothing was determined based on the GCV criterion.

Table S13. General linear mixed model investigating the changes in RBC counts ($10^6$/µl) in single Py infected and coinfected (Hp+Py, Hp-7+Py, Hp-28+Py) mice. We report the parameter estimates with the 95% CI, df (degrees of freedom), F and p values. Mouse ID was included as a random effect to take into account the non-independence of observations for the same individual over time. N = 74 individuals and 371 observations.

| **Fixed effects** | | **Estimate** | **95% CI** | **df** | **F** | **P** |
|---|---|---|---|---|---|---|
| Treatment | | | | 3,285 | 13.23 | <0.0001 |
| | Py | 0 | | | | |
| | Hp+Py | 0.313 | -0.365/0.991 | | | |
| | Hp-7+Py | -1.56 | -2.19/-0.93 | | | |
| | Hp-28+Py | -0.028 | -0.661/0.606 | | | |
| Time | | 0.092 | -0.116/0.301 | 1,285 | 0.34 | 0.5613 |
| Squared time | | -0.096 | -0.121/-0.07 | 1,285 | 99.84 | <0.0001 |
| Cubic time | | 0.004 | 0.003/0.005 | 1,285 | 162.67 | <0.0001 |
| Time*Treatment | | | | 3,285 | 3.76 | 0.0113 |
| | Py | 0 | | | | |
| | Hp+Py | -0.487 | -0.821/-0.154 | | | |
| | Hp-7+Py | 0.038 | -0.26/0.336 | | | |
| | Hp-28+Py | -0.052 | -0.349/0.246 | | | |
| Squared Time*Treatment | | | | 3,285 | 2.91 | 0.0351 |
| | Py | 0 | | | | |
| | Hp+Py | 0.061 | 0.02/0.103 | | | |
| | Hp-7+Py | 0.017 | -0.019/0.054 | | | |
| | Hp-28+Py | 0.021 | -0.015/0.058 | | | |
| Cubic Time*Treatment | | | | 3,285 | 3.57 | 0.0146 |
| | Py | 0 | | | | |
| | Hp+Py | -0.002 | -0.003/-0.0008 | | | |
| | Hp-7+Py | -0.001 | -0.002/0.0002 | | | |
| | Hp-28+Py | -0.001 | -0.002/-0.0001 | | | |
| **Random effect** | | **Estimate** | **SE** | **z** | **p** | |
| ID | | 0.302 | 0.078 | 3.85 | <0.0001 | |

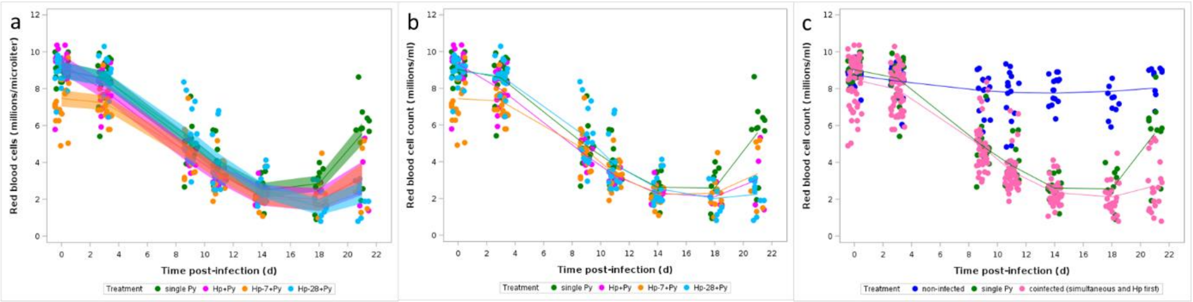


Figure S12. Changes in RBC counts ($10^6$/µl) over the course of the infection. a) Changes in RBCs for single Py infected and coinfected (Hp+Py, Hp-7+Py, Hp-28+Py) mice. Dots represent the raw data, the lines represent the fit of the GLMM, and the shaded area around the line the 95% CI; b) Same as a) but the lines correspond to the fit of a GAM where the level of smoothing was determined based on the GCV criterion; c) Changes in RBCs for non-infected, single Py infected and coinfected (simultaneous infection and Hp first) groups. Dots represent the raw data, the lines correspond to the fit of a GAM where the level of smoothing was determined based on the GCV criterion.

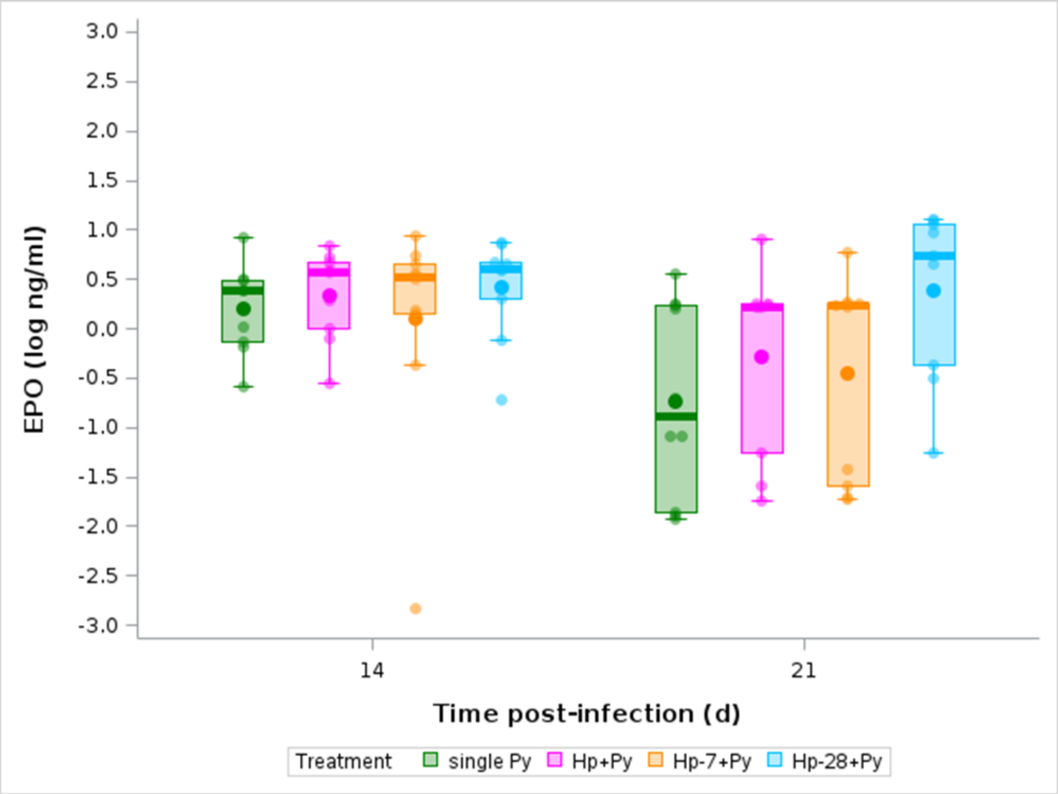


Figure S13. Levels of erythropoietin (EPO, log ng/ml) in the plasma at day 14 and 21 p.i. in single Py infected and coinfected (Hp+Py, Hp-7+Py, Hp-28+Py) mice. Dots represent the raw data, the boxes represent the interquartile range (IQR), the horizontal lines the median, and whiskers the range of data within 1.5 the IQR.

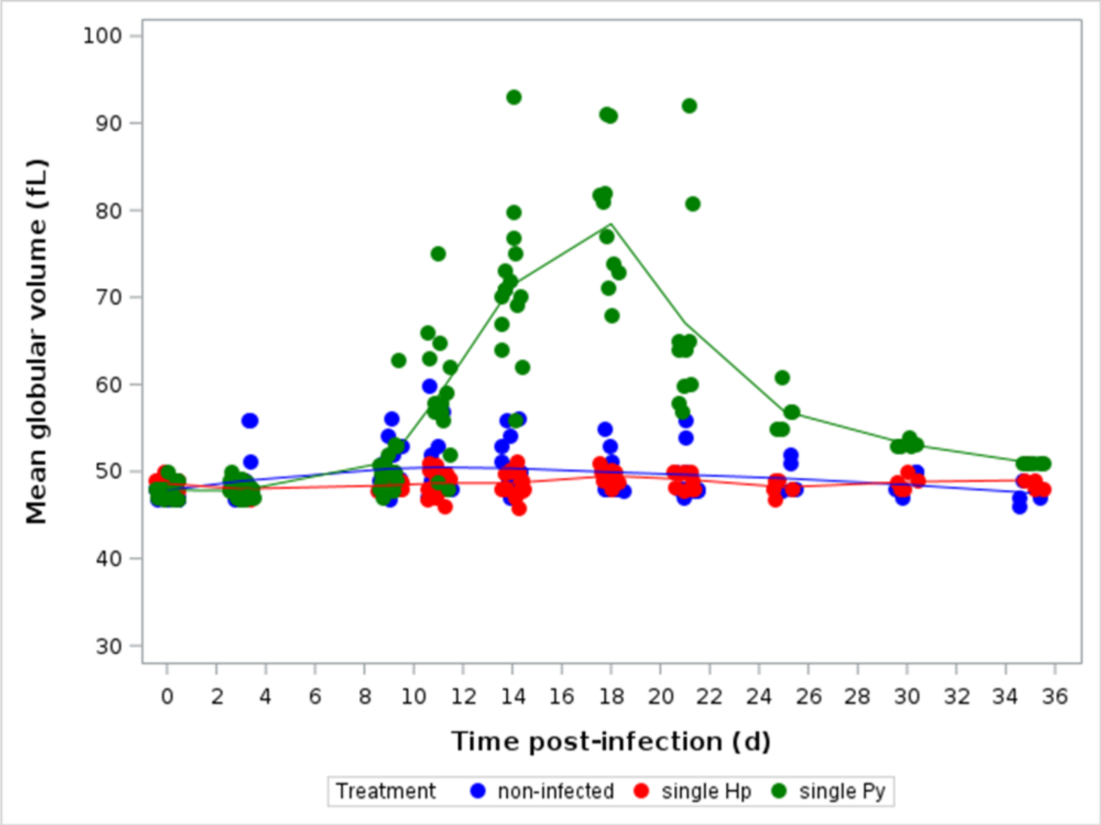


Figure S14. Changes in mean globular volume (fL) over the course of the infection for non-infected and single (Hp, Py) infected mice. Dots represent the raw data, the lines correspond to the fit of a GAM where the level of smoothing was determined based on the GCV criterion.

Table S14. General linear mixed model investigating the changes in mean globular volume (fL) in single Py infected and coinfected (Hp+Py, Hp-7+Py Hp-28+Py) mice. We report the parameter estimates with the 95% CI, df (degrees of freedom), F and p values. Mouse ID was included as a random effect to take into account the non-independence of observations for the same individual over time. N = 74 individuals and 371 observations.

| **Fixed effects** | | **Estimate** | **95% CI** | **df** | **F** | **p** |
|---|---|---|---|---|---|---|
| Treatment | | | | 3,285 | 0.41 | 0.7431 |
| | Py | 0 | | | | |
| | Hp+Py | -0.302 | -4.06/3.46 | | | |
| | Hp-7+Py | -1.40 | -4.86/2.07 | | | |
| | Hp-28+Py | 0.51 | -2.99/4.02 | | | |
| Time | | -3.75 | -5.02/-2.49 | 1,285 | 80.61 | <0.0001 |
| Squared time | | 0.67 | 0.51/0.82 | 1,285 | 173.66 | <0.0001 |
| Cubic time | | -0.021 | -0.026/-0.016 | 1,285 | 159.58 | <0.0001 |
| Time*Treatment | | | | 3,285 | 0.99 | 0.4000 |
| | Py | 0 | | | | |
| | Hp+Py | 1.50 | -0.52/3.53 | | | |
| | Hp-7+Py | 0.97 | -0.84/2.77 | | | |
| | Hp-28+Py | 0.13 | -1.68/1.94 | | | |
| Squared Time*Treatment | | | | 3,285 | 1.09 | 0.3519 |
| | Py | 0 | | | | |
| | Hp+Py | -0.22 | -0.47/0.03 | | | |
| | Hp-7+Py | -0.13 | -0.35/0.096 | | | |
| | Hp-28+Py | -0.067 | -0.29/0.15 | | | |
| Cubic Time*Treatment | | | | 3,285 | 1.18 | 0.3162 |
| | Py | 0 | | | | |
| | Hp+Py | 0.007 | -0.001/0.015 | | | |
| | Hp-7+Py | 0.004 | -0.002/0.012 | | | |
| | Hp-28+Py | 0.004 | -0.003/0.011 | | | |
| **Random effect** | | **Estimate** | **SE** | **z** | **p** | |
| ID | | 4.9 | 1.68 | 2.91 | 0.0018 | |

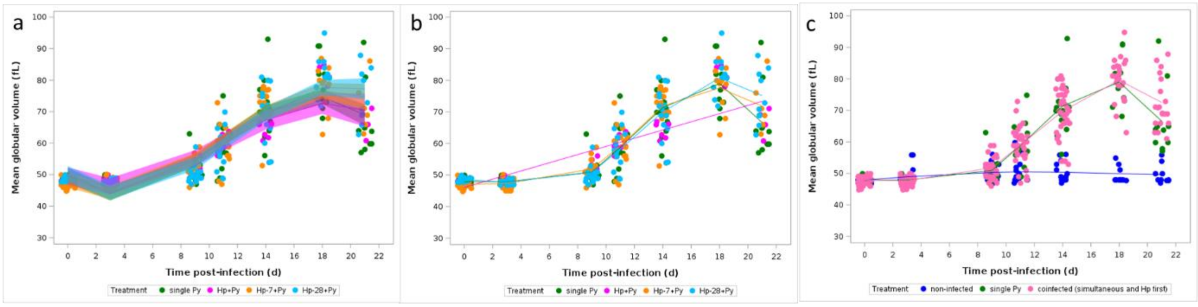


Figure S15. Changes in mean globular volume (MGV, fL) over the course of the infection. a) Changes in MVG for single Py infected mice and coinfected (Hp+Py, Hp-7+Py, Hp-28+Py) mice. Dots represent the raw data, the lines represent the fit of the GLMM, and the shaded area around the line the 95% CI; b) Same as a) but the lines correspond to the fit of a GAM where the level of smoothing was determined based on the GCV criterion; c) Changes in MVG for single Py infected and coinfected (simultaneous infection and Hp first) groups. Dots represent the raw data, the lines correspond to the fit of a GAM where the level of smoothing was determined based on the GCV criterion.

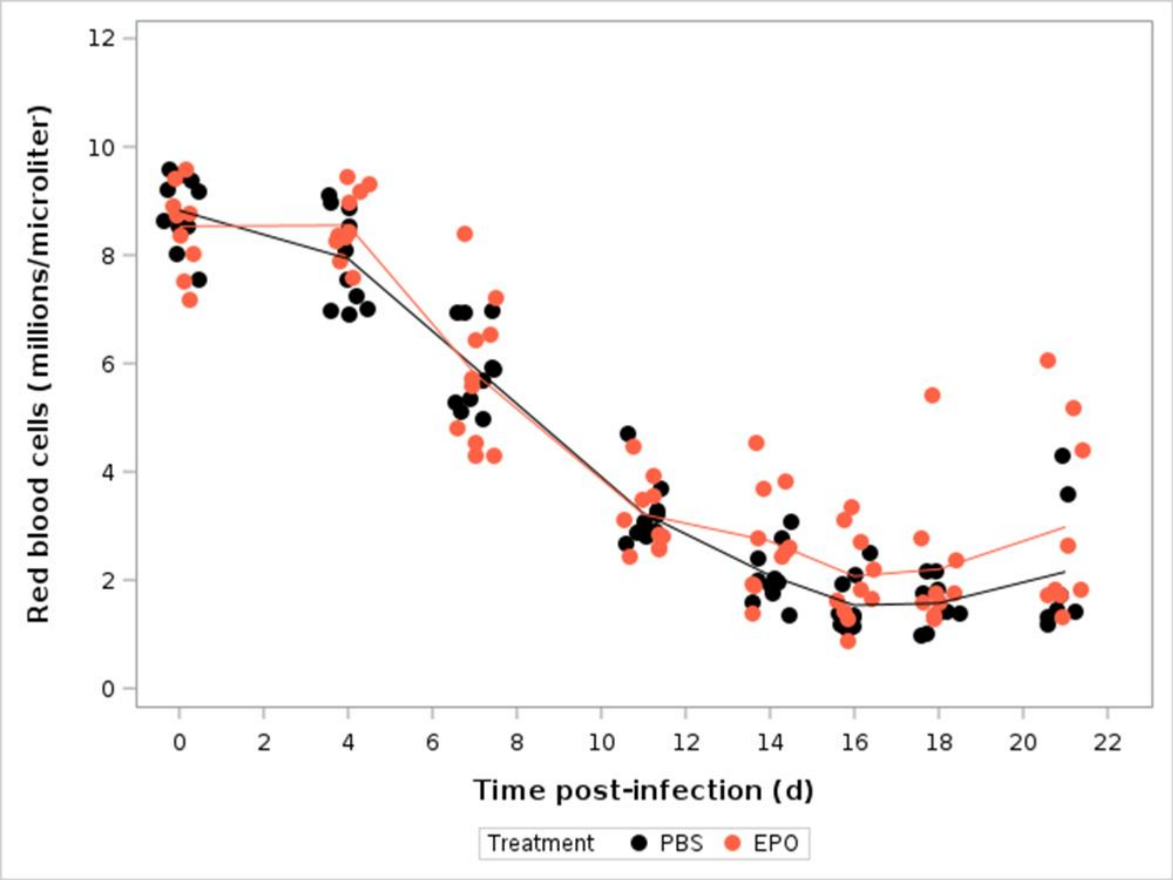


Figure S16. Changes in RBCs ($10^6/\mu l$) in coinfected (Hp-28+Py) mice treated with rEPO or left as control (PBS). Dots represent the raw data, the lines correspond to the fit of a GAM where the level of smoothing was determined based on the GCV criterion.

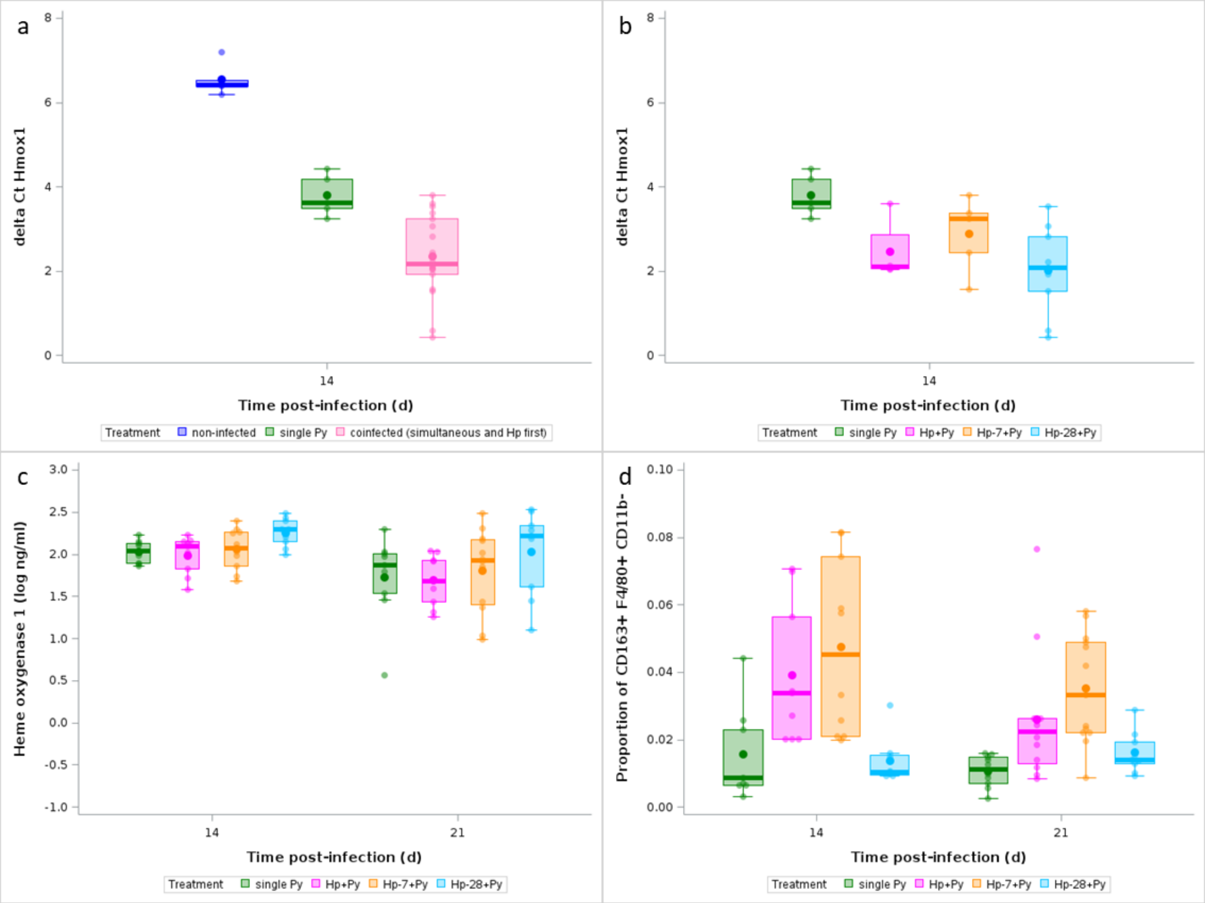


Figure S17. a) Gene expression levels relative to β-actin of *Hmox1* in splenocytes (the lower the delta Ct values the higher the expression) at day 14 p.i. in non-infected, single Py infected and coinfected (simultaneous infection and Hp first) mice; b) Expression levels of *Hmox1* in splenocytes (the lower the delta Ct values the higher the expression) at day 14 p.i. in single Py infected and coinfected (Hp+Py, Hp-7+Py, Hp-28+Py) mice; c) Levels of Heme oxygenase-1 in the plasma (log ng/ml) at day 14 and 21 p.i. in single Py infected and coinfected (Hp+Py, Hp-7+Py, Hp-28+Py) mice; d) proportion of CD163$^{+}$ F4/80$^{+}$ CD11b$^{-}$ at day 14 and 21 p.i. in single Py infected and coinfected (Hp+Py, Hp-7+Py, Hp-28+Py) mice. Dots represent the raw data, the boxes represent the interquartile range (IQR), the horizontal lines the median, and whiskers the range of data within 1.5 the IQR.

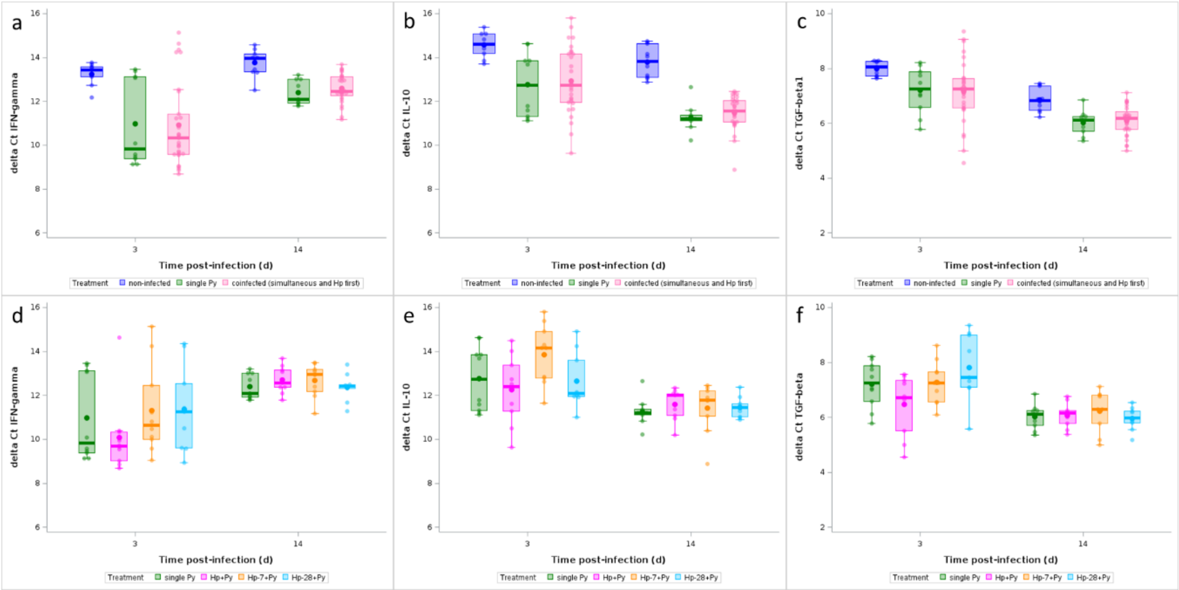


Figure S18. Gene expression levels relative to β-actin of a) IFN-γ, b) IL-10, c) TGF-β1 in splenocytes (the lower the delta Ct values the higher the expression) at day 3 and 14 p.i. in non-infected, single Py infected and coinfected (simultaneous infection and Hp first) mice. Gene expression levels of d) IFN-γ, e) IL-10, f) TGF-β1 in splenocytes (the lower the delta Ct values the higher the expression) at day 3 and 14 p.i. in single Py infected and coinfected (Hp+Py, Hp-7+Py, Hp-28+Py) mice. Dots represent the raw data, the boxes represent the interquartile range (IQR), the horizontal lines the median, and whiskers the range of data within 1.5 the IQR.

Table S15. Generalized linear model with a beta distribution of errors investigating the changes in the proportion of $FoxP3^{+}$ $CD4^{+}$ T cells in single Py infected, and coinfected (Hp+Py, Hp-7+Py, Hp-28+Py) mice at day 14 and 21 post-infection. We report the degrees of freedom (df), and the F and p values. N = 80 individuals.

| **Source of variation** | **df** | **F** | **p** |
|---|---|---|---|
| Treatment | 3,72 | 1.50 | 0.2220 |
| Time | 1,72 | 0.07 | 0.7865 |
| Treatment*Time | 3,72 | 0.21 | 0.8917 |

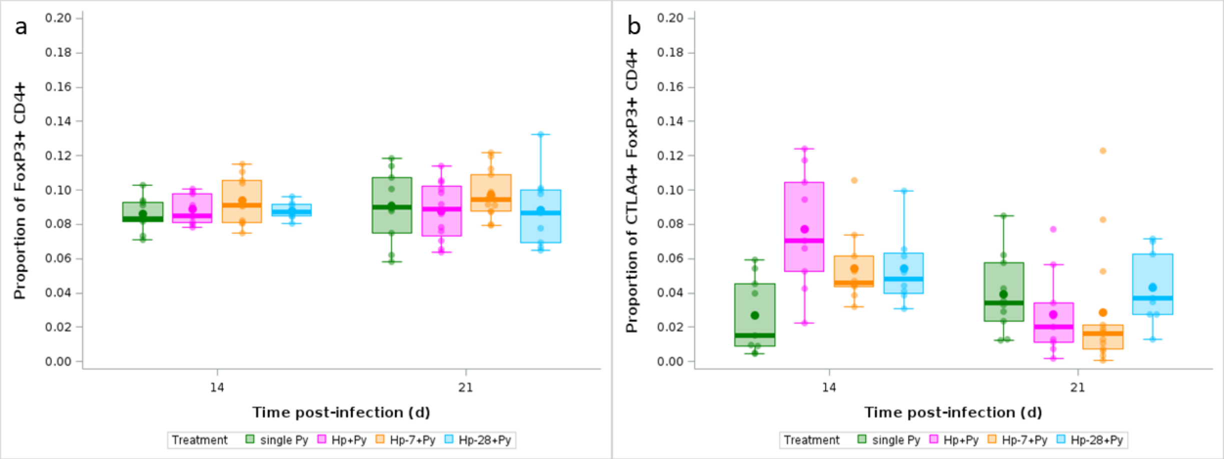


Figure 19. a) FoxP3$^{+}$ (proportion of CD4$^{+}$) T cells analyzed by flow cytometry over the course of the infection in single Py infected and coinfected (Hp+Py, Hp-7+Py, Hp-28+Py) mice; b) CTLA-4$^{+}$ FoxP3$^{+}$ (proportion of CD4$^{+}$) T cells over the course of the infection in single Py infected and coinfected (Hp+Py, Hp-7+Py, Hp-28+Py) mice. Dots represent the raw data, the boxes represent the interquartile range (IQR), the horizontal lines the median, and whiskers the range of data within 1.5 the IQR.

Table S16. Generalized linear model with a beta distribution of errors investigating the changes in the proportion of CTLA-4⁺ FoxP3⁺ CD4⁺ T cells in single Py infected, and coinfected (Hp+Py, Hp-7+Py, Hp-28+Py) mice at day 14 and 21 post-infection. We report the degrees of freedom (df), and the F and p values. N = 77 individuals.

| **Source of variation** | **df** | **F** | **p** |
|---|---|---|---|
| Treatment | 3,69 | 2.44 | 0.0720 |
| Time | 1,69 | 12.16 | 0.0009 |
| Treatment*Time | 3,69 | 6.97 | 0.0004 |

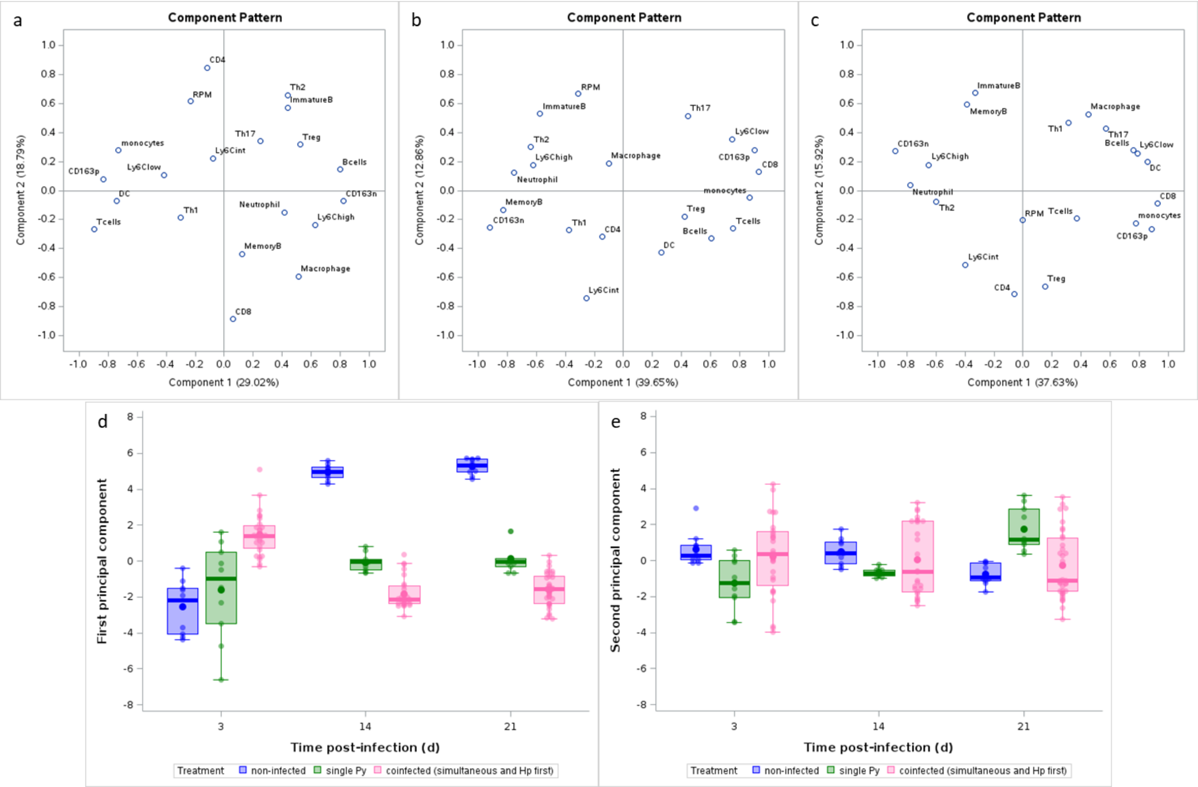


Figure S20. Immunophenotyping of spleen over the course of the infection. Principal component analysis (PCA) on the proportion of 20 immune cell populations at day 3 (a,d), 14 (b,e), 21 (c,f) p.i. in non-infected, single Py infected, and coinfected (simultaneous infection and Hp first) hosts. We report the loadings of the 20 variables on the first two components of the PCA at each sampling day (a,b,c); the scores on the PC1 (d) and PC2 (e) for the three experimental groups. For d) and e), dots represent the raw data, the boxes represent the interquartile range (IQR), the horizontal line the median, and whiskers the range of data within 1.5 the IQR.

Table S17. List of immune cell populations identified by flow cytometry analysis included in the PCA and variable loadings on the two principal components (PC1, PC2) at each sampling day. N= 148.

| | **Loadings** | | | | | |
|---|---|---|---|---|---|---|
| | ***Day 3 p.i.*** | | ***Day 14 p.i.*** | | ***Day 21 p.i.*** | |
| | *PC1* | *PC2* | *PC1* | *PC2* | *PC1* | *PC2* |
| $CD3^{+}$ (T cells) | -0.899 | -0.265 | 0.756 | -0.261 | 0.368 | -0.191 |
| $CD4^{+}$ (helper T cells) | -0.118 | 0.846 | -0.143 | -0.317 | -0.057 | -0.715 |
| $CD8^{+}$ (cytotoxic T cells) | 0.060 | -0.883 | 0.933 | 0.1305 | 0.923 | -0.086 |
| T-$bet^{+}$ $CD4^{+}$ (Th1 cells) | -0.300 | -0.188 | -0.372 | -0.271 | 0.311 | 0.468 |
| ROR$\gamma t^{+}$ $CD4^{+}$ (Th17 cells) | 0.249 | 0.343 | 0.444 | 0.515 | 0.569 | 0.425 |
| GATA-$3^{+}$ $CD4^{+}$ (Th2 cells) | 0.438 | 0.658 | -0.637 | 0.302 | -0.597 | -0.077 |
| $FoxP3^{+}$ $CD4^{+}$ (Treg cells) | 0.525 | 0.320 | 0.421 | -0.182 | 0.150 | -0.659 |
| $CD19^{+}$ (B cells) | 0.798 | 0.144 | 0.604 | -0.331 | 0.759 | 0.275 |
| T-$bet^{+}$ $CD19^{+}$ Memory B cells | 0.125 | -0.438 | -0.830 | -0.137 | -0.388 | 0.591 |
| $CD127^{+}$ $CD19^{+}$ Immature B cells | 0.438 | 0.573 | -0.575 | 0.527 | -0.331 | 0.674 |
| $Ly6G^{+}$ LIN (Neutrophils) | 0.415 | -0.150 | -0.751 | 0.123 | -0.778 | 0.040 |
| $MHCII^{hi}$ $CD11c^{hi}$ (Dendritic cells) | -0.743 | -0.071 | 0.258 | -0.424 | 0.854 | 0.198 |
| F4/$80^{+}$ $CD11b^{+}$ (Macrophages) | 0.514 | -0.594 | -0.100 | 0.184 | 0.447 | 0.525 |
| F4/$80^{+}$ $CD11b^{-}$ (Red pulp macrophages, RPM) | -0.234 | 0.616 | -0.309 | 0.666 | -0.001 | -0.206 |
| $CD163^{+}$ F4/$80^{+}$ $CD11b^{-}$ $MHCII^{+}$ (scavenger RPM) | -0.833 | 0.077 | 0.901 | 0.275 | 0.887 | -0.264 |
| $CD163^{-}$ F4/$80^{+}$ $CD11b^{-}$ $MHCII^{+}$ (other RPM) | 0.823 | -0.071 | -0.919 | -0.253 | -0.880 | 0.270 |
| F4/$80^{-/low}$ $CD11b^{+}$ (Monocytes) | -0.729 | 0.279 | 0.869 | -0.047 | 0.776 | -0.225 |
| $Ly6C^{hi}$ F4/$80^{-/low}$ $CD11b^{+}$ $MHCII^{-/low}$ (Monocytes) | 0.628 | -0.237 | -0.624 | 0.173 | -0.651 | 0.177 |
| $Ly6C^{int}$ F4/$80^{-/low}$ $CD11b^{+}$ $MHCII^{-/low}$ (Monocytes) | -0.077 | 0.222 | -0.255 | -0.741 | -0.400 | -0.511 |
| $Ly6C^{-/low}$ F4/$80^{-/low}$ $CD11b^{+}$ $MHCII^{-/low}$ (Monocytes) | -0.413 | 0.106 | 0.750 | 0.352 | 0.7875 | 0.2577 |
